# Identification of Key Genes and Biological Pathways in Bipolar Disorder by Bioinformatics and Next Generation Sequencing Data Analysis

**DOI:** 10.1101/2022.04.29.489994

**Authors:** Basavaraj Vastrad, Chanabasayya Vastrad

## Abstract

Bipolar disorder (BD), also known as psychiatric disorder, affects millions of people all over the world. The aim of this investigation was to screen and verify hub genes involved in BD as well as to explore potential molecular mechanisms. The next generation sequencing (NGS) dataset GSE124326 was downloaded from the Gene Expression Omnibus (GEO) database, which contained 480 samples, including 240 BD and 240 normal controls. Differentially expressed genes (DEGs) were filtered and subjected to gene ontology (GO) and pathway enrichment analyses. A protein–protein interaction (PPI) network and module analysis were used to identify hub genes. The miRNA-hub gene regulatory network and TF-hub gene regulatory network were also constructed. Receiver operating characteristic curve (ROC) analysis was used to validate hub genes. In this work, 957 DEGs, including 477 up regulated and 480 down regulated, were obtained from NGS data. The GO and pathway enrichment analyses of the DEGs showed that the up regulated genes were enriched in the neutrophil degranulation, immune system, transport, cytoplasm and enzyme regulator activity, and the down regulated genes were enriched in extracellular matrix organization, diseases of metabolism, multicellular organismal process, cell periphery and metal ion binding. Finally, through analyzing the PPI network, miRNA-hub gene regulatory network and TF-hub gene regulatory network, we screened hub genes UBB, UBE2D1, TUBA1A, RPL11, RPS24, NOTCH3, CAV1, CNBD2, CCNA1 and MYH11 by the Cytoscape software. The current investigation illustrates a characteristic NGS data in BD, which might contribute to the interpretation of the progression of BD and provide novel biomarkers and therapeutic targets for BD.

## Introduction

Bipolar disorder (BD) is a psychiatric disorder characterized by recurrent manic or hypomanic and depressive episodes [1]. According to the report of World Health Organization (WHO), BD is the sixth cause of disability-adjusted life years among all psychiatric diseases [2]. The numbers of cases of BD are rising worldwide and it has become an important mental health concern. It is estimated that the incidence of BD was 30% to 69% in Europe and in the United States [3]. The risk factors associated with BD such as obesity [4], anxiety [5], depression [6], cognitive dysfunction [7], pregnancy [8], hypertension [9], cardiovascular diseases [10], diabetes mellitus [11] and genetic factor [12]. A numerous tests are available for screening and detecting BD; all, however, have their disadvantages [13]. Therefore, an in depth understanding of molecular pathogenesis of BD is great significance.

Studies have shown the role of genes and signaling pathway in the pathogenesis of BD. In recent years, several biomarkers were widely used to identify BD patients, such as CACNA1C [14], BDNF (brain derived neurotrophic factor) [15], GSK3 (glycogen synthetase kinase-3) [16], DUSP6 [17] and SYNE1 [18]. Several signaling pathways were involved in pathogenesis of BD such as kynurenine signaling pathway [19], Wnt and GSK3 signaling pathways [20], cAMP--CREB signaling pathways [21] PI3K/AKT/HIF1-a signaling pathway [22], MAP kinase and phosphoinositide signaling pathway [23], and Notch signaling pathway [24] were responsible for advancement of BD. Therefore, we aimed to further explore the pathogenesis of BD and identify specific molecular targets.

Next generation sequencing (NGS) [25] and bioinformatics analysis [26] are widely used in the investigation of the molecular mechanism of various diseases. NGS data are usually deposited and available in free public websites, such as the NCBI-Gene Expression Omnibus database (NCBI-GEO) (https://www.ncbi.nlm.nih.gov/geo) [27]. Integrated bioinformatics analyses of NGS data derived from investigation of BD could help identify the hub genes and further demonstrate their related functions and potential therapeutic targets in BD.

In the current investigation, NGS data of GSE124326 [28] from NCBI-GEO was downloaded. A total of 240 BD samples and 240 normal control samples were available. Differentially expressed genes (DEGs) between BD and normal control samples were filtered and obtained using the online tool DESeq2. Gene Ontology (GO) and REACTOME pathway enrichment analyses of the DEGs were performed using the g:Profiler. The functions of the DEGs were further assessed by PPI network and modular analyses to identify the hub genes in BD. Moreover, miRNA-hub gene regulatory network and TF-hub gene regulatory network of the hub genes were established, miRNAs and TFs were selected. The diagnostic roles of hub genes were analyzed using receiver operating characteristic curve (ROC) analysis. These add a basis for enhanced understanding of molecular basis of BD pathology and effective pharmaceutical targets.

## Materials and methods

### Data resources

The NGS dataset GSE124326 [28] based on GPL16791 Illumina HiSeq 2500 (Homo sapiens) was acquired from the Gene Expression Ominibus (GEO) database. The GSE124326 dataset contained 240 BD samples and 240 normal control samples.

### Identification of DEGs

DESeq2 package of R software [29] was used to analyze the DEGs between PD and normal control in the NGS data of GSE124326. The adjusted P-value and [log FC] were calculated. The Benjamini & Hochberg false discovery rate method was used as a correction factor for the adjusted P-value in DESeq2 [30]. The statistically significant DEGs were identified according to adjusted P-value < 0.05, [logFC] > 0.235 for up regulated genes and [logFC] < −0.52 for down regulated genes. The DEGs are presented as volcano plot and heat map generated using ggplot2 and gplot in R Bioconductor.

### GO and pathway enrichment analyses of DEGs

g:Profiler (http://biit.cs.ut.ee/gprofiler/) [31] was used to perform GO functional and REACTOME pathway enrichment analyses. GO (http://www.geneontology.org) [32] annotation was applied to define gene functions in three terms: biological process (BP), cellular component (CC) and molecular function (MF). REACTOME (https://reactome.org/) [33] is a pathway database resource for understanding high-level biological functions and utilities. P < 0.05 was considered as statistically significant.

### Construction of the protein-protein interaction (PPI) and module analysis

In order to obtain interacting proteins related to DEGs, the STRING version 11.5 database (https://string-db.org/) was used [34]. Cytoscape (http://www.cytoscape.org/) (version 3.9.1) [35] was used to visualize the PPI network of DEGs. The Network Analyzer plug-in was used to explore hub genes, and the hub genrs were generated using node degree [36], betweenness [37], stress [38] and closeness [39] methods. The intersect function was used to identify the hub genes. The PEWCC1 [40] was used to search modules of the PPI network and the default parameters (Degree cutoff ≥10, node score cutoff ≥0.4, K-core ≥4, and max depth=100.) were set in the functional interface of Cytoscape.

### miRNA-hub gene regulatory network construction

MiRNA regulates gene expression under defined disease conditions through interaction with hub genes during the post transcriptional stage was analyzed. We applied miRNet database (https://www.mirnet.ca/) [41] to integrate miRNA databases (TarBase, miRTarBase, miRecords, miRanda (S mansoni only), miR2Disease, HMDD, PhenomiR, SM2miR, PharmacomiR, EpimiR, starBase, TransmiR, ADmiRE, and TAM 2.0.). We visualized miRNA-hub gene regulatory network by employing Cytoscape software [35].

### TF-hub gene regulatory network construction

TF regulates gene expression under defined disease conditions through interaction with hub genes during the post transcriptional stage was analyzed. We applied NetworkAnalyst database (https://www.networkanalyst.ca/) [42] to integrate TF database (ENCODE). We visualized TF-hub gene regulatory network by employing Cytoscape software [35].

### Receiver operating characteristic curve (ROC) analysis

The hub genes were used to identify biomarkers with high sensitivity and specificity for BD diagnosis. The ROC curves were plotted and area under curve (AUC) was calculated separately to evaluate the performance of each model using the R packages “pROC” [43]. A AUC > 0.8 indicated that the model had a good fitting effect.

## Results

### Identification of DEGs

GSE124326 was selected and underwent DEGs analysis using “DESeq2” package in R software. There was a total of 957 DEGs between BD and normal control samples, including 477 up regulated DEGs and 480 down regulated genes (Table 1). A volcano plot was constructed for the DEGs and is presented in Fig. 1. The DEGs are presented by a heat map in Fig. 2.

**Fig. 1.**
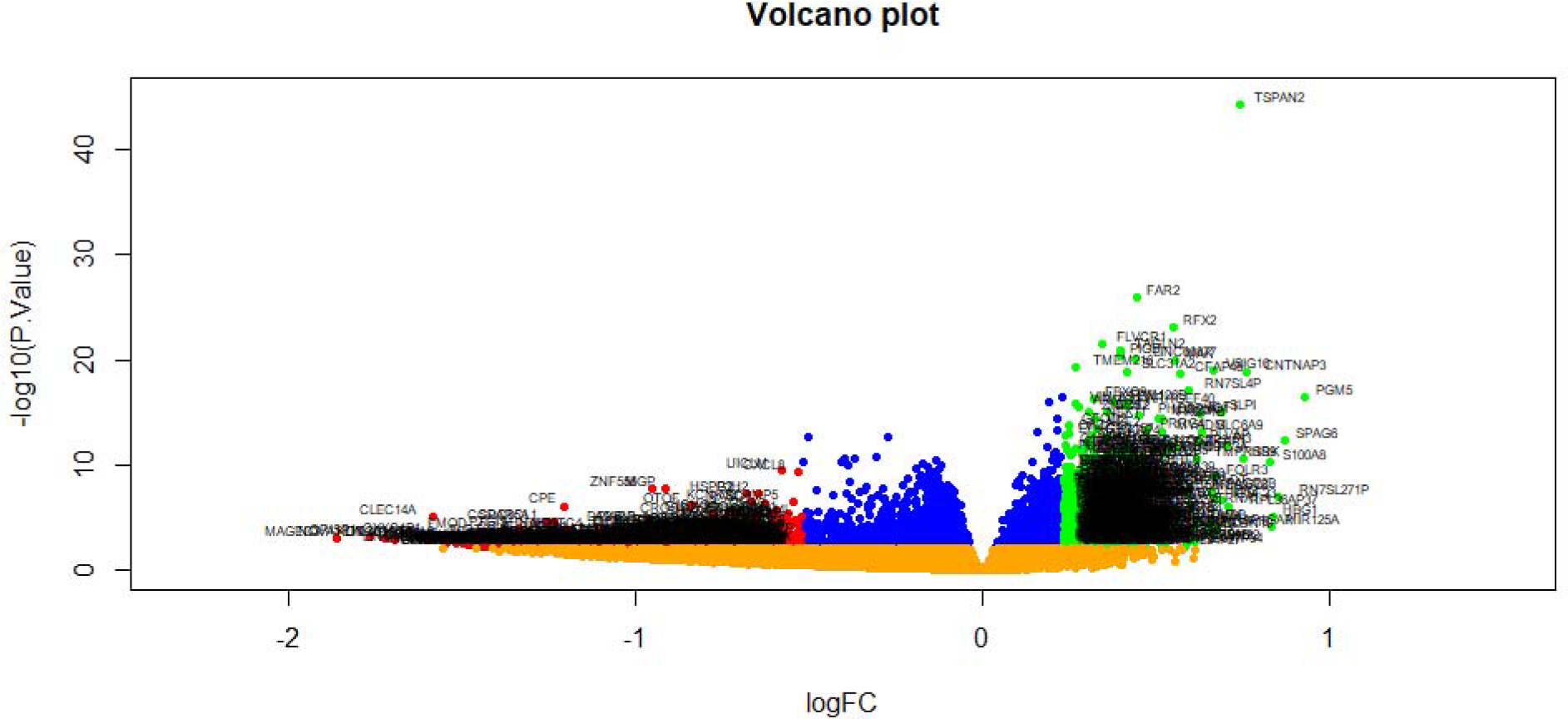
Volcano plot of differentially expressed genes. Genes with a significant change of more than two-fold were selected. Green dot represented up regulated significant genes and red dot represented down regulated significant genes.

**Fig. 2.**
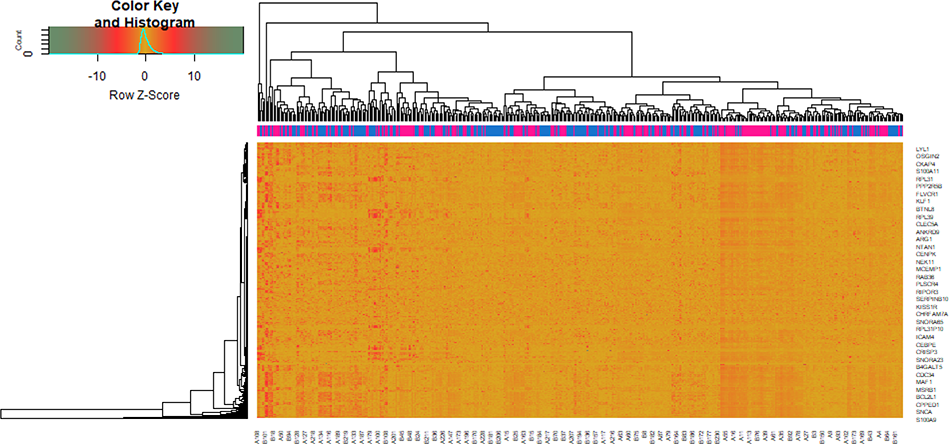
Heat map of differentially expressed genes. Legend on the top left indicate log fold change of genes. (A1 – A240 = normal control samples; B1 – B240 = BD samples)

**Table 1.**
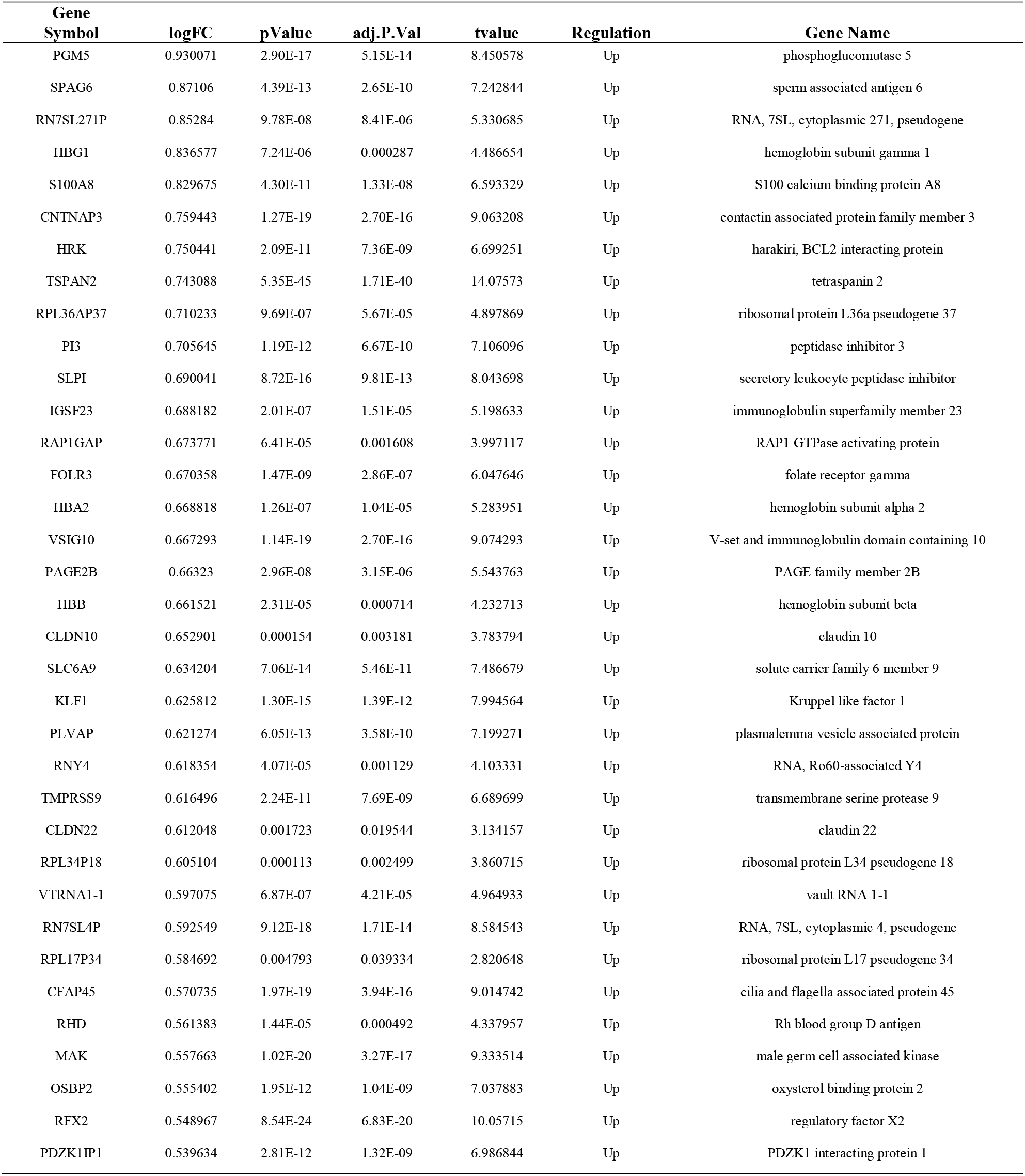
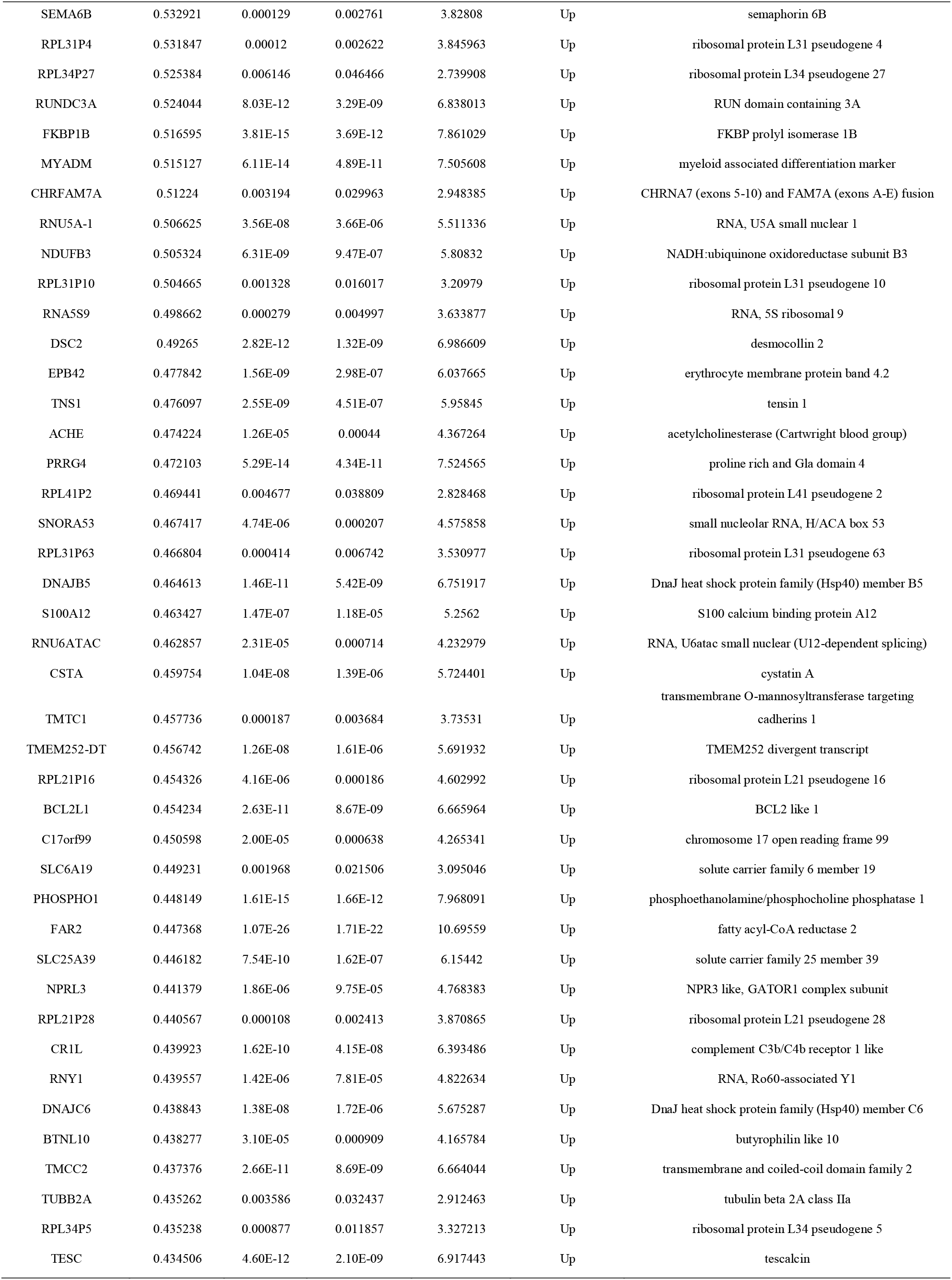
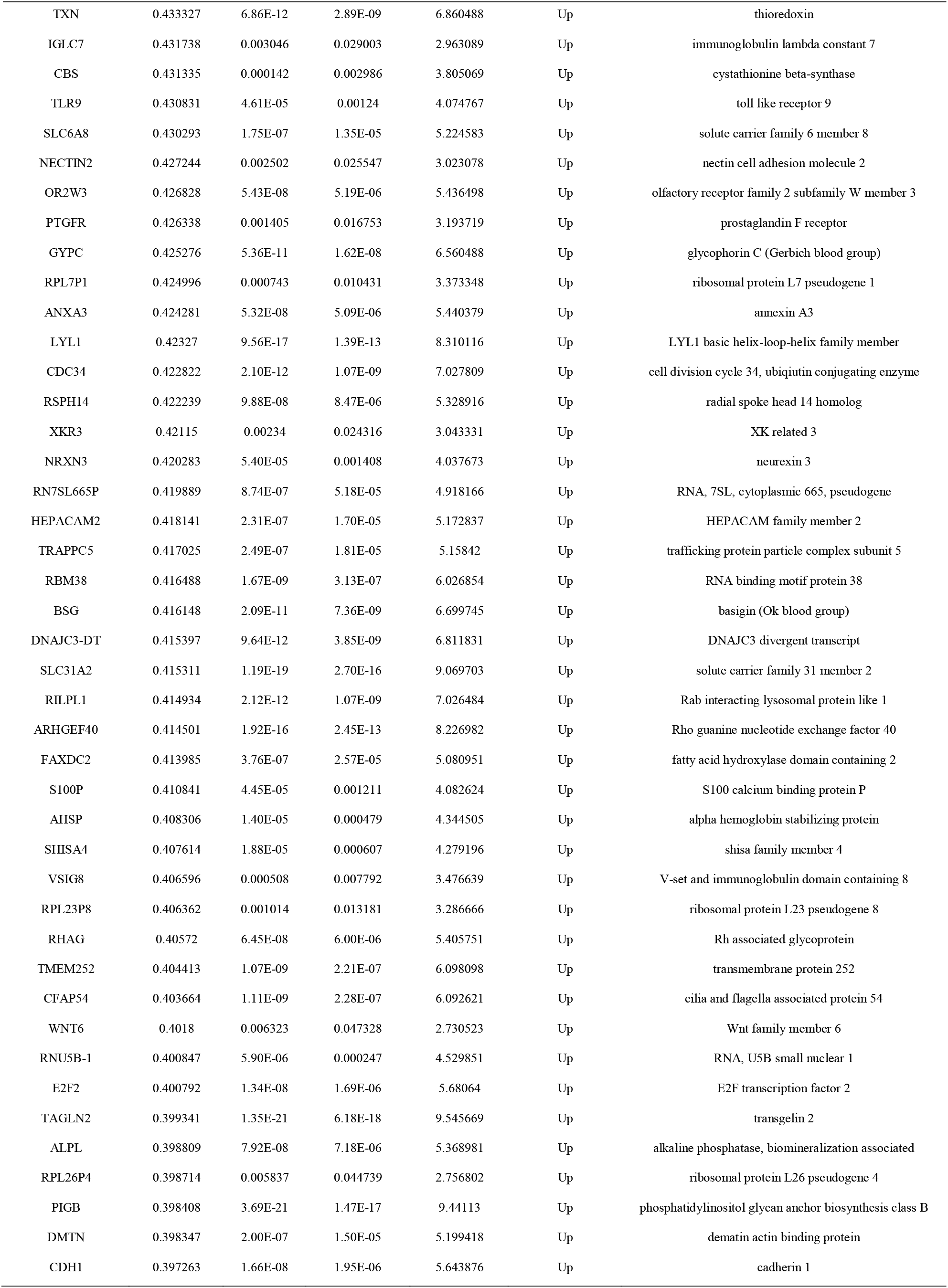
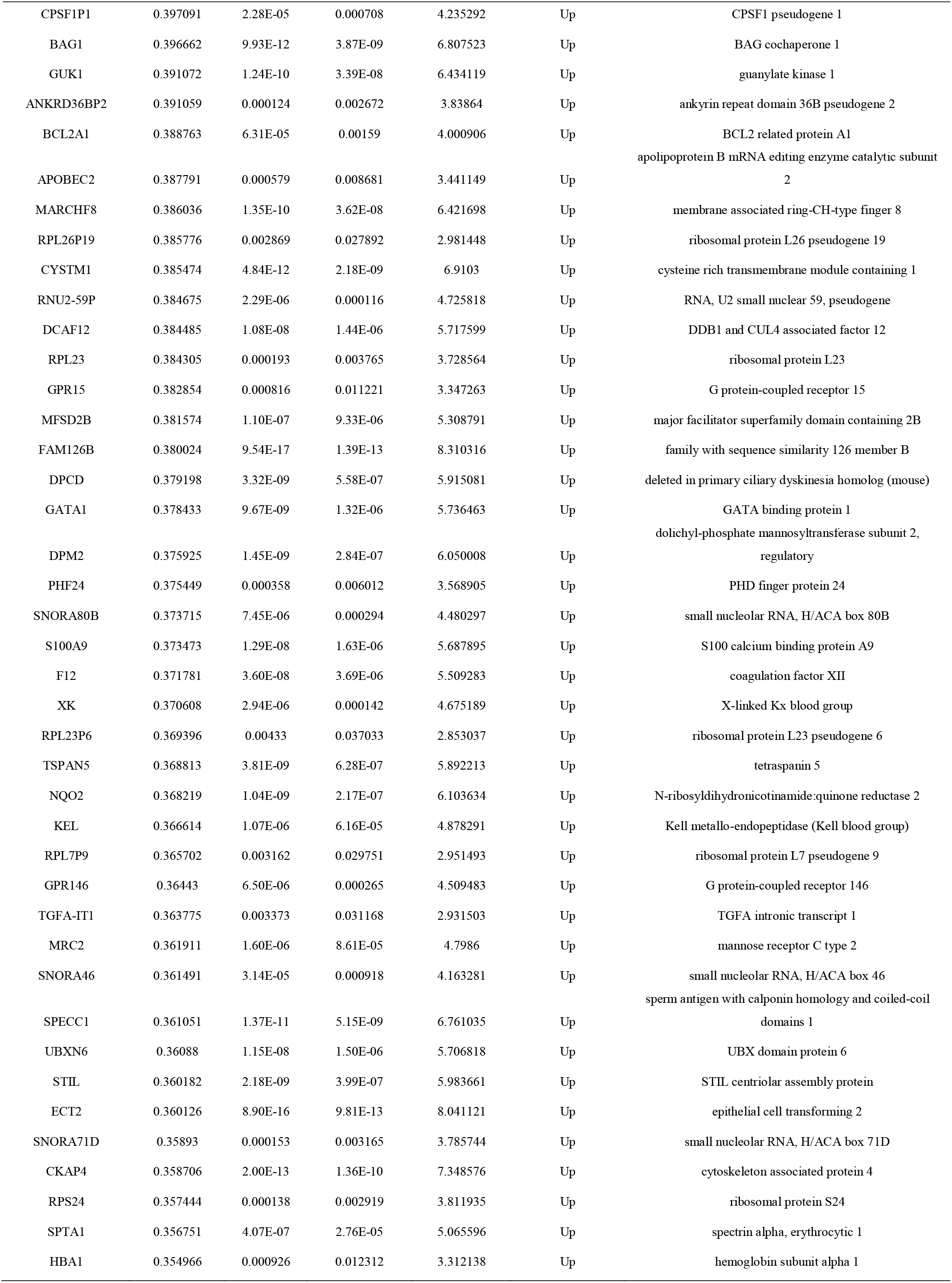
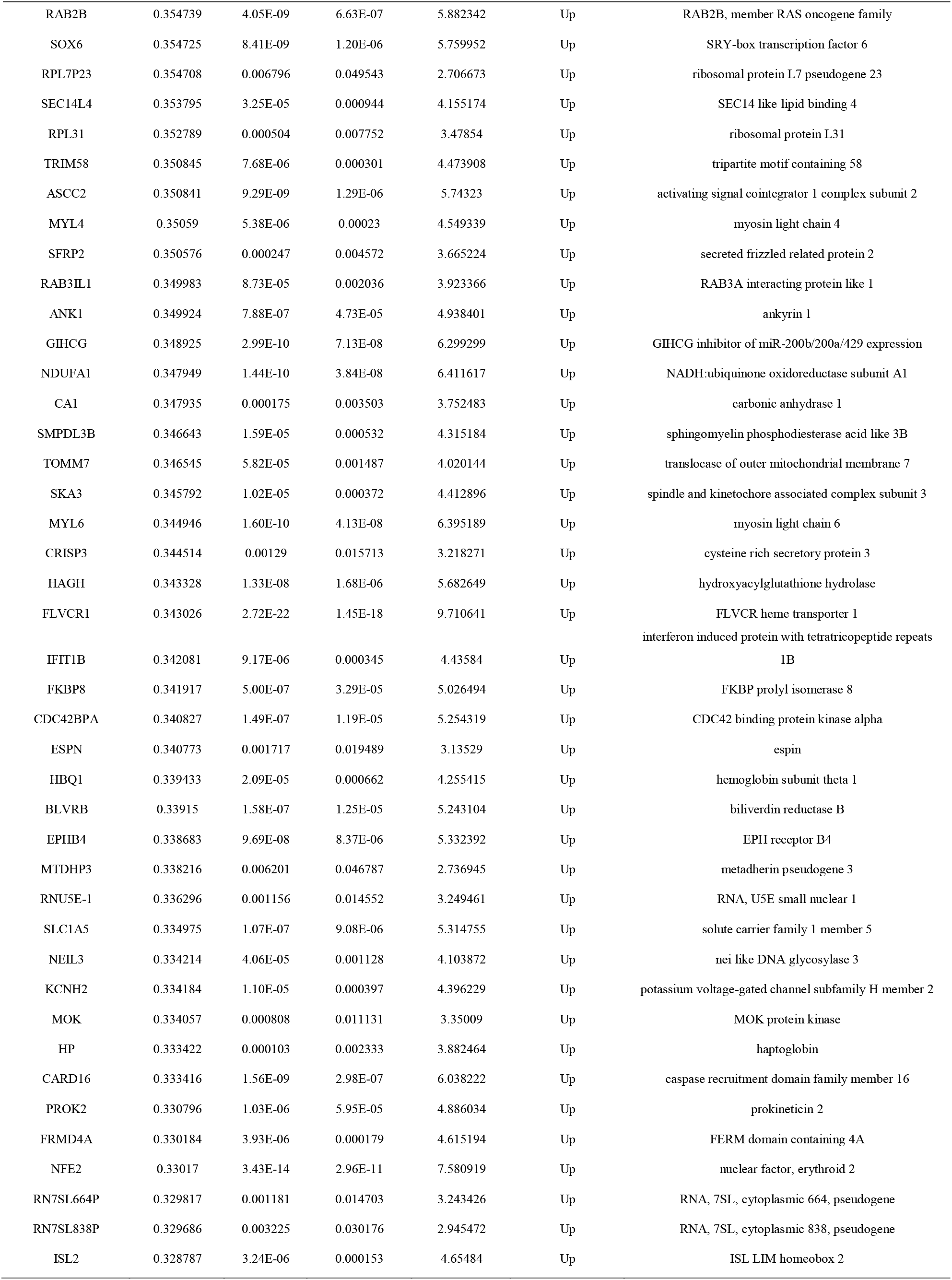
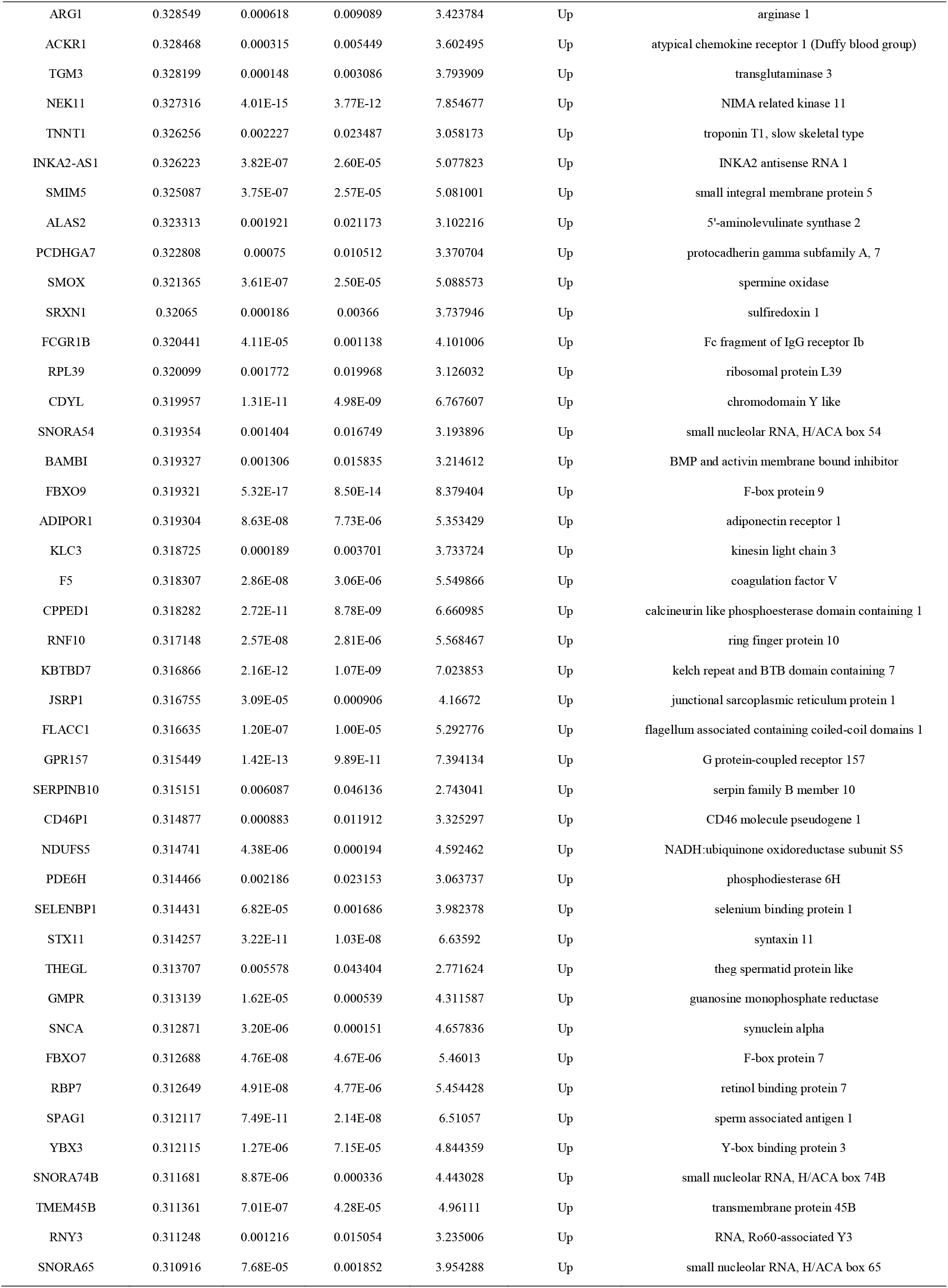
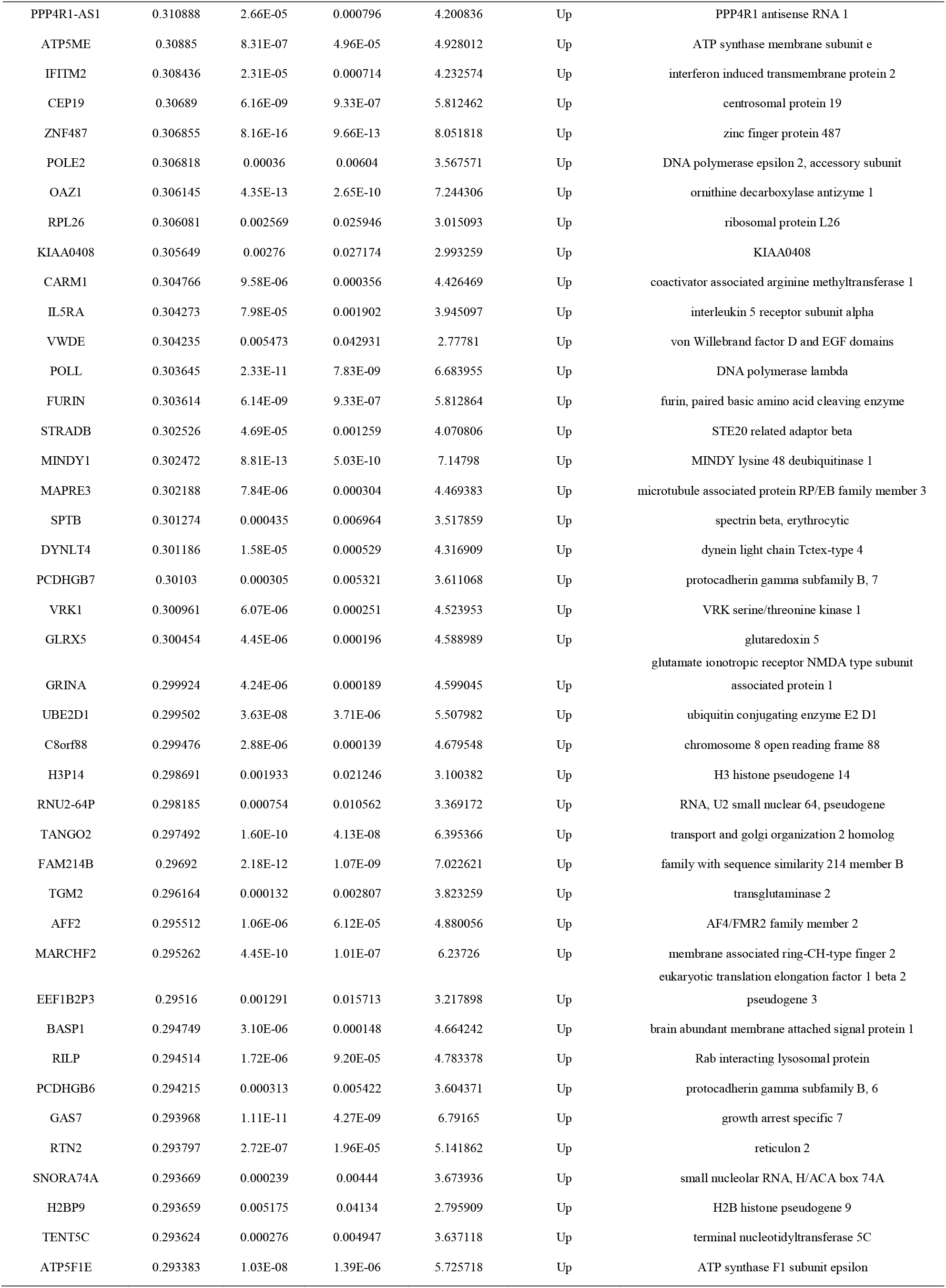
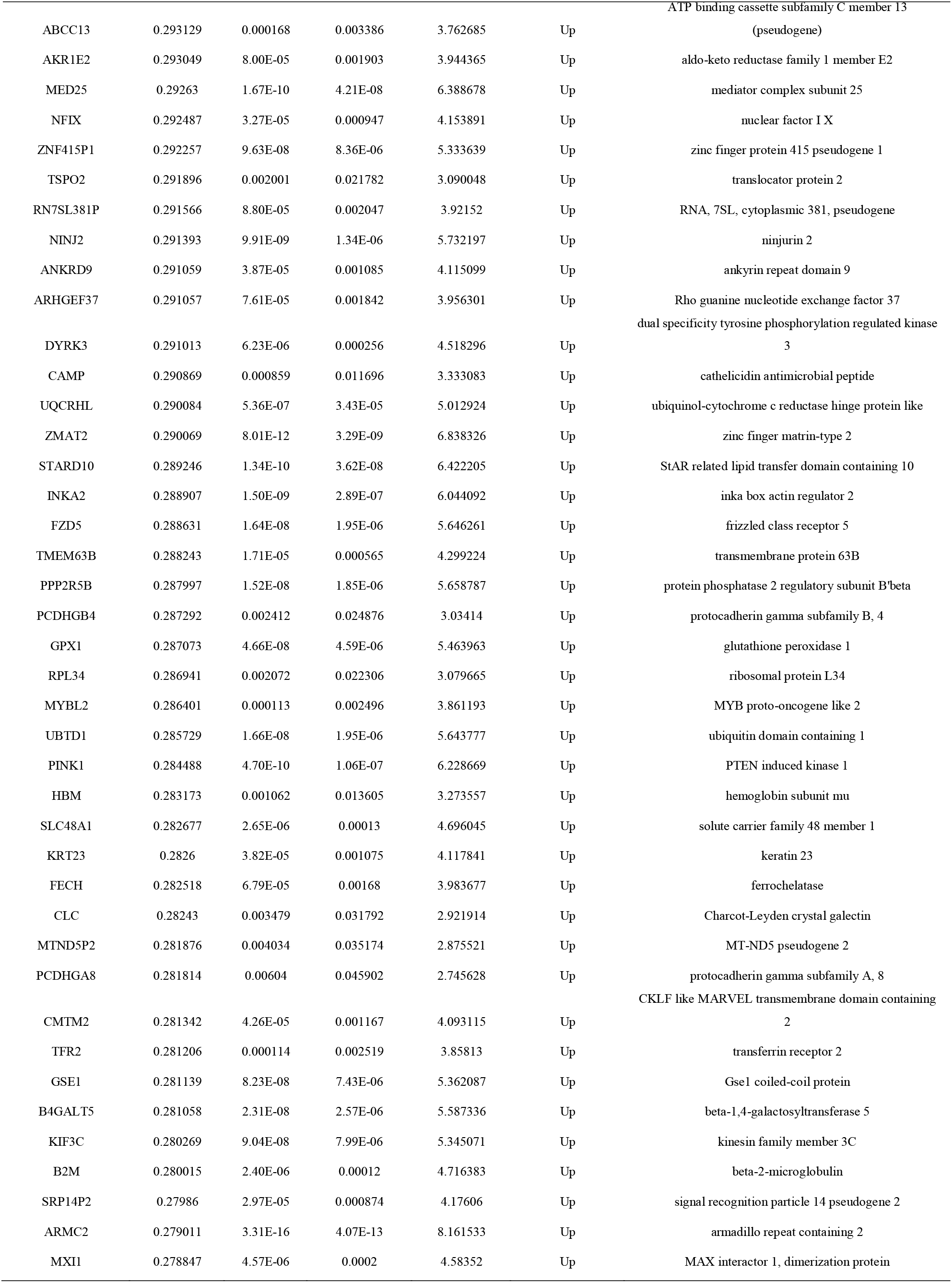
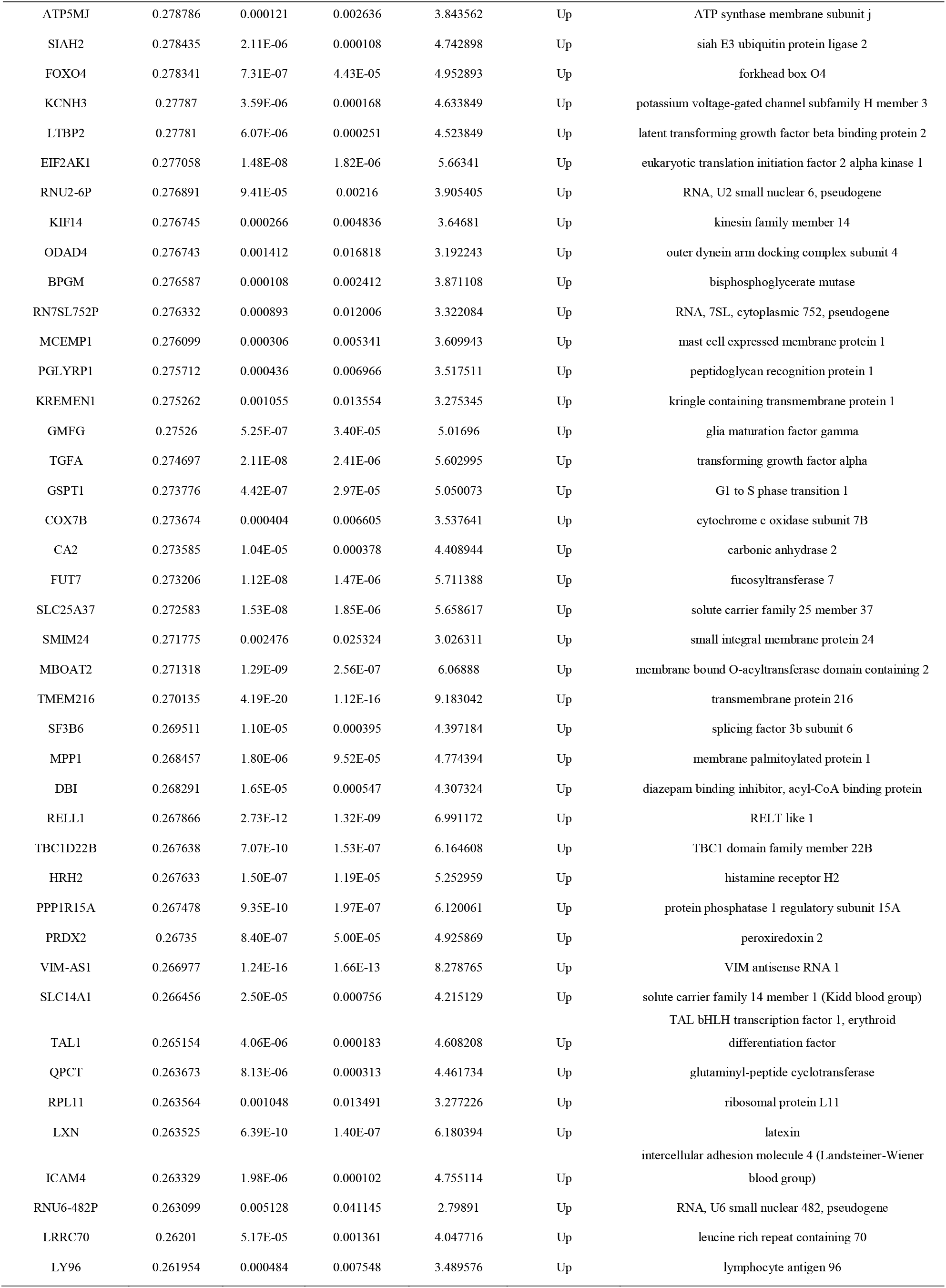
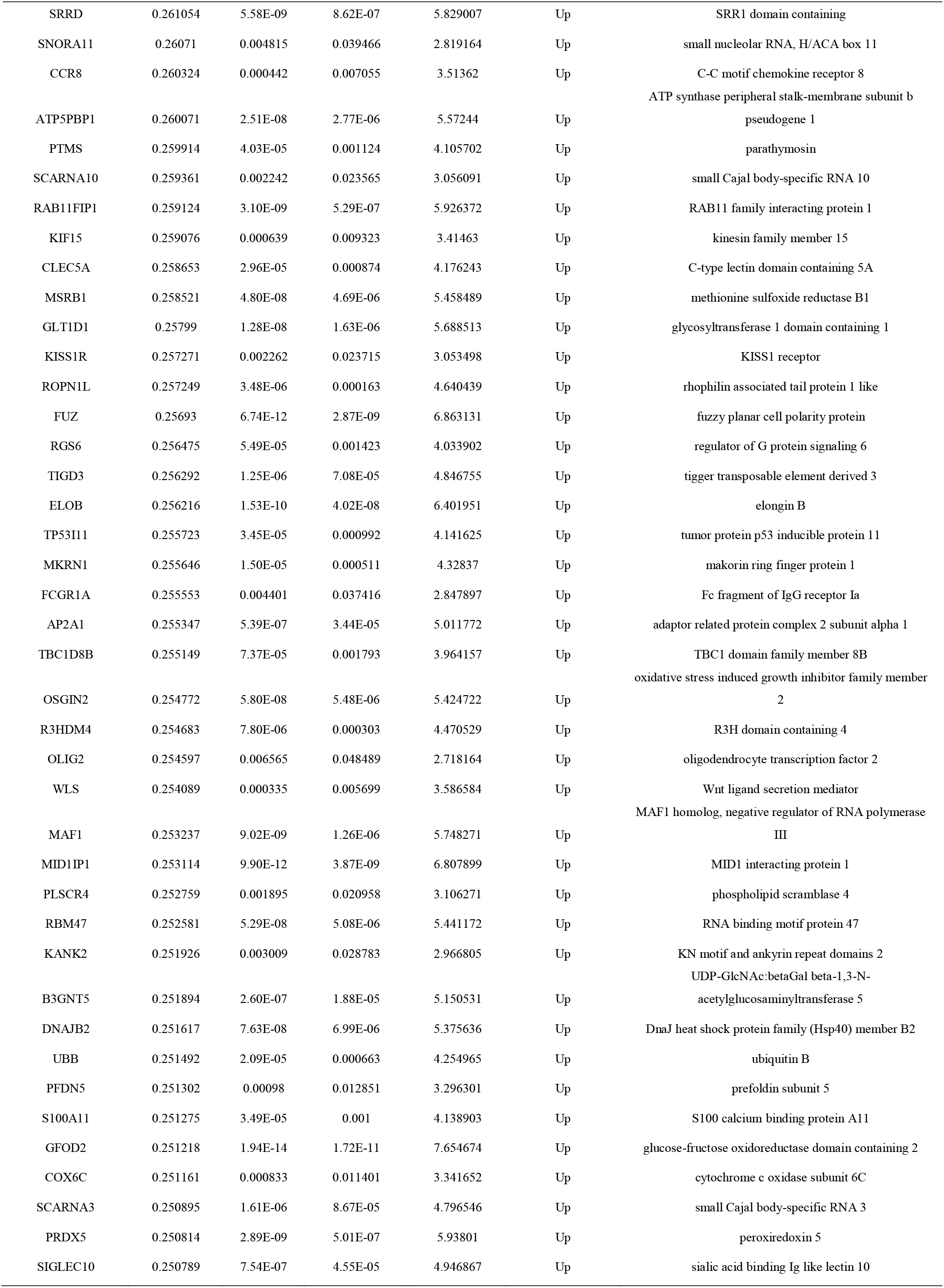
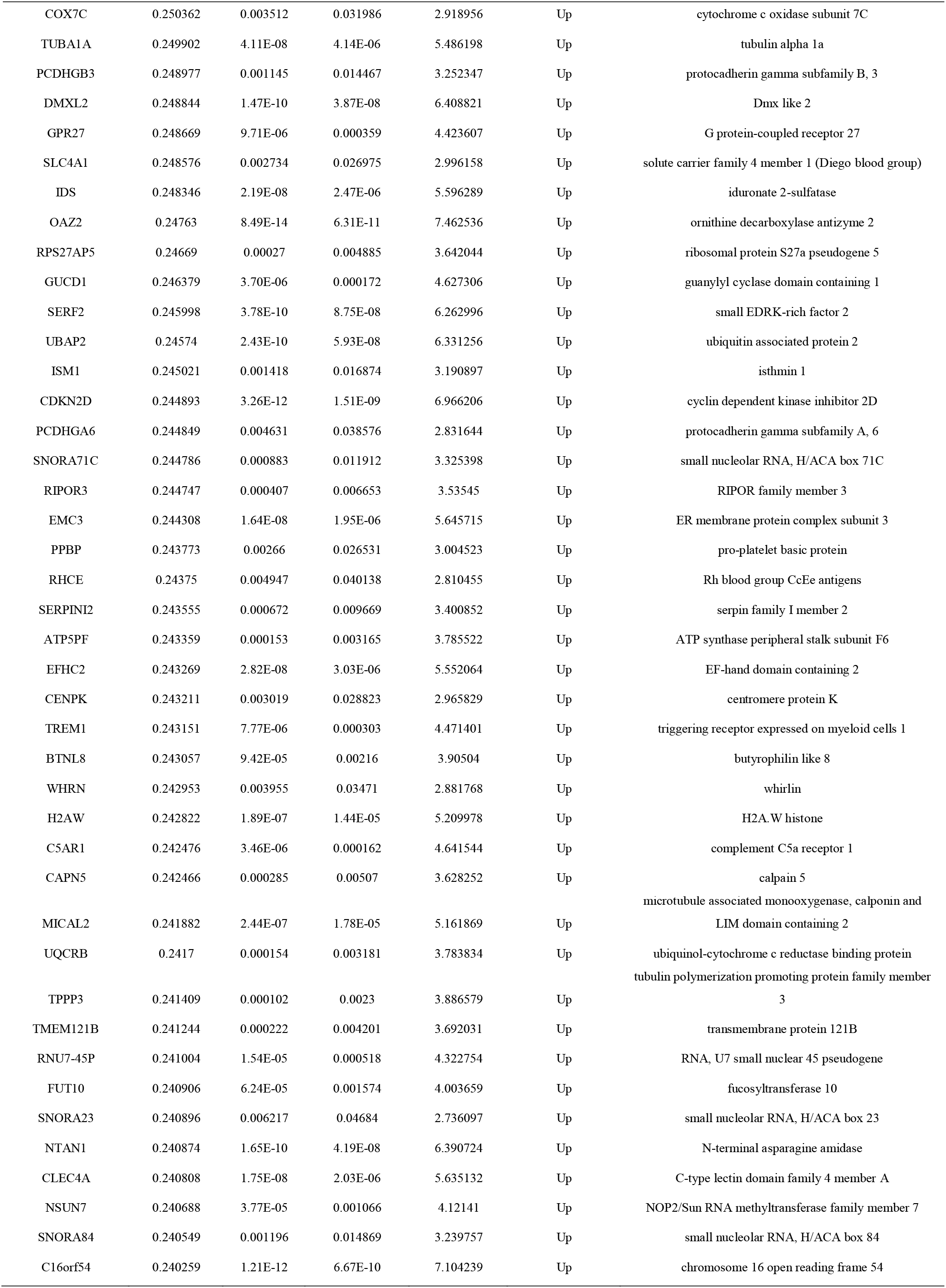
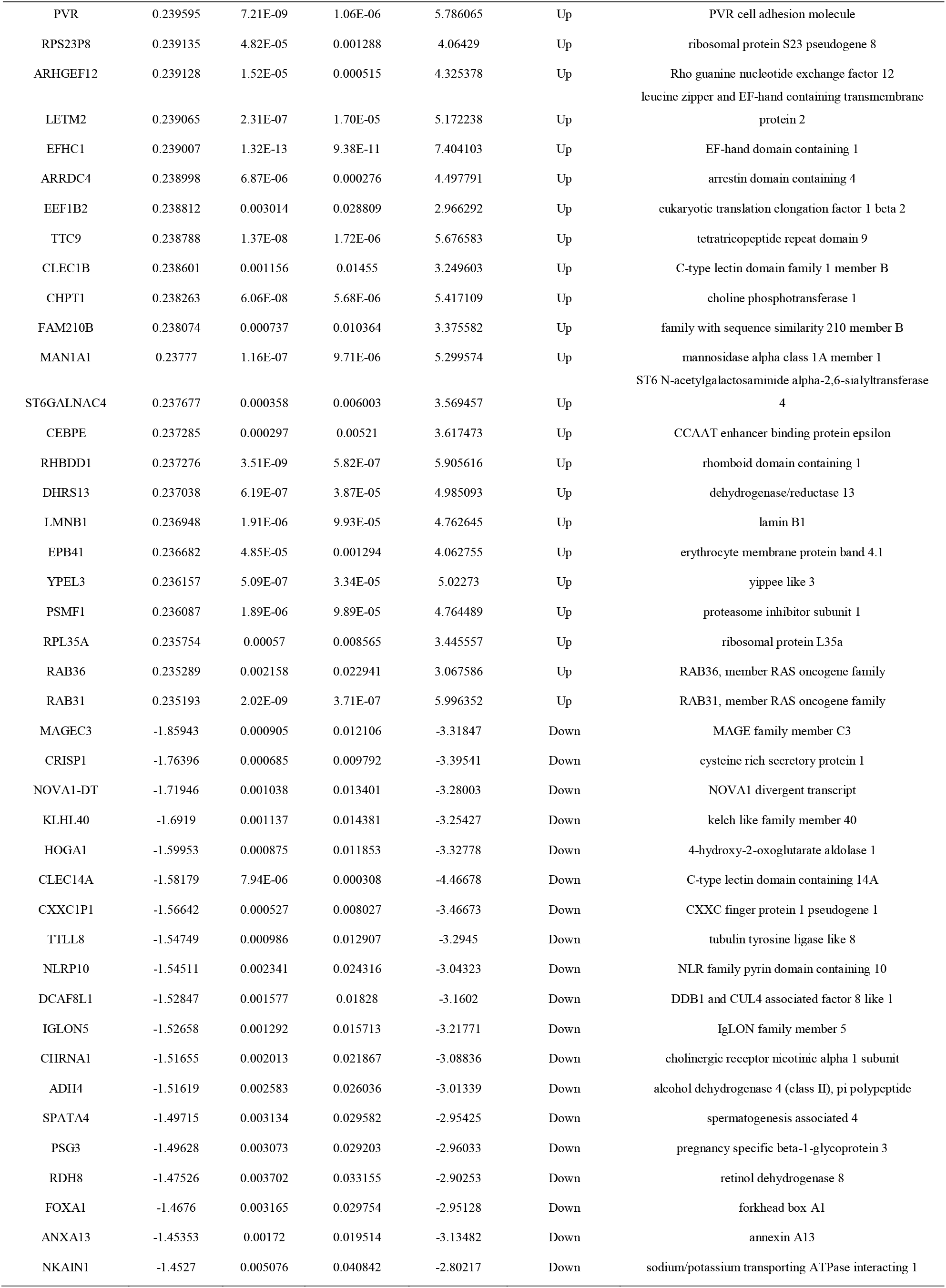
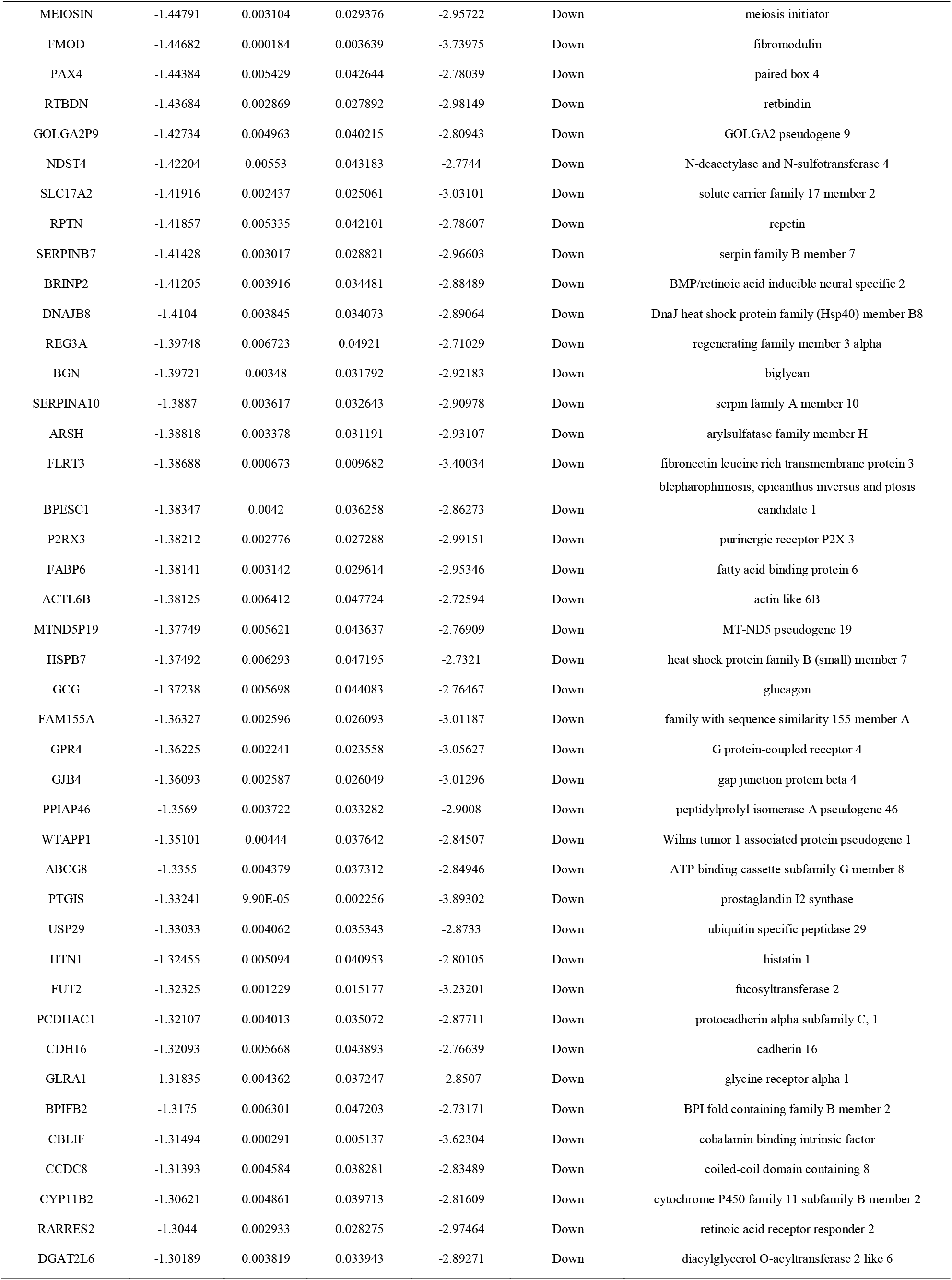
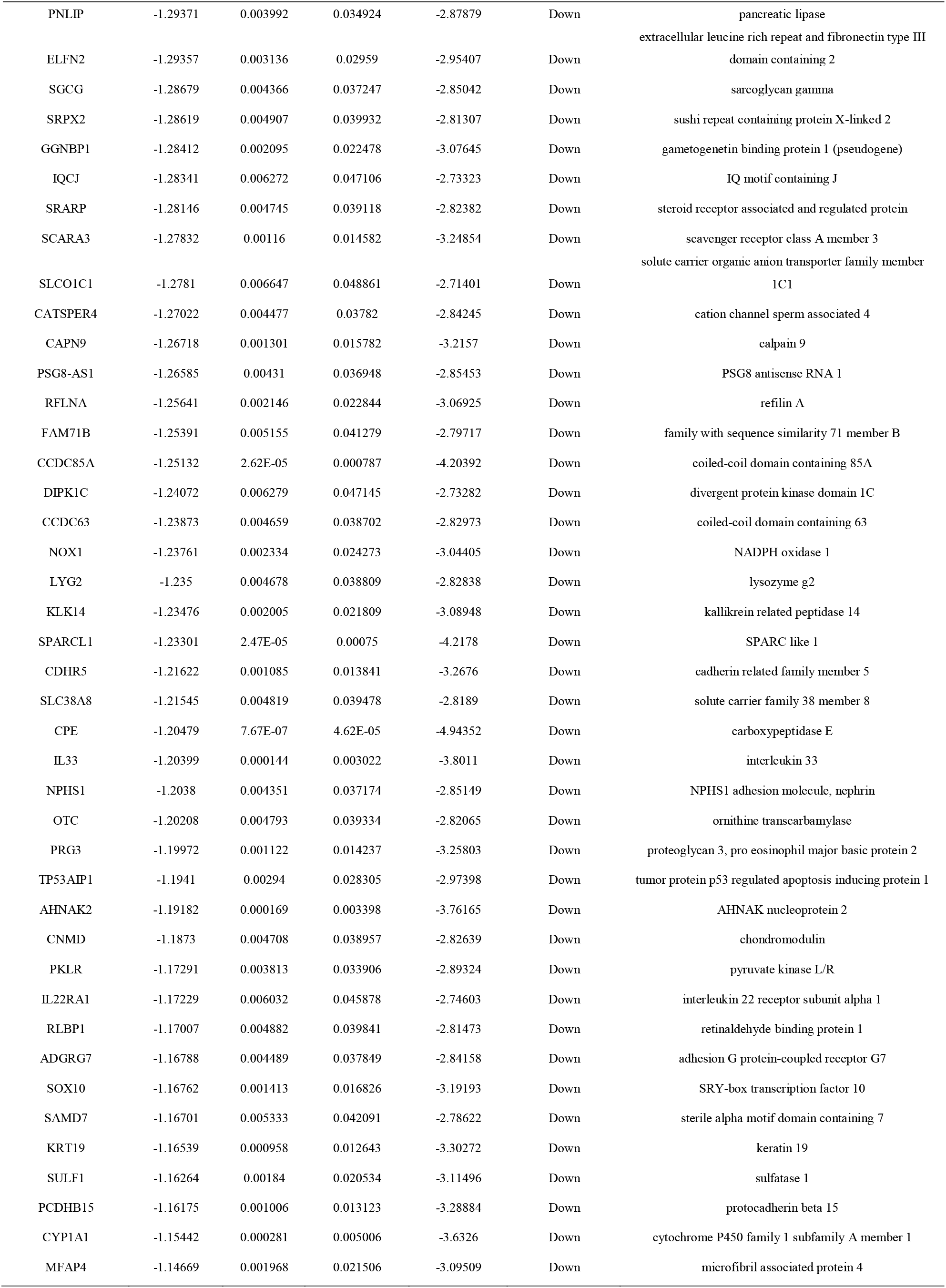
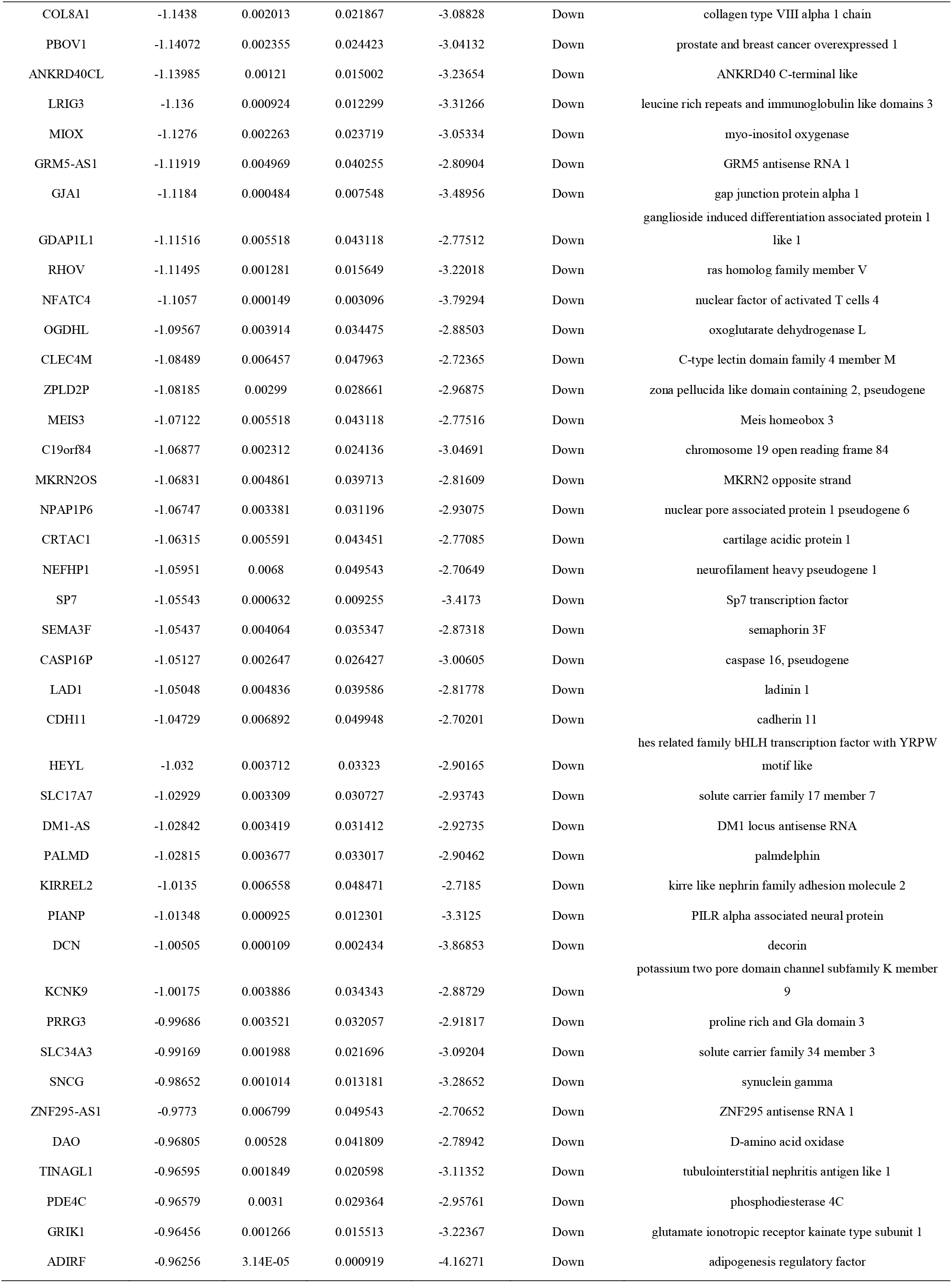
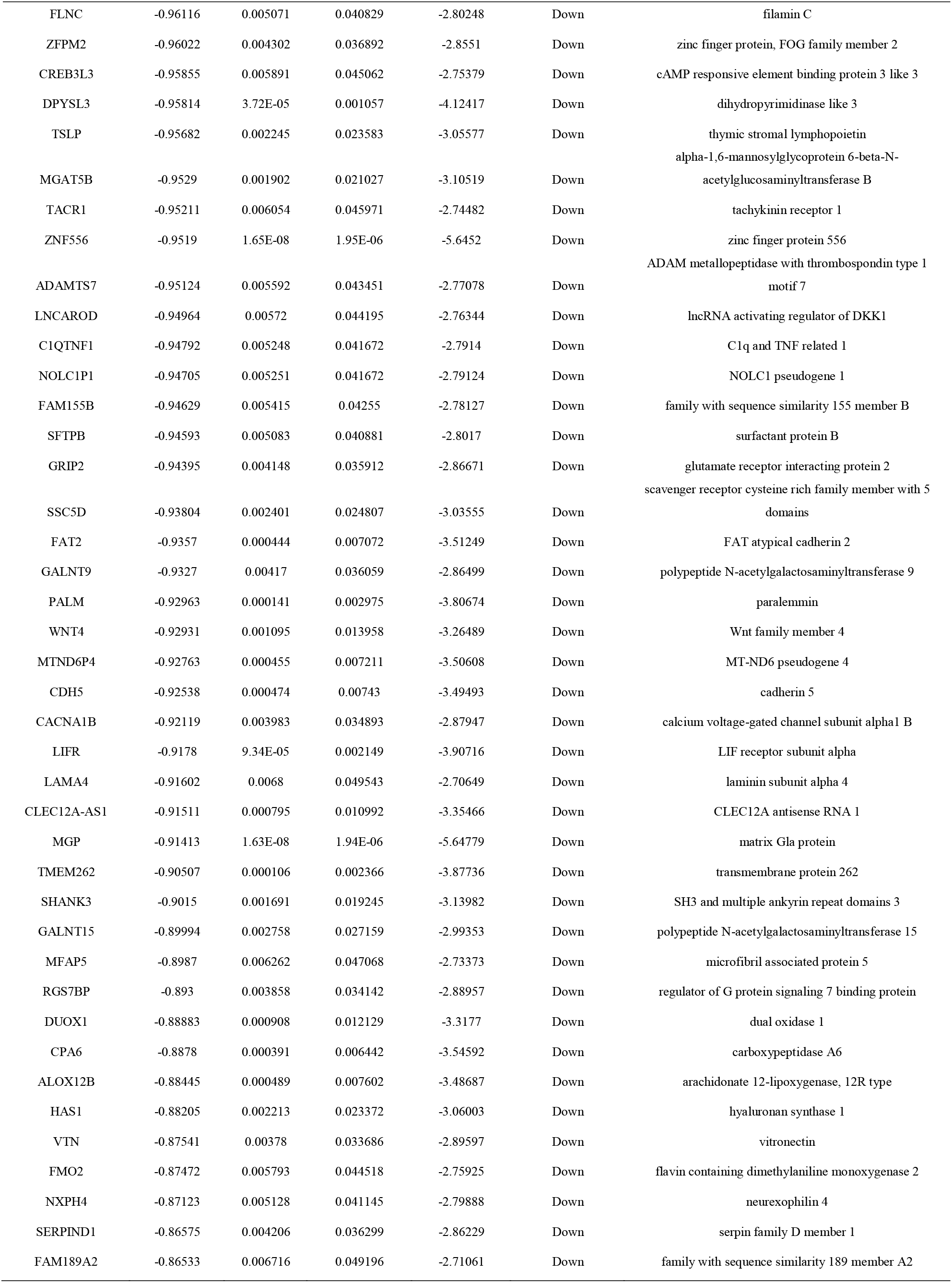
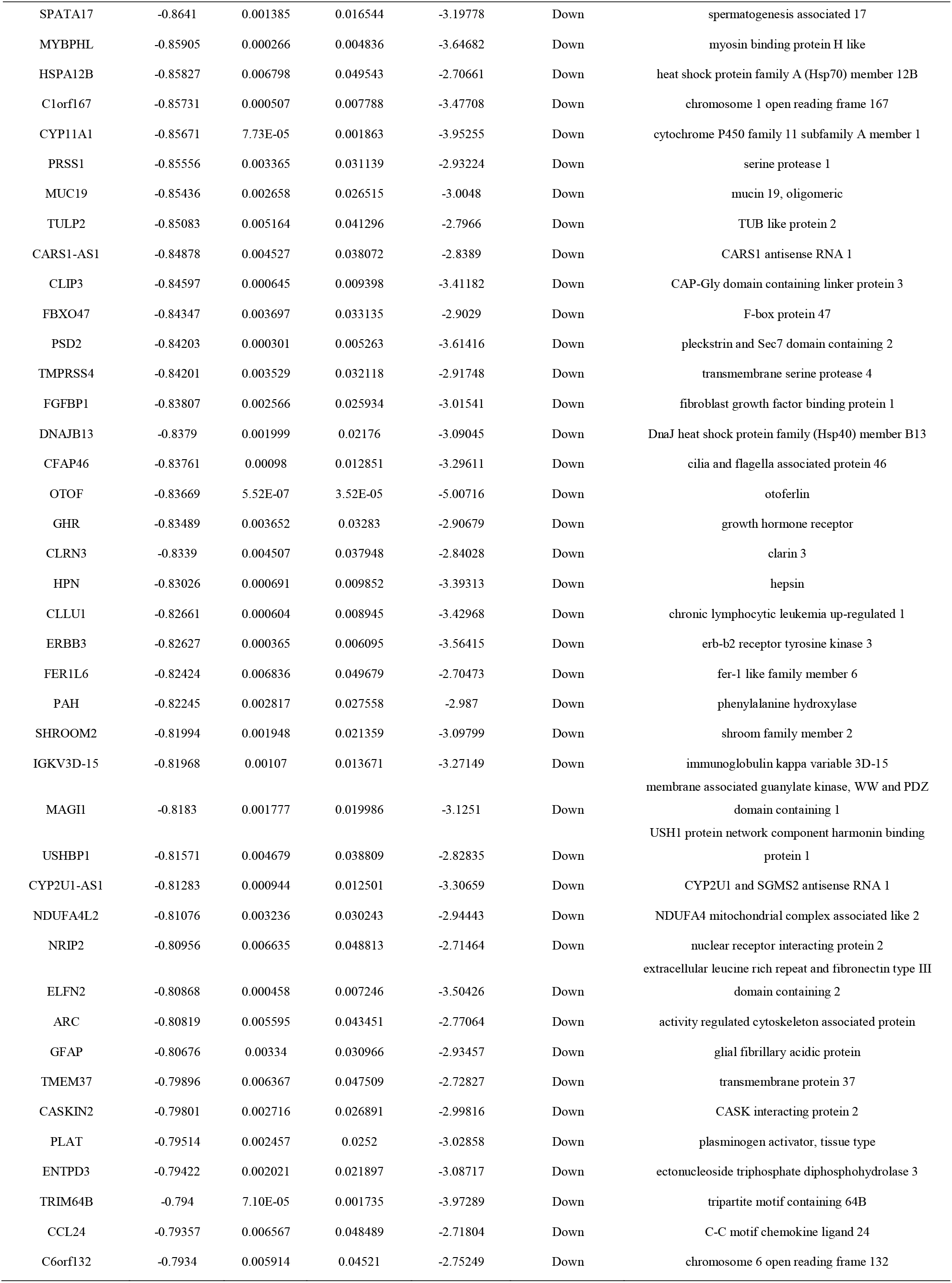
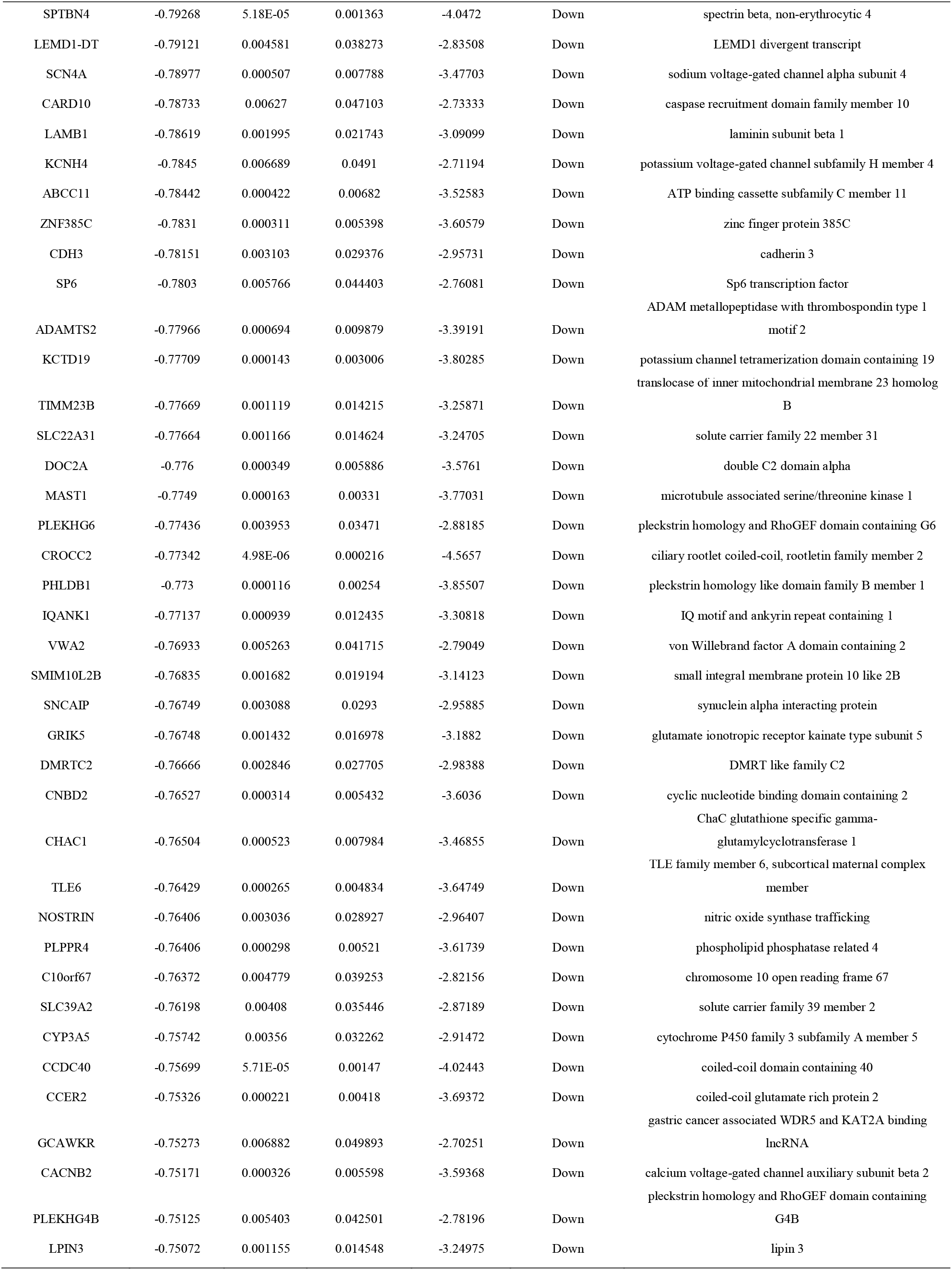
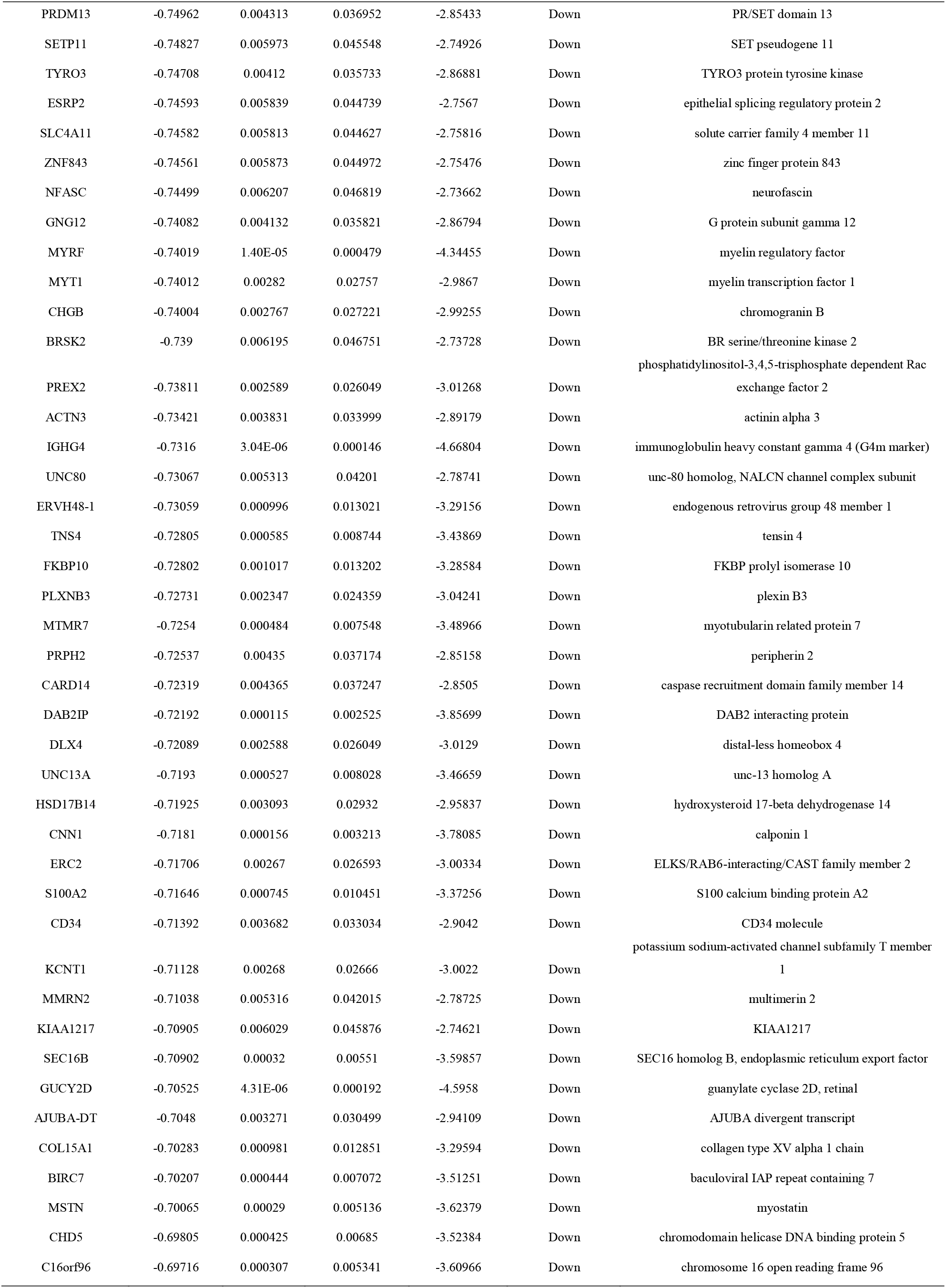
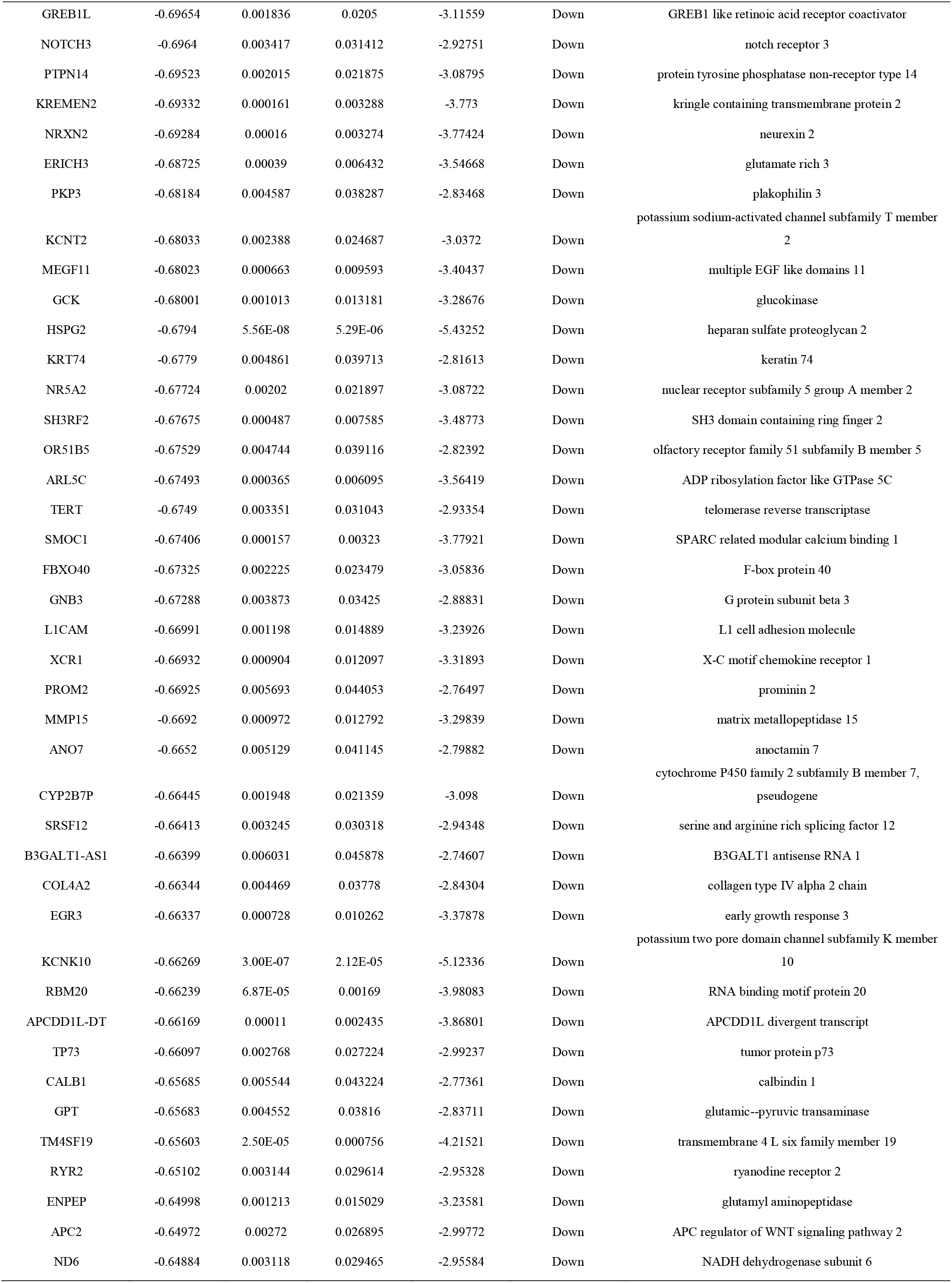
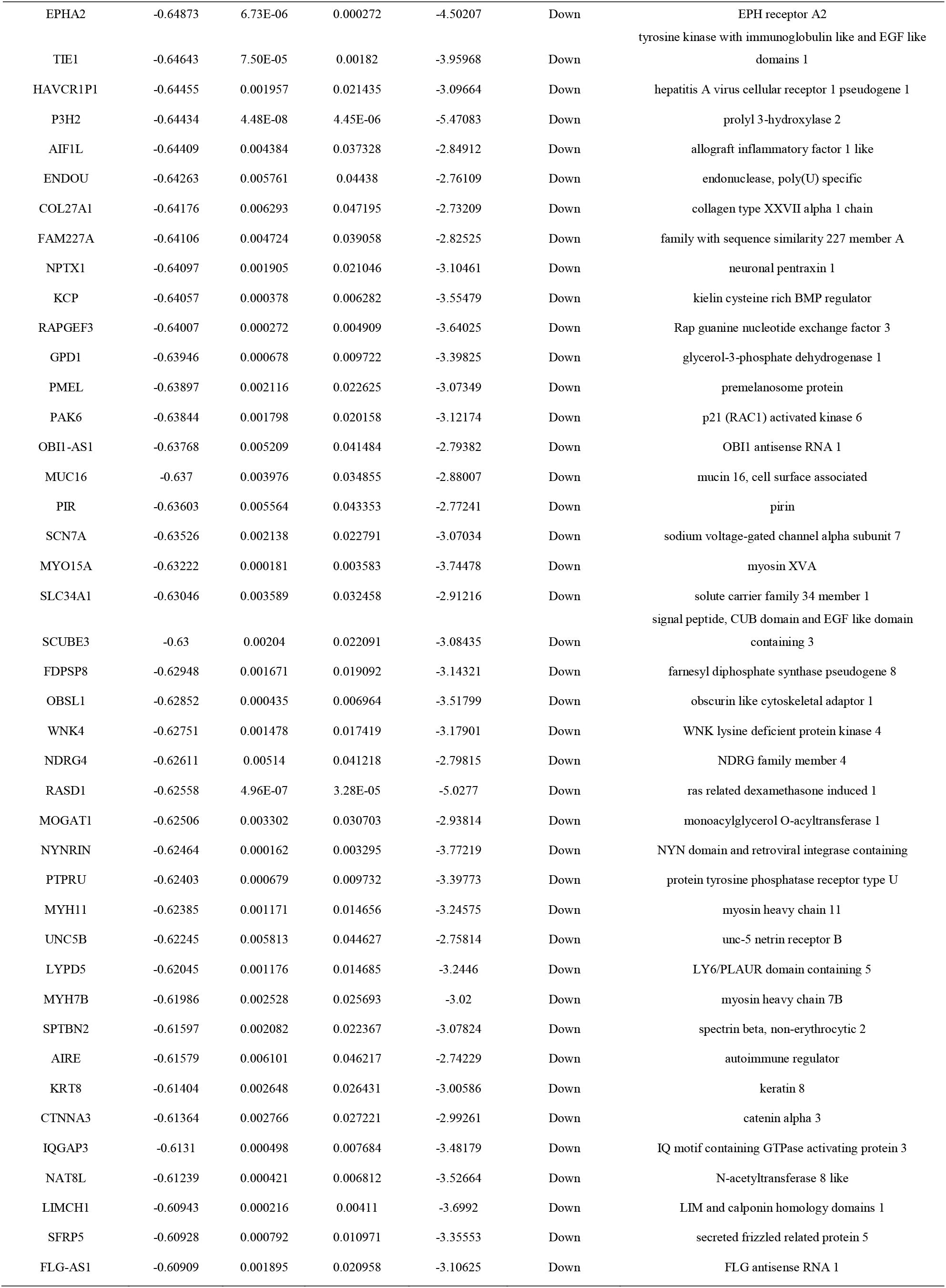
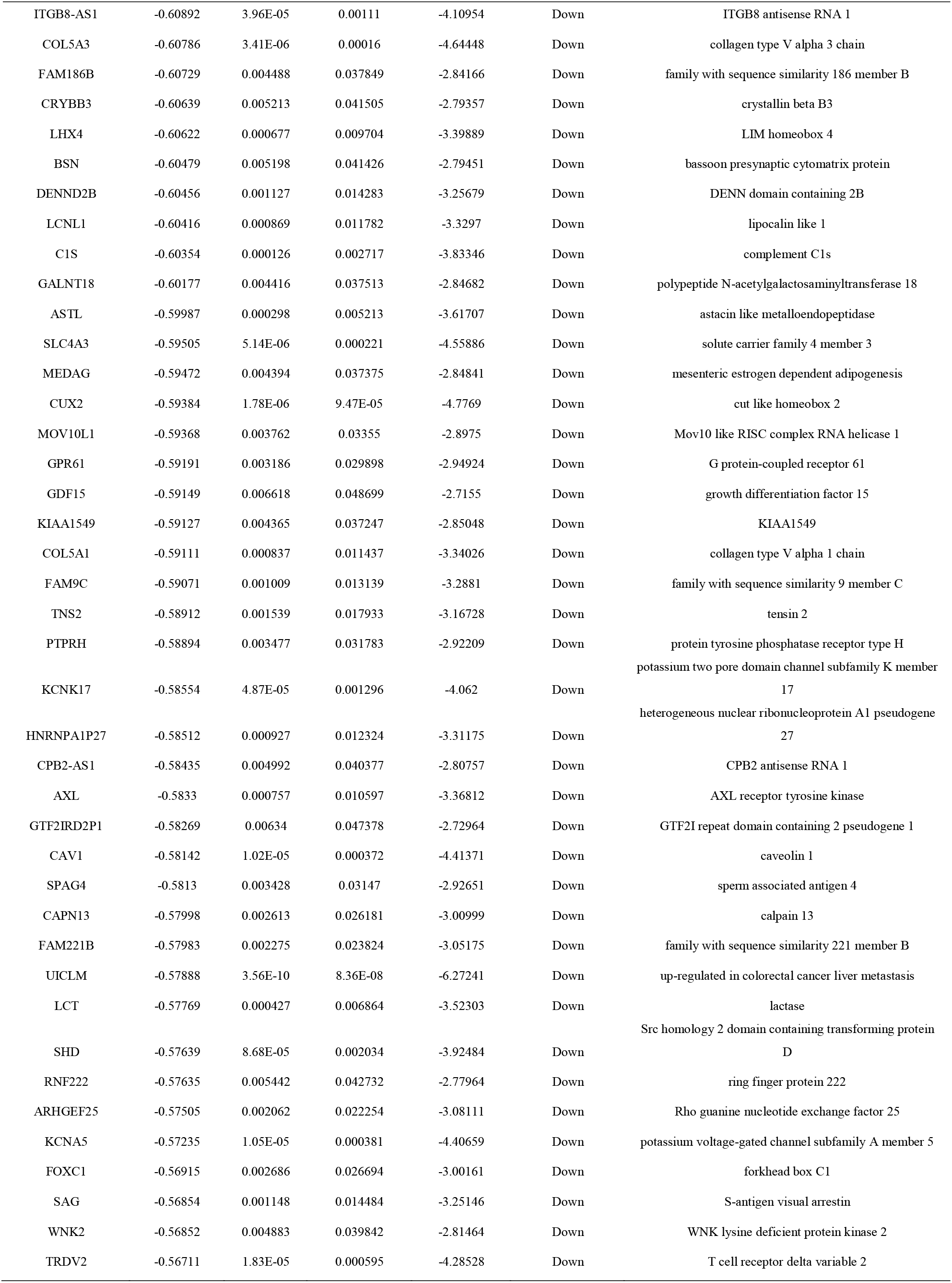
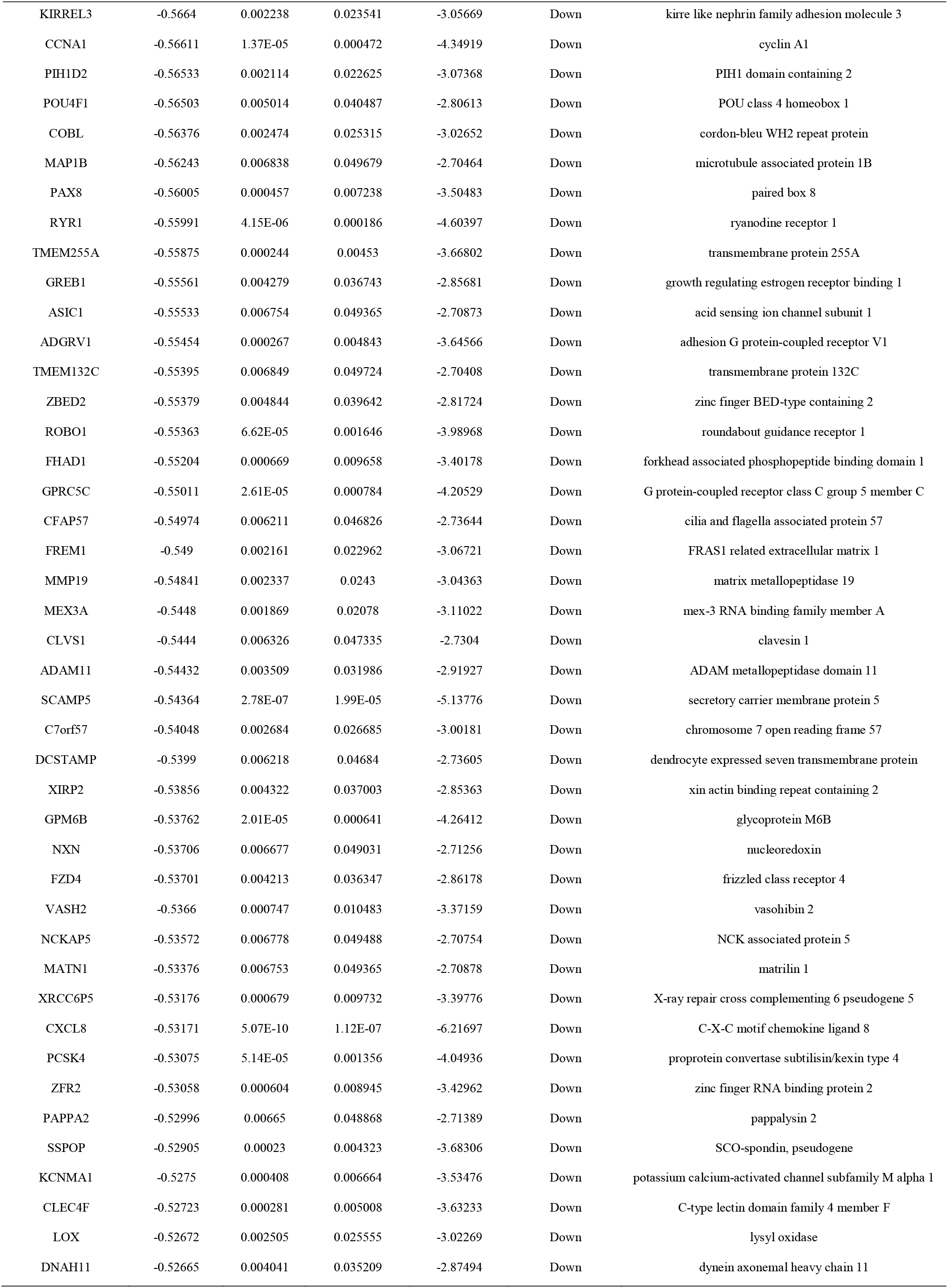
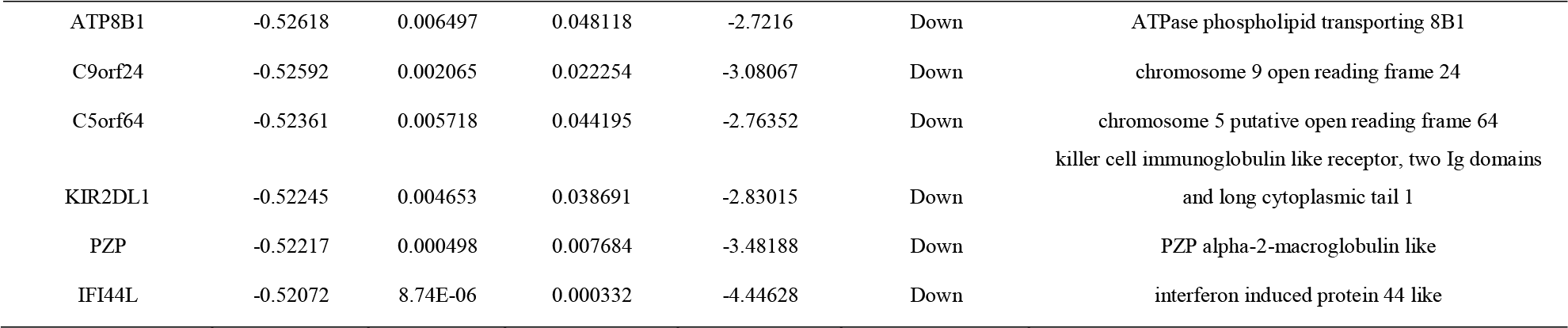
The statistical metrics for key differentially expressed genes (DEGs)

### GO and pathway enrichment analyses of DEGs

To obtain a deeper insight into the biological functions of DEGs, GO annotation and REACTOME pathway enrichment analyses were performed. The enriched GO terms were shown in Table 2. In the present investigation, up regulated genes were mainly enriched in transport (BP), localization (BP), cytoplasm (CC),intrinsic component (CC),enzyme regulator activity (MF) and protein binding (MF). Down regulated genes were mainly involved in multicellular organismal process (BP), developmental process (BP), cell periphery (CC), plasma membrane (CC), metal ion binding (MF) and cation binding (MF). The up regulated and down regulated genes from the REACTOME pathway enrichment analysis are shown in Table 3. The up regulated genes from the REACTOME pathway enrichment analysis were those for the neutrophil degranulation and immune system. The most down regulated genes from the REACTOME pathway enrichment were for extracellular matrix organization and diseases of metabolism.

**Table 2.**
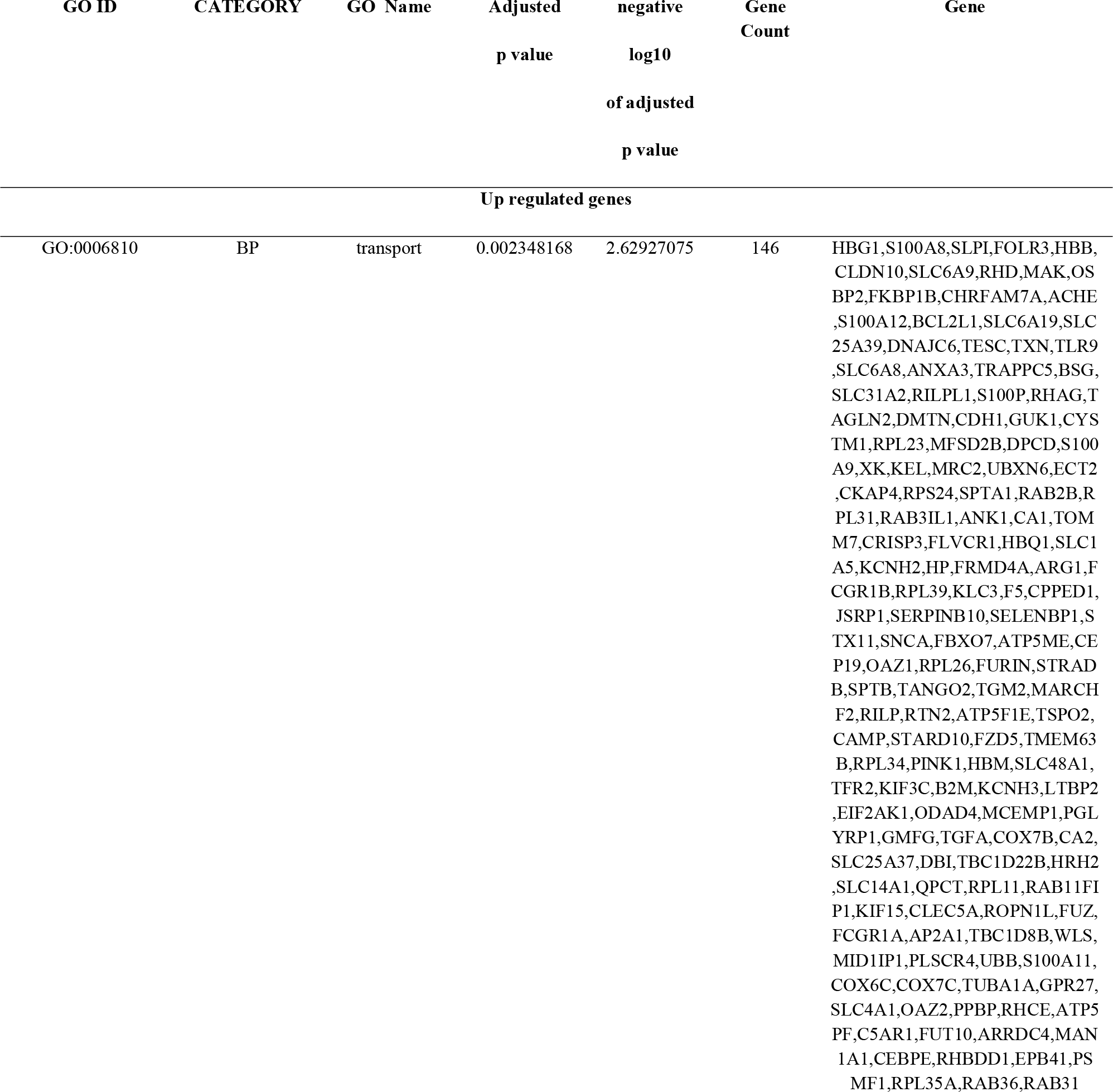
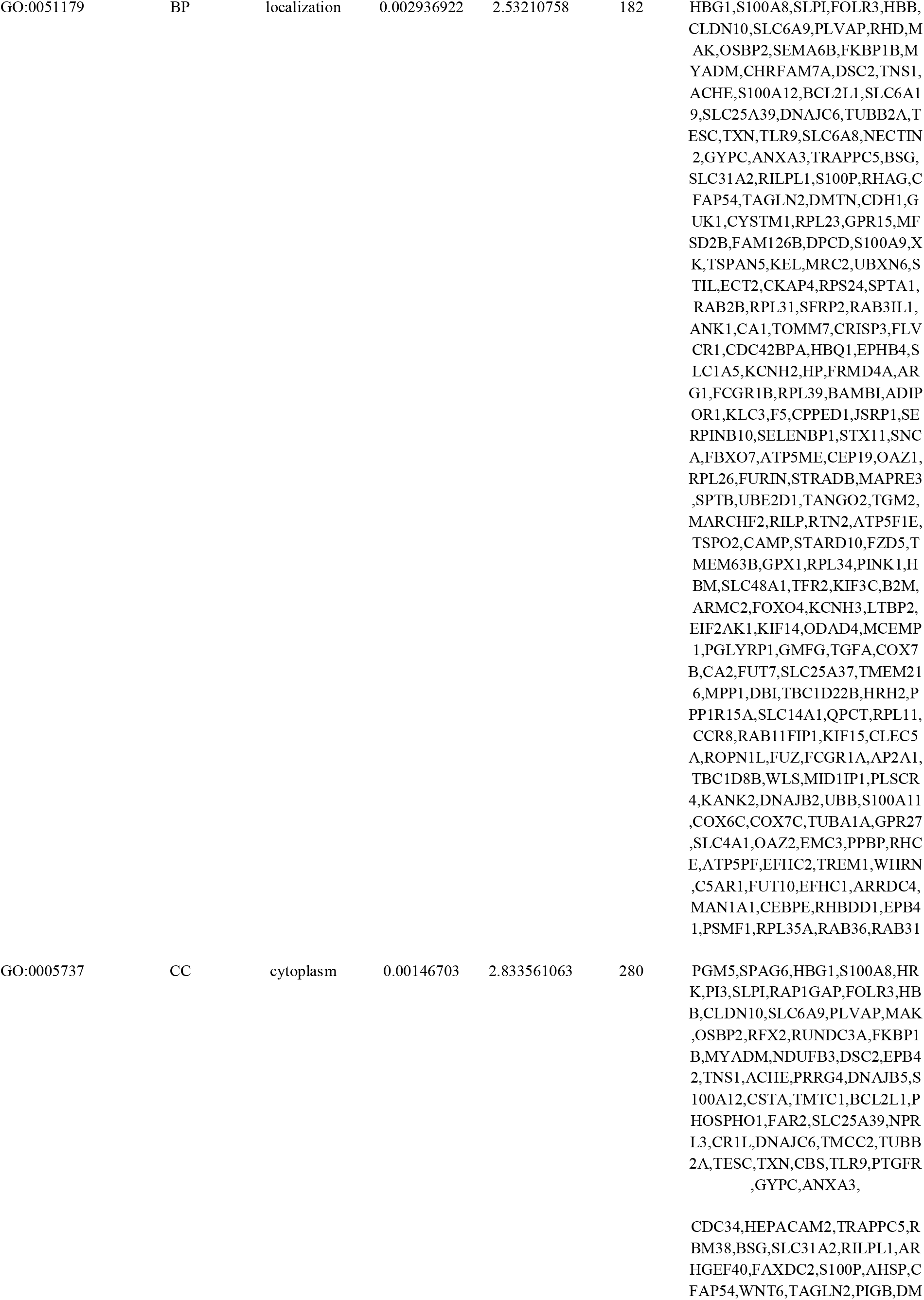
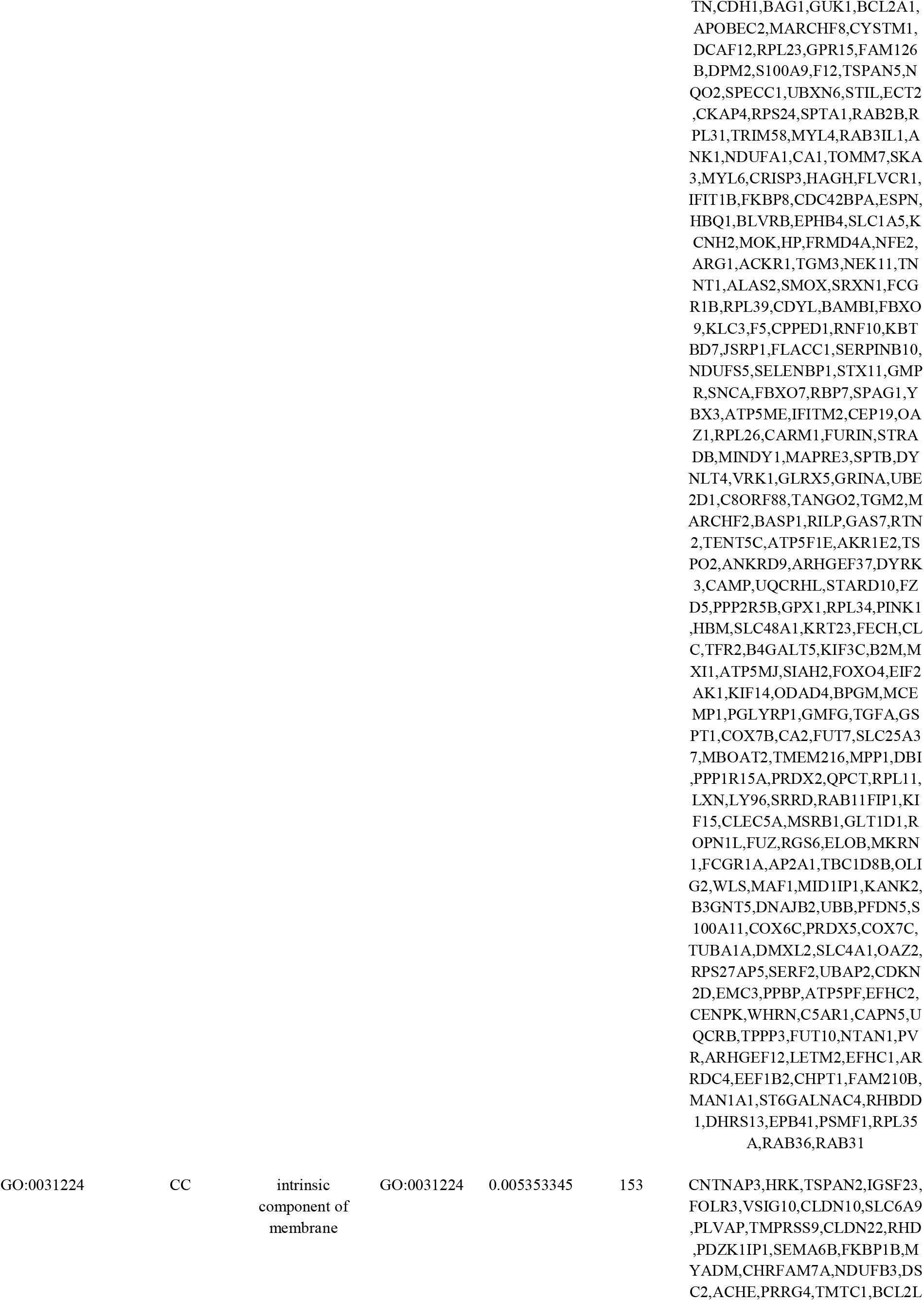
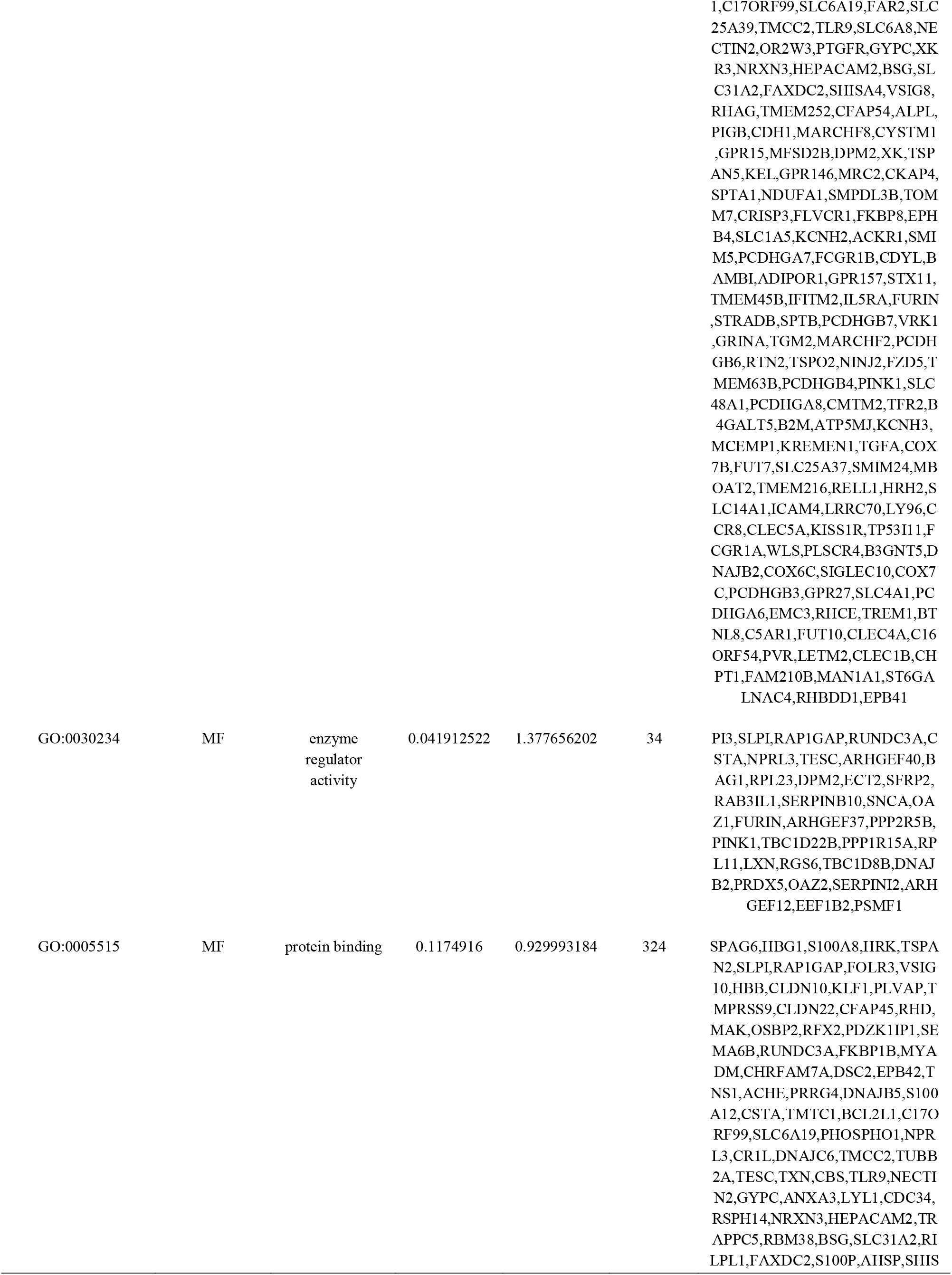
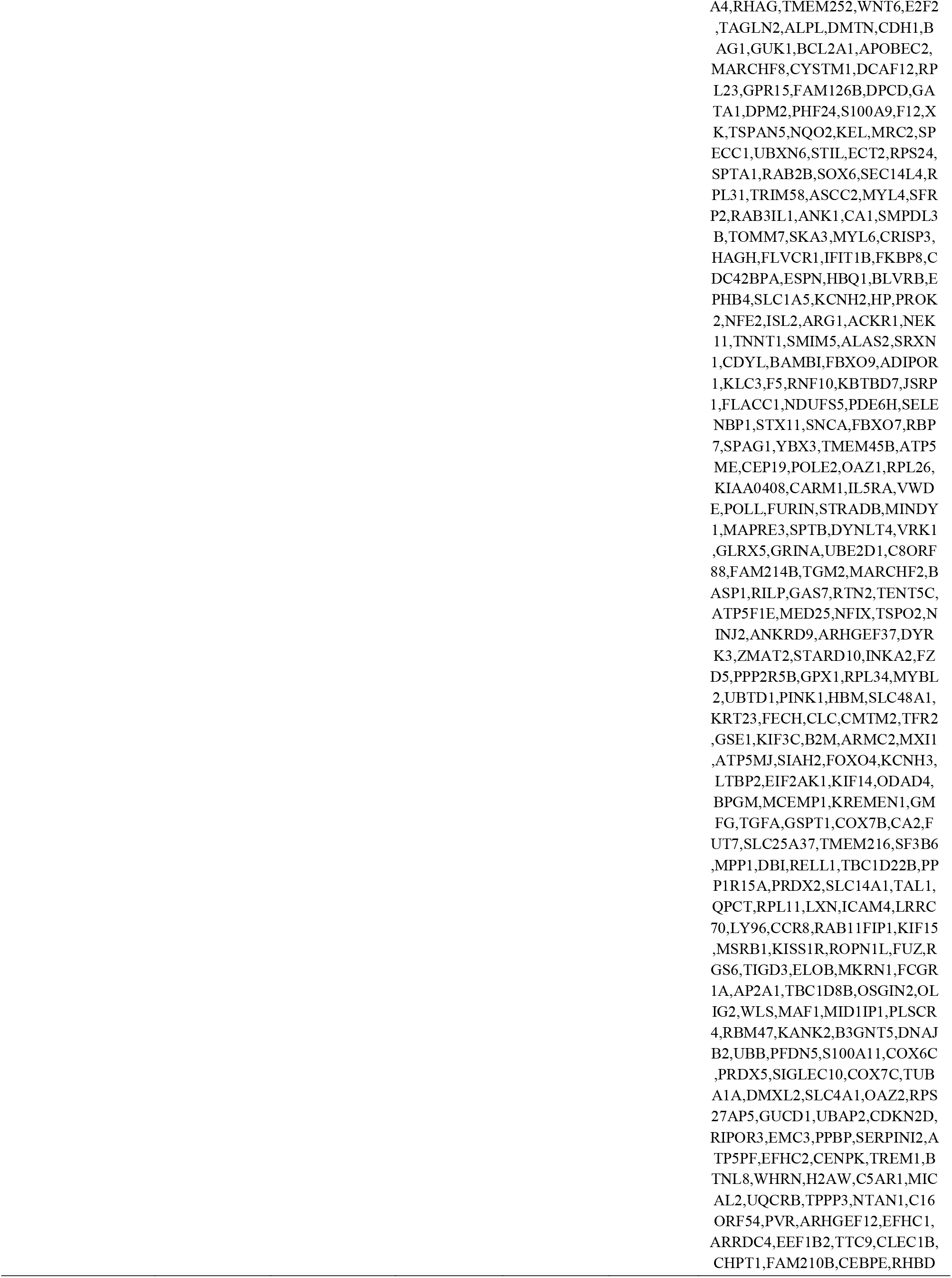
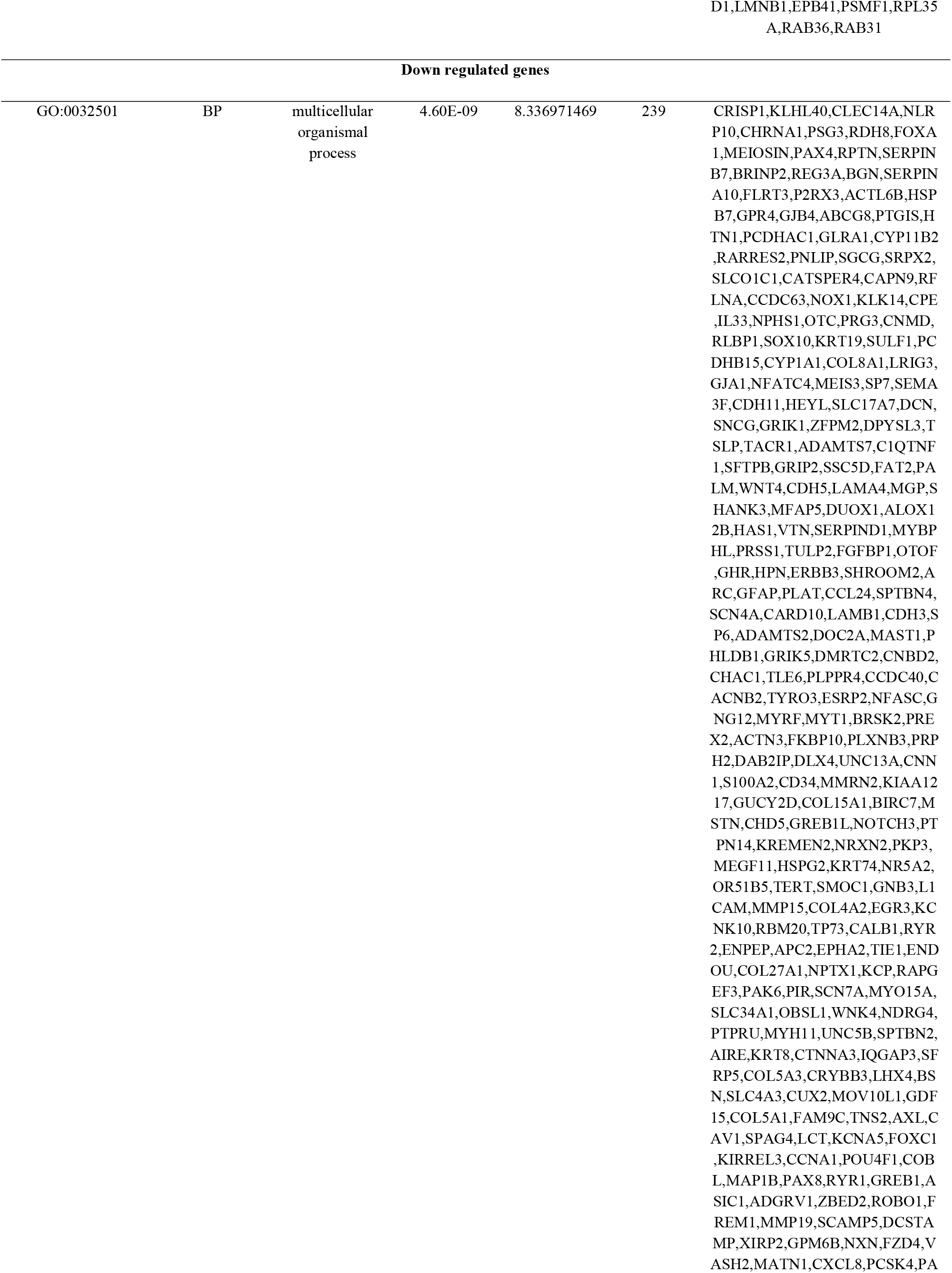
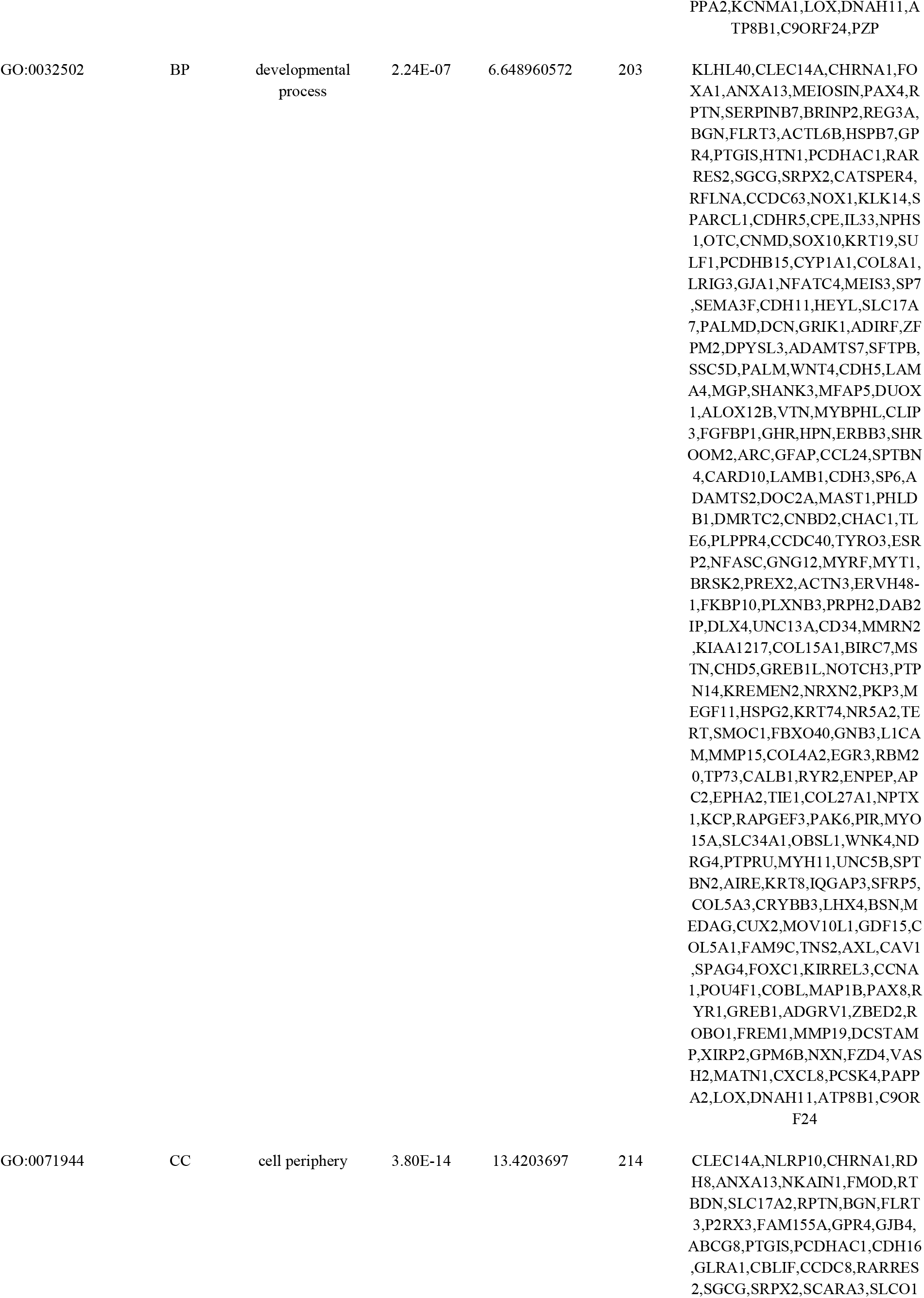
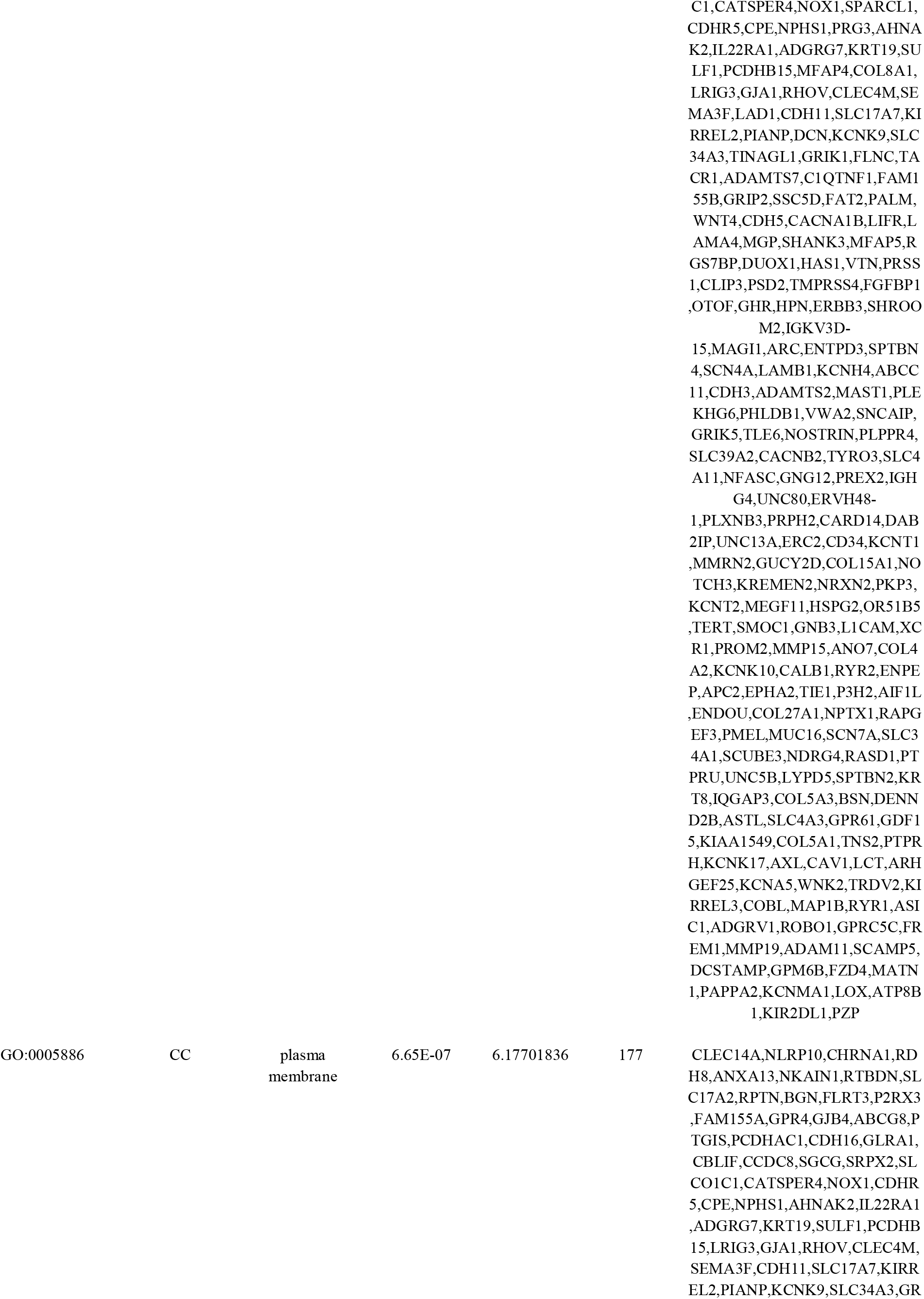
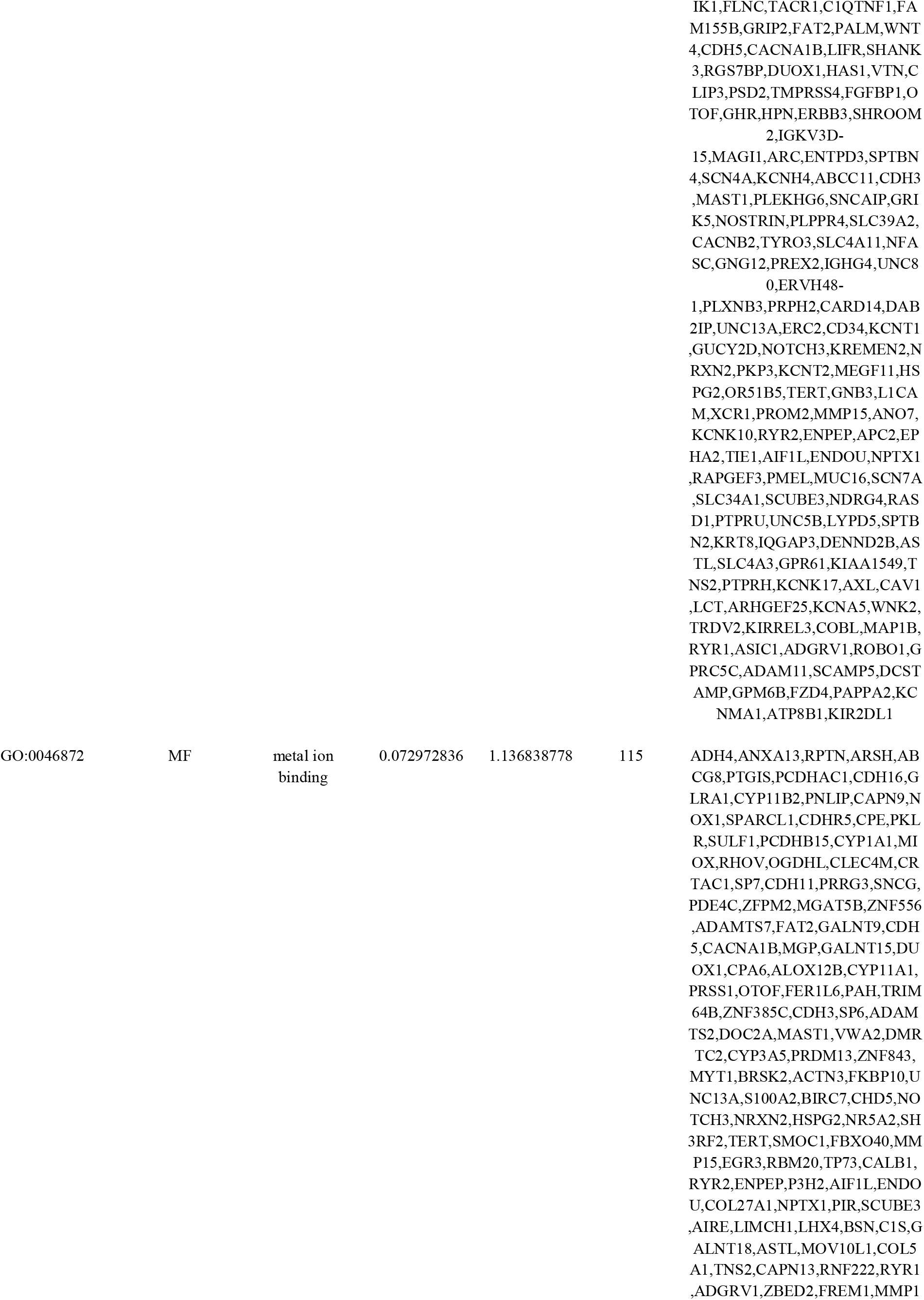
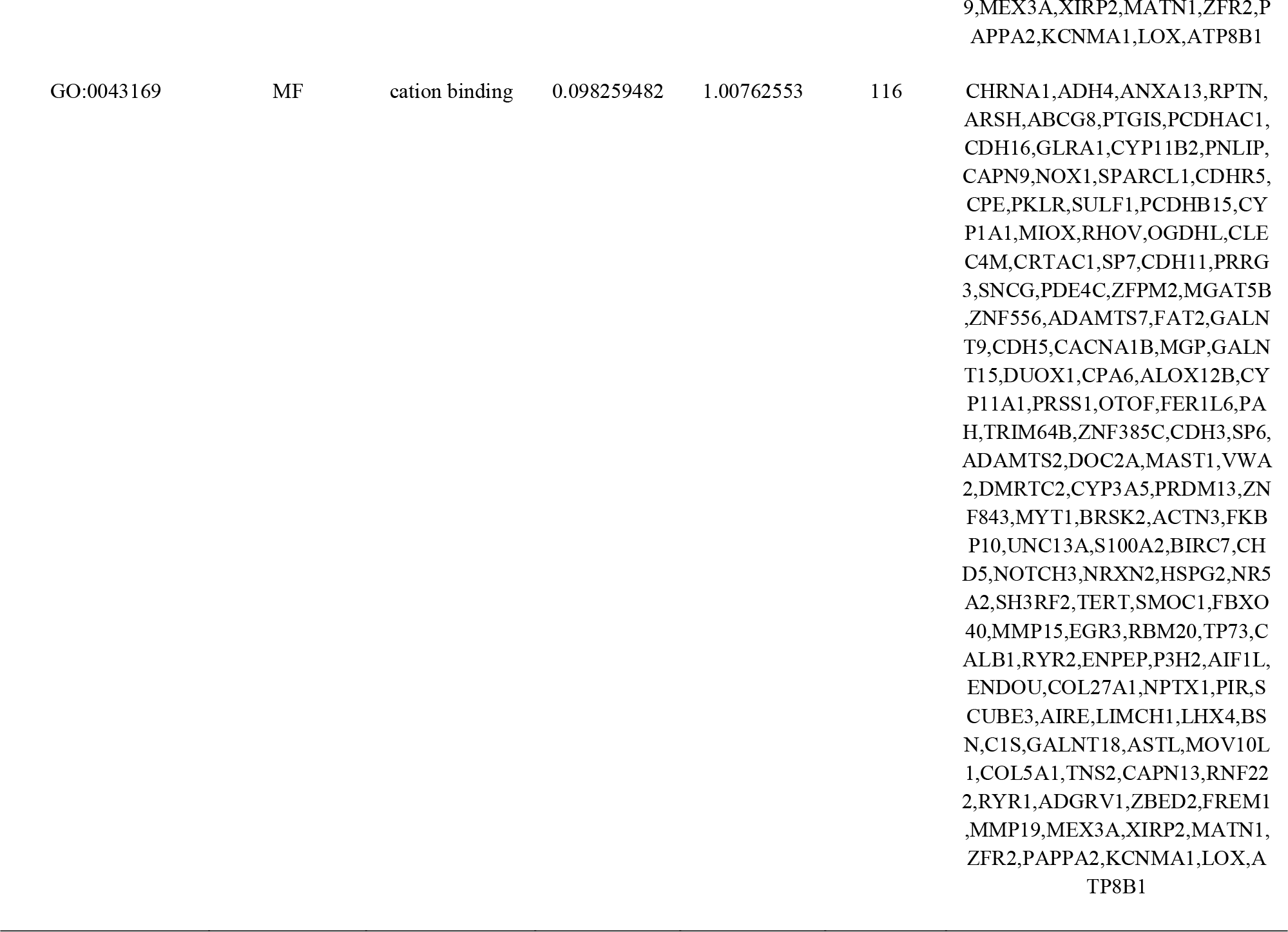
The enriched GO terms of the up and down regulated differentially expressed genes

**Table 3.**
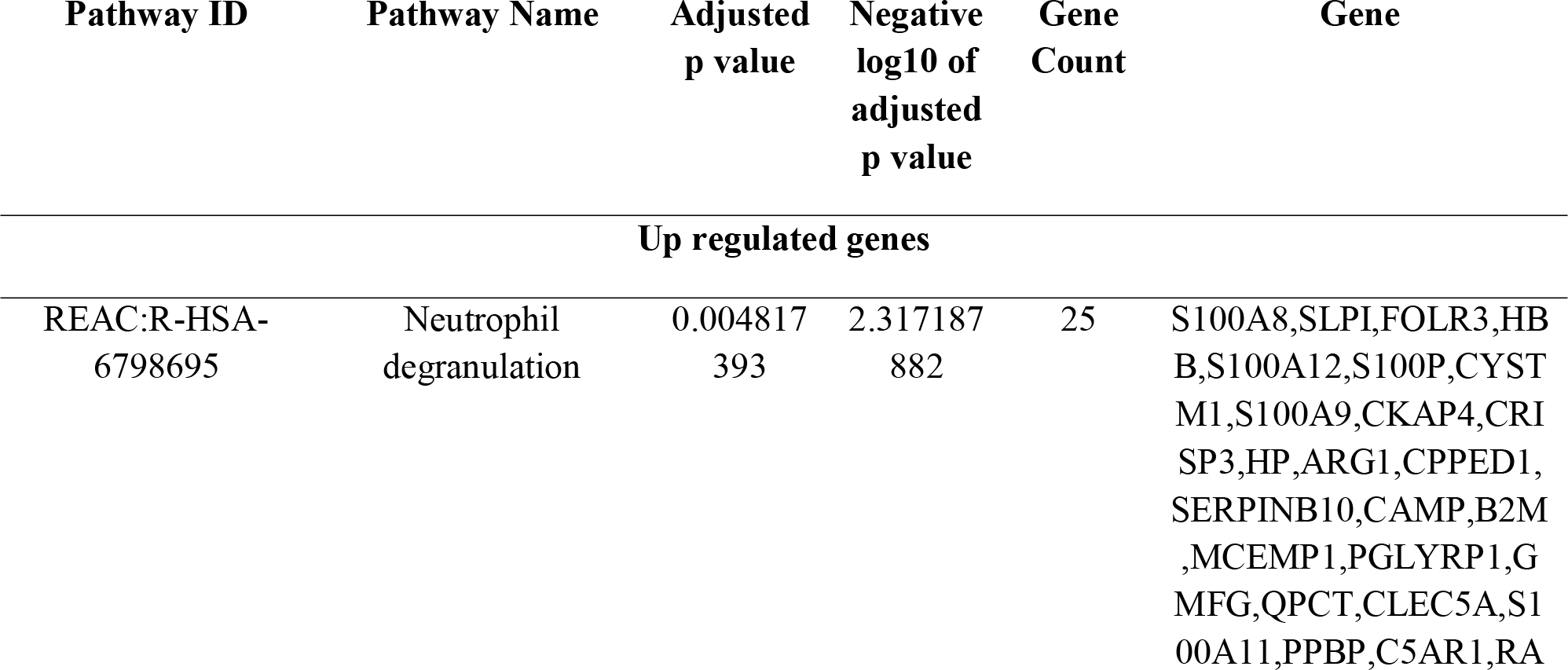
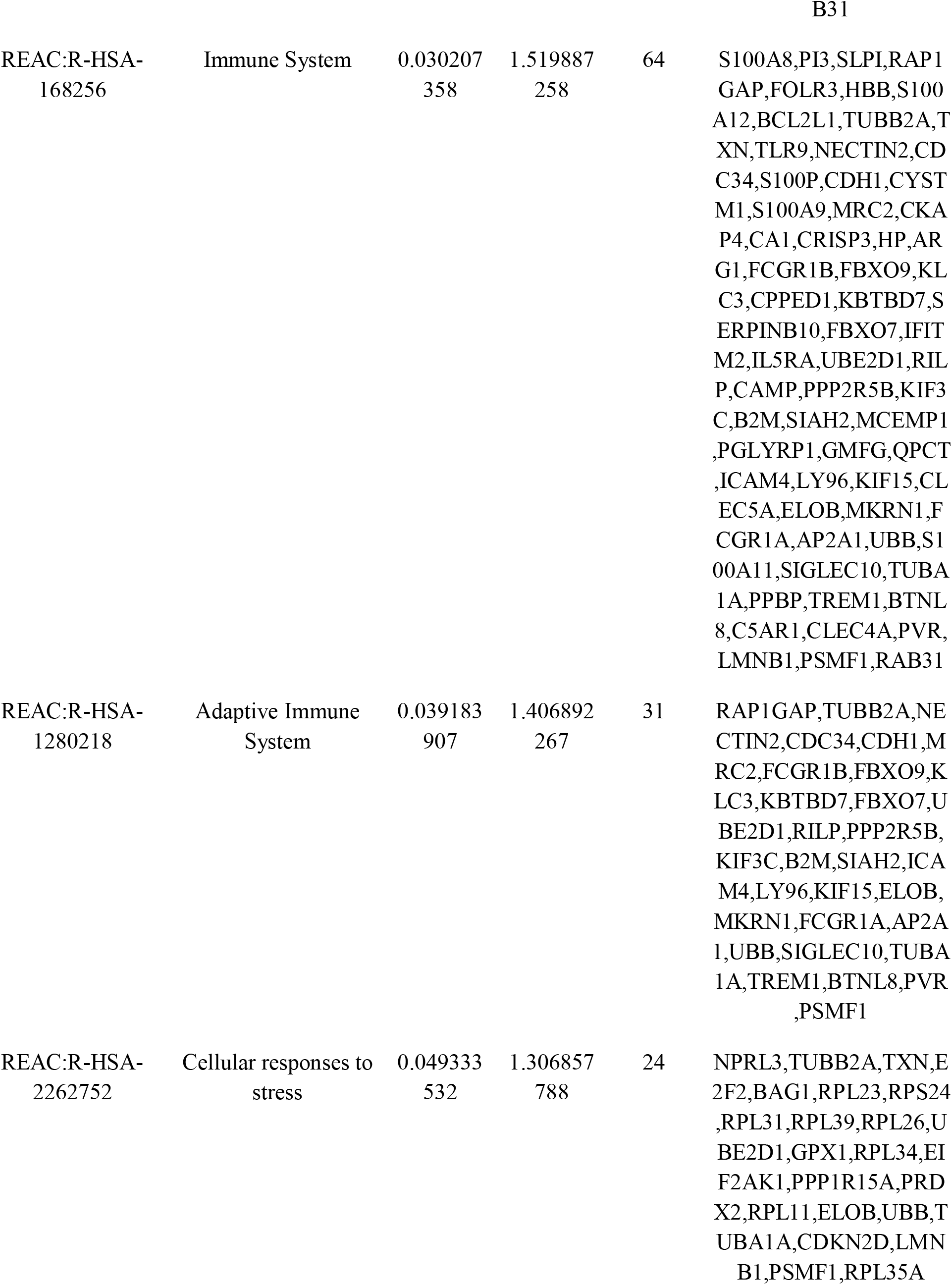
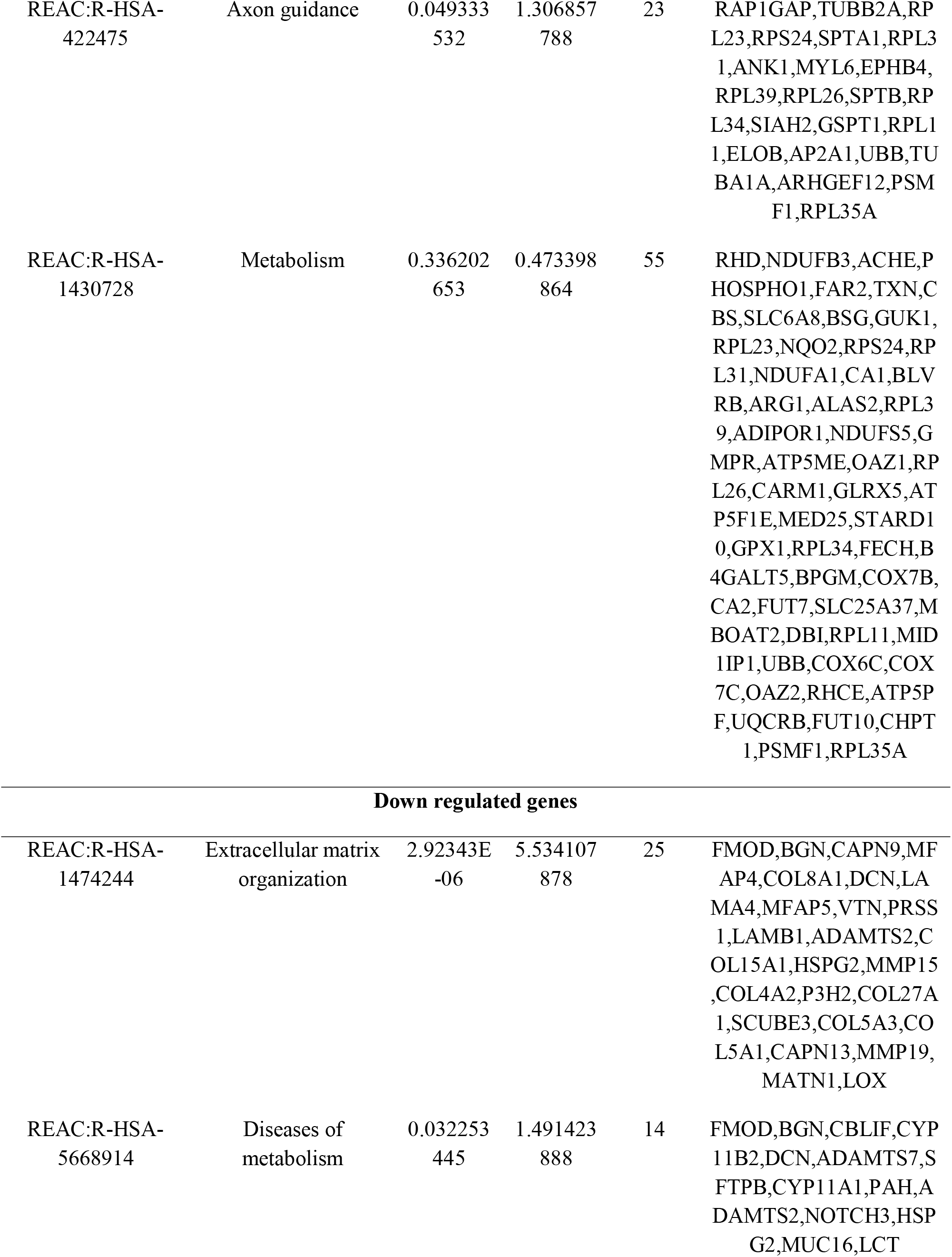
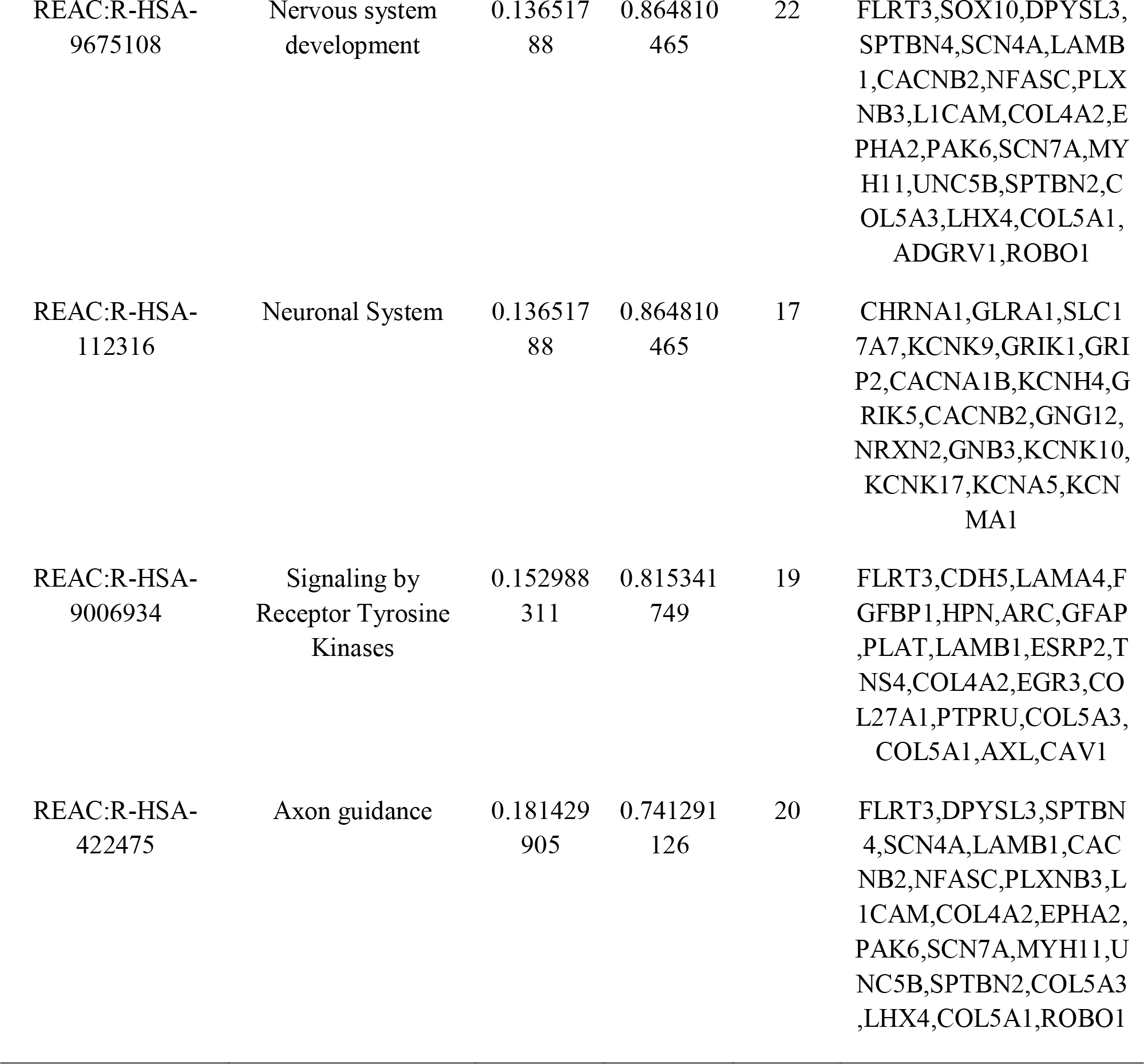
The enriched pathway terms of the up and down regulated differentially expressed genes

### Construction of the PPI and module analysis

Based on the information in the STRING database, the hub nodes with higher node degree, betweenness, stress and closeness were screened (Table 4). The PPI network contained 4360 nodes and 9761 edges (Fig.3). UBB, UBE2D1, TUBA1A, RPL11, RPS24, NOTCH3, CAV1, CNBD2, CCNA1 and MYH11 were the hub genes with the highest values of topological parameters (node degree, betweenness, stress and closeness). Furthermore, the 2 significant modules were extracted from the PPI network. Module 1 contained 71 gene nodes, including RPL39, RPL31, RPL23, RPL35A, RPL11, RPS24, UBB and RPL34 with 434 edges (Fig. 4A). Functional enrichment analysis of the hub genes in this module was mainly related to cellular responses to stress, axon guidance, metabolism, transport, localization, cytoplasm, immune system and protein binding. Module 2 contained 70 gene nodes, including FZD4, MSTN, EGR3, NOTCH3, PAX8, SFRP5 and WNT4 with 132 edges (Fig. 4B). Functional enrichment analysis of the hub genes in this module was mainly related to multicellular organismal process, developmental process, signaling by receptor tyrosine, diseases of metabolism and cell periphery.

**Fig. 3.**
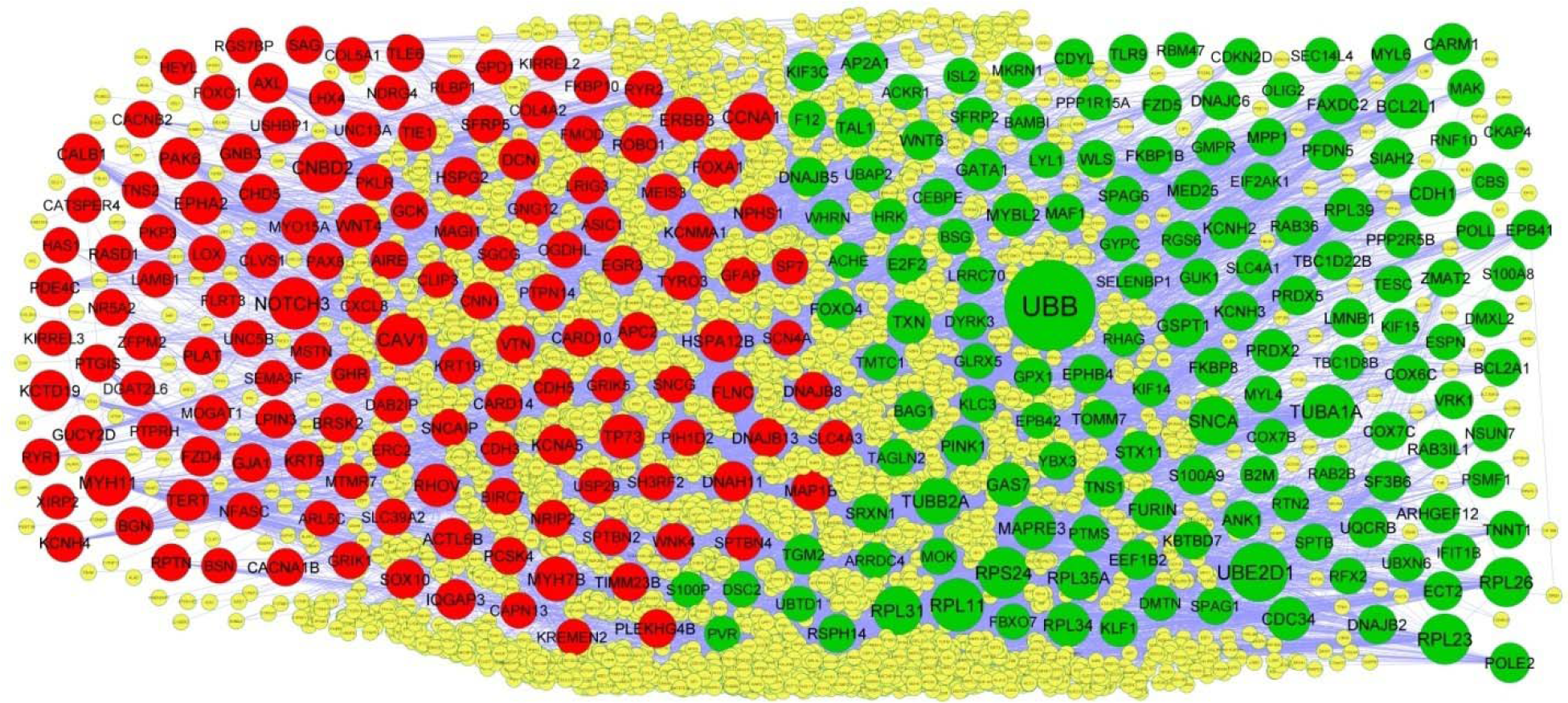
PPI network of DEGs. Up regulated genes are marked in green; down regulated genes are marked in red

**Fig. 4.**
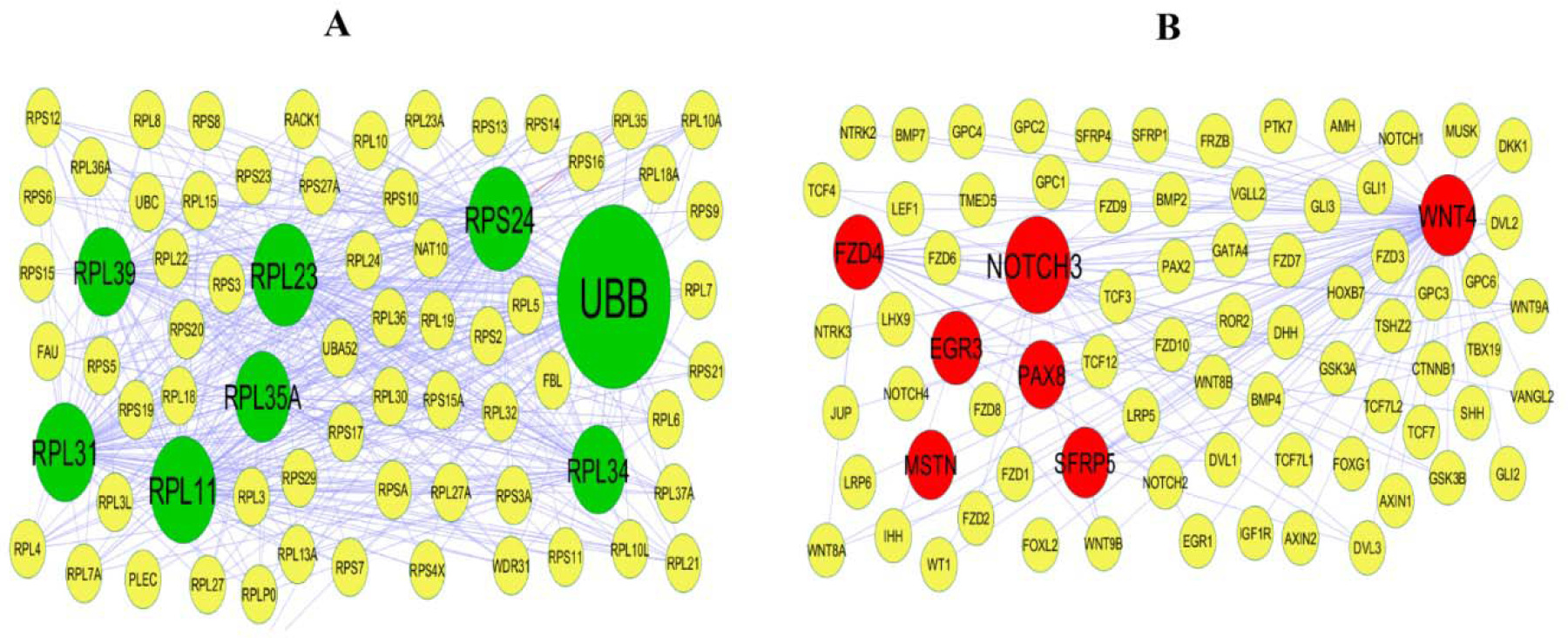
Modules selected from the DEG PPI between patients with BD and normal controls. (A) The most significant module was obtained from PPI network with 71 nodes and 434 edges for up regulated genes (B) The most significant module was obtained from PPI network with 70 nodes and 132 edges for down regulated genes. Up regulated genes are marked in green; down regulated genes are marked in red

**Table 4.** Topology table for up and down regulated genes

| Regulation | Node | Degree | Betweenness | Stress | Closeness |
| --- | --- | --- | --- | --- | --- |
| Up | UBB | 559 | 0.235727 | 99123156 | 0.380932 |
| Up | UBE2D1 | 235 | 0.058667 | 16144192 | 0.343012 |
| Up | TUBA1A | 181 | 0.057118 | 26071688 | 0.339539 |
| Up | RPL11 | 175 | 0.019652 | 9318422 | 0.337724 |
| Up | RPS24 | 160 | 0.021256 | 6961260 | 0.335617 |
| Up | RPL23 | 154 | 0.015493 | 7230944 | 0.336213 |
| Up | RPL26 | 142 | 0.01018 | 5252810 | 0.318175 |
| Up | SNCA | 135 | 0.044028 | 11884152 | 0.357207 |
| Up | RPL31 | 134 | 0.009194 | 4386284 | 0.334921 |
| Up | TUBB2A | 121 | 0.024242 | 10055526 | 0.327203 |
| Up | CDH1 | 108 | 0.034294 | 13478026 | 0.329678 |
| Up | BCL2L1 | 96 | 0.026458 | 6951328 | 0.320869 |
| Up | RPL35A | 93 | 0.003926 | 2632580 | 0.331735 |
| Up | RPL39 | 85 | 0.003148 | 1965128 | 0.33113 |
| Up | TXN | 85 | 0.019653 | 8351348 | 0.315572 |
| Up | RPL34 | 84 | 0.003072 | 1993106 | 0.331937 |
| Up | CDC34 | 82 | 0.015715 | 4631868 | 0.336862 |
| Up | MYBL2 | 82 | 0.023425 | 5077908 | 0.313304 |
| Up | GSPT1 | 81 | 0.015545 | 6377210 | 0.335824 |
| Up | GAS7 | 79 | 0.016953 | 4310252 | 0.328312 |
| Up | MAPRE3 | 78 | 0.006682 | 1696768 | 0.284456 |
| Up | FURIN | 69 | 0.025604 | 5609560 | 0.314957 |
| Up | GATA1 | 67 | 0.015983 | 4492280 | 0.303805 |
| Up | PRDX2 | 66 | 0.015033 | 3825266 | 0.310315 |
| Up | KCNH2 | 62 | 0.024215 | 6313724 | 0.315595 |
| Up | CARM1 | 60 | 0.013651 | 6121268 | 0.309368 |
| Up | AP2A1 | 59 | 0.017102 | 3708354 | 0.311513 |
| Up | PINK1 | 59 | 0.01307 | 3707782 | 0.328485 |
| Up | STX11 | 57 | 0.016174 | 8682644 | 0.263241 |
| Up | E2F2 | 55 | 0.007655 | 3723984 | 0.286324 |
| Up | PFDN5 | 50 | 0.008403 | 1968846 | 0.318687 |
| Up | SF3B6 | 49 | 0.017982 | 4002954 | 0.307361 |
| Up | UQCRB | 49 | 0.01182 | 3374160 | 0.304145 |
| Up | SIAH2 | 49 | 0.011784 | 2560508 | 0.315732 |
| Up | TNS1 | 48 | 0.005816 | 4296572 | 0.284159 |
| Up | POLE2 | 47 | 0.009518 | 6490004 | 0.298725 |
| Up | WNT6 | 45 | 0.002061 | 638626 | 0.277555 |
| Up | EEF1B2 | 43 | 0.009586 | 1699108 | 0.329205 |
| Up | FAXDC2 | 43 | 0.01133 | 8747764 | 0.234961 |
| Up | TGM2 | 43 | 0.010586 | 2833842 | 0.319715 |
| Up | FZD5 | 43 | 0.005074 | 1586660 | 0.312854 |
| Up | ARHGEF12 | 43 | 0.009673 | 3232552 | 0.307557 |
| Up | MED25 | 42 | 0.016459 | 4710392 | 0.29799 |
| Up | TAL1 | 41 | 0.008979 | 4419814 | 0.301661 |
| Up | EPB41 | 40 | 0.0089 | 3072508 | 0.309105 |
| Up | S100A9 | 40 | 0.00641 | 1886484 | 0.271945 |
| Up | FOXO4 | 40 | 0.008381 | 5014768 | 0.312967 |
| Up | PRDX5 | 39 | 0.007035 | 2185670 | 0.304825 |
| Up | RAB3IL1 | 39 | 0.006139 | 4908502 | 0.245481 |
| Up | BAG1 | 38 | 0.005118 | 2205794 | 0.308777 |
| Up | MYL6 | 37 | 0.007047 | 2273202 | 0.309237 |
| Up | FKBP8 | 36 | 0.009884 | 3006680 | 0.313259 |
| Up | LMNB1 | 36 | 0.008384 | 2689680 | 0.309171 |
| Up | DNAJB2 | 36 | 0.005128 | 2614468 | 0.30611 |
| Up | EPHB4 | 36 | 0.001705 | 1209330 | 0.267473 |
| Up | TBC1D22B | 36 | 0.00895 | 3732342 | 0.241724 |
| Up | DNAJB5 | 36 | 0.004281 | 2275500 | 0.308013 |
| Up | RAB36 | 35 | 0.009686 | 3351236 | 0.240284 |
| Up | KIF3C | 35 | 0.005417 | 2129670 | 0.254897 |
| Up | TLR9 | 35 | 0.007537 | 5597554 | 0.276762 |
| Up | SPAG6 | 35 | 0.011612 | 8123614 | 0.237548 |
| Up | KCNH3 | 34 | 2.73E-04 | 62058 | 0.231726 |
| Up | ECT2 | 33 | 0.006266 | 2585608 | 0.303087 |
| Up | DNAJC6 | 33 | 0.002948 | 1245626 | 0.277449 |
| Up | CDYL | 31 | 0.009246 | 2991466 | 0.295806 |
| Up | SRXN1 | 30 | 0.00391 | 1142608 | 0.272114 |
| Up | SFRP2 | 29 | 6.53E-04 | 204668 | 0.262638 |
| Up | ZMAT2 | 29 | 0.008292 | 2361290 | 0.212448 |
| Up | MAF1 | 29 | 0.004047 | 3116836 | 0.260877 |
| Up | POLL | 28 | 0.005859 | 2925224 | 0.297827 |
| Up | RSPH14 | 27 | 0.002068 | 3695794 | 0.251123 |
| Up | VRK1 | 27 | 0.007833 | 2906102 | 0.300227 |
| Up | COX6C | 27 | 0.003684 | 1303716 | 0.294269 |
| Up | COX7C | 25 | 3.07E-04 | 213844 | 0.245232 |
| Up | ANK1 | 25 | 0.010597 | 2354792 | 0.297502 |
| Up | MPP1 | 25 | 0.004584 | 551834 | 0.268626 |
| Up | FKBP1B | 25 | 0.006692 | 3341772 | 0.266899 |
| Up | TNNT1 | 25 | 0.001358 | 725396 | 0.24648 |
| Up | KLF1 | 24 | 0.003809 | 1989514 | 0.311669 |
| Up | PPP2R5B | 23 | 0.005919 | 3281420 | 0.25605 |
| Up | BCL2A1 | 23 | 0.00143 | 938544 | 0.248419 |
| Up | TOMM7 | 23 | 0.006143 | 1676746 | 0.293259 |
| Up | KBTBD7 | 23 | 0.006534 | 1678442 | 0.29893 |
| Up | B2M | 23 | 0.008498 | 2045152 | 0.293733 |
| Up | WLS | 22 | 0.002225 | 638452 | 0.30139 |
| Up | PSMF1 | 21 | 0.00136 | 611172 | 0.294288 |
| Up | GPX1 | 21 | 0.001777 | 387698 | 0.264567 |
| Up | YBX3 | 20 | 0.005034 | 1487506 | 0.30611 |
| Up | F12 | 20 | 0.004622 | 1167234 | 0.310271 |
| Up | UBXN6 | 20 | 0.004055 | 636686 | 0.308864 |
| Up | RGS6 | 20 | 0.001883 | 2315560 | 0.214065 |
| Up | GMPR | 19 | 0.00433 | 2482218 | 0.235673 |
| Up | SPAG1 | 19 | 0.004494 | 1266070 | 0.304591 |
| Up | LRRC70 | 19 | 0.001794 | 1288356 | 0.273824 |
| Up | CEBPE | 18 | 0.003199 | 612550 | 0.258602 |
| Up | CDKN2D | 17 | 0.00174 | 2103692 | 0.246689 |
| Up | TAGLN2 | 17 | 0.002546 | 978432 | 0.294846 |
| Up | IFIT1B | 17 | 8.79E-04 | 366642 | 0.271928 |
| Up | DMXL2 | 17 | 0.003574 | 638106 | 0.261707 |
| Up | MAK | 16 | 0.00101 | 267508 | 0.276043 |
| Up | GUK1 | 16 | 0.002087 | 746630 | 0.304655 |
| Up | MYL4 | 15 | 3.56E-05 | 18524 | 0.228375 |
| Up | SPTB | 15 | 0.002888 | 608416 | 0.299402 |
| Up | NSUN7 | 15 | 7.29E-04 | 1321010 | 0.22062 |
| Up | TMTC1 | 15 | 9.74E-04 | 350788 | 0.27142 |
| Up | CBS | 13 | 0.001289 | 274140 | 0.321365 |
| Up | DYRK3 | 13 | 0.001509 | 1070842 | 0.242436 |
| Up | KLC3 | 13 | 0.00172 | 876438 | 0.245771 |
| Up | EIF2AK1 | 12 | 0.003598 | 800812 | 0.299938 |
| Up | PTMS | 12 | 0.002509 | 780322 | 0.297685 |
| Up | MOK | 12 | 0.001023 | 208890 | 0.26979 |
| Up | RFX2 | 11 | 0.001037 | 1138250 | 0.211715 |
| Up | ESPN | 11 | 0.004672 | 1411700 | 0.241122 |
| Up | TESC | 9 | 0.002778 | 933996 | 0.1981 |
| Up | RNF10 | 8 | 0.001202 | 172420 | 0.304655 |
| Up | ISL2 | 6 | 0.001838 | 963308 | 0.210052 |
| Up | KIF15 | 5 | 0.001392 | 381452 | 0.228256 |
| Up | KIF14 | 5 | 9.79E-04 | 221060 | 0.294328 |
| Up | RAB2B | 5 | 0.001023 | 297048 | 0.224795 |
| Up | SLC4A1 | 2 | 1.60E-05 | 9030 | 0.237742 |
| Up | HRK | 2 | 8.89E-06 | 14930 | 0.243193 |
| Up | DSC2 | 2 | 1.86E-04 | 70352 | 0.239809 |
| Up | FBXO7 | 2 | 7.49E-06 | 4460 | 0.248376 |
| Up | DMTN | 2 | 2.47E-05 | 13250 | 0.264358 |
| Up | EPB42 | 2 | 1.60E-05 | 9030 | 0.237742 |
| Up | GYPC | 2 | 6.00E-05 | 9802 | 0.242315 |
| Up | SEC14L4 | 2 | 7.26E-06 | 3288 | 0.219619 |
| Up | UBTD1 | 2 | 2.33E-05 | 2974 | 0.257305 |
| Up | PVR | 2 | 2.28E-04 | 152670 | 0.250014 |
| Up | COX7B | 2 | 0 | 0 | 0.22741 |
| Up | LYL1 | 1 | 0 | 0 | 0.231763 |
| Up | RTN2 | 1 | 0 | 0 | 0.242936 |
| Up | MKRN1 | 1 | 0 | 0 | 0.241724 |
| Up | BSG | 1 | 0 | 0 | 0.251225 |
| Up | OLIG2 | 1 | 0 | 0 | 0.235405 |
| Up | S100P | 1 | 0 | 0 | 0.212479 |
| Up | S100A8 | 1 | 0 | 0 | 0.213813 |
| Up | GLRX5 | 1 | 0 | 0 | 0.239888 |
| Up | RBM47 | 1 | 0 | 0 | 0.228292 |
| Up | SELENBP1 | 1 | 0 | 0 | 0.209225 |
| Up | ARRDC4 | 1 | 0 | 0 | 0.247179 |
| Up | PPP1R15A | 1 | 0 | 0 | 0.23594 |
| Up | ACHE | 1 | 0 | 0 | 0.230355 |
| Up | TBC1D8B | 1 | 0 | 0 | 0.193742 |
| Up | UBAP2 | 1 | 0 | 0 | 0.233626 |
| Up | WHRN | 1 | 0 | 0 | 0.170068 |
| Up | ACKR1 | 1 | 0 | 0 | 0.200635 |
| Up | RHAG | 1 | 0 | 0 | 0.2293 |
| Up | BAMBI | 1 | 0 | 0 | 0.238314 |
| Up | CKAP4 | 1 | 0 | 0 | 0.209719 |
| Down | NOTCH3 | 168 | 0.055079 | 18551684 | 0.337645 |
| Down | CAV1 | 166 | 0.054912 | 23161840 | 0.335488 |
| Down | CNBD2 | 152 | 0.055891 | 22160854 | 0.295325 |
| Down | CCNA1 | 125 | 0.03072 | 15117004 | 0.330027 |
| Down | MYH11 | 120 | 0.029866 | 14461722 | 0.325858 |
| Down | ERBB3 | 98 | 0.027362 | 10549184 | 0.327621 |
| Down | MYH7B | 93 | 0.013364 | 7918670 | 0.313057 |
| Down | TP73 | 91 | 0.019381 | 12310222 | 0.322435 |
| Down | WNT4 | 81 | 0.013753 | 3043608 | 0.293733 |
| Down | EPHA2 | 81 | 0.019399 | 10587382 | 0.321365 |
| Down | RHOV | 76 | 0.01927 | 3807074 | 0.285126 |
| Down | HSPA12B | 73 | 0.015451 | 6907454 | 0.276534 |
| Down | TERT | 66 | 0.016789 | 9086160 | 0.318757 |
| Down | FLNC | 65 | 0.020963 | 4387718 | 0.315367 |
| Down | PAK6 | 64 | 0.012017 | 6881200 | 0.290001 |
| Down | ACTL6B | 60 | 0.015257 | 8049238 | 0.30087 |
| Down | DCN | 59 | 0.016307 | 8557802 | 0.291826 |
| Down | KCTD19 | 57 | 0.015943 | 23365060 | 0.23016 |
| Down | FOXA1 | 56 | 0.010658 | 2393526 | 0.28858 |
| Down | CALB1 | 54 | 0.006522 | 1963692 | 0.277803 |
| Down | IQGAP3 | 51 | 0.003166 | 1814440 | 0.305423 |
| Down | DNAH11 | 49 | 0.005505 | 1652270 | 0.266052 |
| Down | KCNA5 | 48 | 0.012218 | 3239446 | 0.296935 |
| Down | FZD4 | 48 | 0.010838 | 2607416 | 0.333665 |
| Down | SOX10 | 46 | 0.017944 | 3502486 | 0.307861 |
| Down | HSPG2 | 46 | 0.011717 | 5959184 | 0.268494 |
| Down | MAP1B | 45 | 0.010903 | 2542390 | 0.314479 |
| Down | AXL | 45 | 0.006626 | 3832530 | 0.31178 |
| Down | GNB3 | 40 | 0.008299 | 6860166 | 0.248249 |
| Down | KRT8 | 40 | 0.010354 | 2664144 | 0.315321 |
| Down | EGR3 | 40 | 0.004773 | 1445998 | 0.289962 |
| Down | GHR | 39 | 0.00729 | 2538360 | 0.302142 |
| Down | CHD5 | 37 | 0.007617 | 2863732 | 0.297787 |
| Down | BRSK2 | 37 | 0.007801 | 3221358 | 0.300787 |
| Down | TYRO3 | 35 | 0.004255 | 2267676 | 0.30553 |
| Down | GJA1 | 35 | 0.005356 | 1960438 | 0.314797 |
| Down | KCNH4 | 34 | 2.73E-04 | 62058 | 0.231726 |
| Down | DNAJB13 | 34 | 0.001534 | 1259956 | 0.27685 |
| Down | RASD1 | 33 | 0.004682 | 3193548 | 0.267752 |
| Down | GUCY2D | 33 | 0.007445 | 2911470 | 0.24464 |
| Down | PLEKHG4B | 32 | 0.008619 | 3574886 | 0.311313 |
| Down | PDE4C | 32 | 0.002931 | 1507002 | 0.253666 |
| Down | CARD10 | 31 | 0.005693 | 2092664 | 0.312608 |
| Down | CDH5 | 31 | 0.005122 | 903600 | 0.288867 |
| Down | KCNMA1 | 31 | 0.005838 | 1401676 | 0.292511 |
| Down | VTN | 30 | 0.006914 | 3665516 | 0.250201 |
| Down | BGN | 30 | 0.004661 | 2638574 | 0.274358 |
| Down | TIE1 | 30 | 0.003627 | 4131006 | 0.247361 |
| Down | TIMM23B | 30 | 0.00781 | 2399166 | 0.29411 |
| Down | MAGI1 | 30 | 0.007946 | 1054376 | 0.277643 |
| Down | DNAJB8 | 28 | 8.34E-04 | 868048 | 0.265534 |
| Down | LPIN3 | 28 | 0.007833 | 9933232 | 0.240139 |
| Down | APC2 | 27 | 0.003535 | 1439816 | 0.315048 |
| Down | ROBO1 | 27 | 0.007344 | 2225388 | 0.303235 |
| Down | PLAT | 26 | 0.005969 | 764484 | 0.265356 |
| Down | NPHS1 | 26 | 0.005821 | 1441232 | 0.309984 |
| Down | PTGIS | 25 | 0.007512 | 1674446 | 0.276955 |
| Down | SNCAIP | 25 | 0.00324 | 1055652 | 0.331786 |
| Down | SFRP5 | 25 | 6.26E-04 | 583618 | 0.250604 |
| Down | CARD14 | 24 | 0.001554 | 716090 | 0.264888 |
| Down | TLE6 | 23 | 0.004166 | 726774 | 0.275346 |
| Down | PCSK4 | 23 | 0.002496 | 1235892 | 0.24662 |
| Down | CACNA1B | 23 | 0.008952 | 1641246 | 0.301786 |
| Down | SAG | 23 | 0.005896 | 2535550 | 0.235545 |
| Down | GCK | 22 | 0.006185 | 3432410 | 0.237898 |
| Down | AIRE | 22 | 0.00462 | 3733612 | 0.25682 |
| Down | NRIP2 | 22 | 0.004742 | 3094270 | 0.251196 |
| Down | RPTN | 22 | 0.006492 | 5712046 | 0.230696 |
| Down | BIRC7 | 21 | 0.002382 | 795084 | 0.302498 |
| Down | LOX | 21 | 0.00354 | 1579368 | 0.251573 |
| Down | RYR2 | 21 | 0.002756 | 921910 | 0.255555 |
| Down | ZFPM2 | 21 | 0.001959 | 502922 | 0.261441 |
| Down | DAB2IP | 20 | 0.001461 | 875988 | 0.269707 |
| Down | FMOD | 20 | 0.001634 | 1281220 | 0.26939 |
| Down | COL4A2 | 20 | 0.003725 | 930468 | 0.261503 |
| Down | MSTN | 19 | 0.005012 | 504404 | 0.267538 |
| Down | CNN1 | 19 | 0.00113 | 966030 | 0.25581 |
| Down | NR5A2 | 19 | 0.003478 | 1735388 | 0.259418 |
| Down | PTPN14 | 18 | 0.002998 | 656768 | 0.310669 |
| Down | GRIK1 | 18 | 0.006648 | 1099152 | 0.297644 |
| Down | WNK4 | 17 | 0.004851 | 1167040 | 0.303552 |
| Down | SPTBN2 | 17 | 0.002019 | 799022 | 0.297299 |
| Down | LRIG3 | 17 | 0.001188 | 1092168 | 0.274496 |
| Down | CACNB2 | 17 | 0.003678 | 446370 | 0.259743 |
| Down | SNCG | 17 | 0.001579 | 901622 | 0.256126 |
| Down | LAMB1 | 16 | 0.004945 | 931360 | 0.299279 |
| Down | CXCL8 | 16 | 0.004475 | 3208596 | 0.250979 |
| Down | RYR1 | 16 | 0.002088 | 754680 | 0.258848 |
| Down | SH3RF2 | 16 | 0.002458 | 1059370 | 0.254555 |
| Down | OGDHL | 16 | 0.00406 | 1083468 | 0.293535 |
| Down | MEIS3 | 16 | 0.004187 | 895770 | 0.241992 |
| Down | TNS2 | 15 | 0.001495 | 269368 | 0.26959 |
| Down | KRT19 | 15 | 8.70E-04 | 363806 | 0.300559 |
| Down | GFAP | 14 | 0.002533 | 1030522 | 0.306497 |
| Down | NFASC | 14 | 0.0046 | 735652 | 0.234267 |
| Down | PKLR | 14 | 0.004287 | 856214 | 0.213342 |
| Down | MYO15A | 13 | 0.004408 | 856128 | 0.204908 |
| Down | SLC39A2 | 13 | 0.005041 | 2191092 | 0.197885 |
| Down | UNC5B | 13 | 0.004058 | 2253212 | 0.200028 |
| Down | GNG12 | 13 | 0.002494 | 911124 | 0.300683 |
| Down | PAX8 | 13 | 0.001585 | 570488 | 0.252491 |
| Down | ERC2 | 12 | 0.004146 | 727408 | 0.207098 |
| Down | USHBP1 | 11 | 0.004373 | 1170750 | 0.203113 |
| Down | SP7 | 10 | 0.001802 | 590558 | 0.304676 |
| Down | PKP3 | 10 | 0.001892 | 762932 | 0.219841 |
| Down | SPTBN4 | 9 | 0.00142 | 157756 | 0.241643 |
| Down | PIH1D2 | 9 | 0.001201 | 909904 | 0.256986 |
| Down | CAPN13 | 8 | 9.25E-04 | 542252 | 0.225761 |
| Down | CLIP3 | 8 | 0.001099 | 489142 | 0.255256 |
| Down | HAS1 | 7 | 5.92E-04 | 127254 | 0.258833 |
| Down | USP29 | 7 | 9.47E-04 | 567976 | 0.241375 |
| Down | ARL5C | 7 | 0.001564 | 442358 | 0.226336 |
| Down | KREMEN2 | 7 | 1.10E-04 | 16216 | 0.210305 |
| Down | FKBP10 | 7 | 5.41E-04 | 472476 | 0.207928 |
| Down | PTPRH | 7 | 6.42E-04 | 94260 | 0.256231 |
| Down | FOXC1 | 6 | 7.06E-04 | 245654 | 0.23594 |
| Down | MTMR7 | 5 | 9.40E-04 | 543354 | 0.225738 |
| Down | RLBP1 | 5 | 9.85E-04 | 174658 | 0.221033 |
| Down | SEMA3F | 4 | 0.001022 | 226518 | 0.19297 |
| Down | LHX4 | 1 | 0 | 0 | 0.173596 |
| Down | XIRP2 | 1 | 0 | 0 | 0.239769 |
| Down | SGCG | 1 | 0 | 0 | 0.239769 |
| Down | GPD1 | 1 | 0 | 0 | 0.243221 |
| Down | DGAT2L6 | 1 | 0 | 0 | 0.193647 |
| Down | KIRREL2 | 1 | 0 | 0 | 0.236645 |
| Down | CLVS1 | 1 | 0 | 0 | 0.216413 |
| Down | GRIK5 | 1 | 0 | 0 | 0.229385 |
| Down | ASIC1 | 1 | 0 | 0 | 0.21732 |
| Down | RGS7BP | 1 | 0 | 0 | 0.176328 |
| Down | CDH3 | 1 | 0 | 0 | 0.247952 |
| Down | SLC4A3 | 1 | 0 | 0 | 0.2293 |
| Down | BSN | 1 | 0 | 0 | 0.171574 |
| Down | FLRT3 | 1 | 0 | 0 | 0.166692 |
| Down | COL5A1 | 1 | 0 | 0 | 0.225913 |
| Down | HEYL | 1 | 0 | 0 | 0.252432 |
| Down | NDRG4 | 1 | 0 | 0 | 0.198887 |
| Down | CATSPER4 | 1 | 0 | 0 | 0.231837 |
| Down | SCN4A | 1 | 0 | 0 | 0.239901 |
| Down | MOGAT1 | 1 | 0 | 0 | 0.193647 |
| Down | UNC13A | 1 | 0 | 0 | 0.171574 |
| Down | KIRREL3 | 1 | 0 | 0 | 0.236645 |

### miRNA-hub gene regulatory network construction

To predict the hub genes for the miRNAs, we used independent online analytical tool (miRNet). The miRNA-hub gene regulatory network contained 2378 nodes, including 2090 miRNAs and 288 hub genes, and 11478 edges (Fig. 5). TUBB2A that was modulated by 206 miRNAs (ex; hsa-mir-8085), BCL2L1 that was modulated by 179 miRNAs (ex; hsa-mir-6735-5p), UBE2D1 that was modulated by 84 miRNAs (ex; hsa-mir-548ap-5p), UBB that was modulated by 81 miRNAs (ex; hsa-mir-132-3p), RPS24 that was modulated by 79 miRNAs (ex; hsa-mir-27a-3p), CAV1 that was modulated by 115 miRNAs (ex; hsa-mir-4514), EPHA2 that was modulated by 90 miRNAs (ex; hsa-mir-3133), ERBB3 that was modulated by 53 miRNAs (ex; hsa-mir-4328), MYH11 that was modulated by 50 miRNAs (ex; hsa-mir-643) and WNT4 that was modulated by 42 miRNAs (ex; hsa-mir-6749-3p) and are listed in Table 5.

**Fig. 5.**
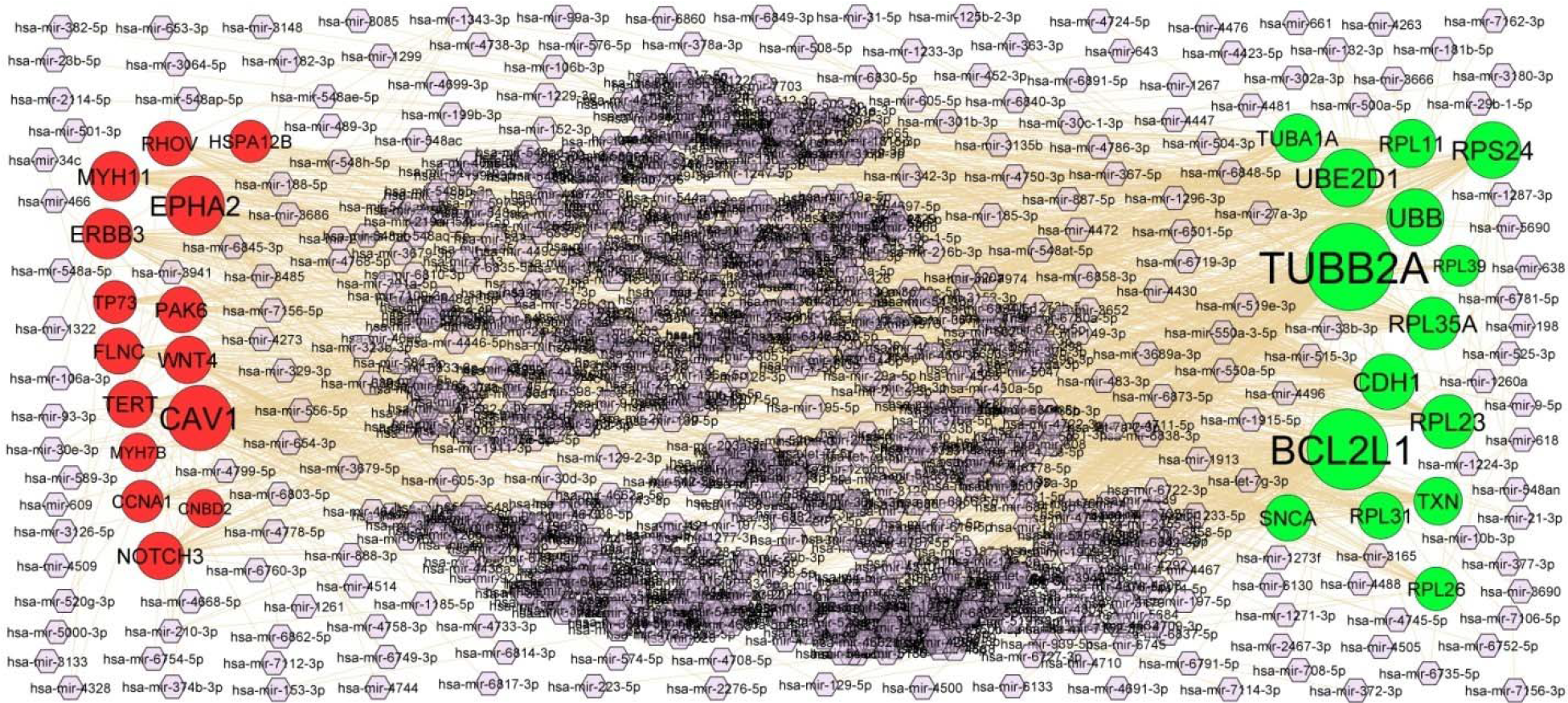
Target gene -miRNA regulatory network between target genes. The purple color diamond nodes represent the key miRNAs; up regulated genes are marked in green; down regulated genes are marked in red.

**Table 5.**
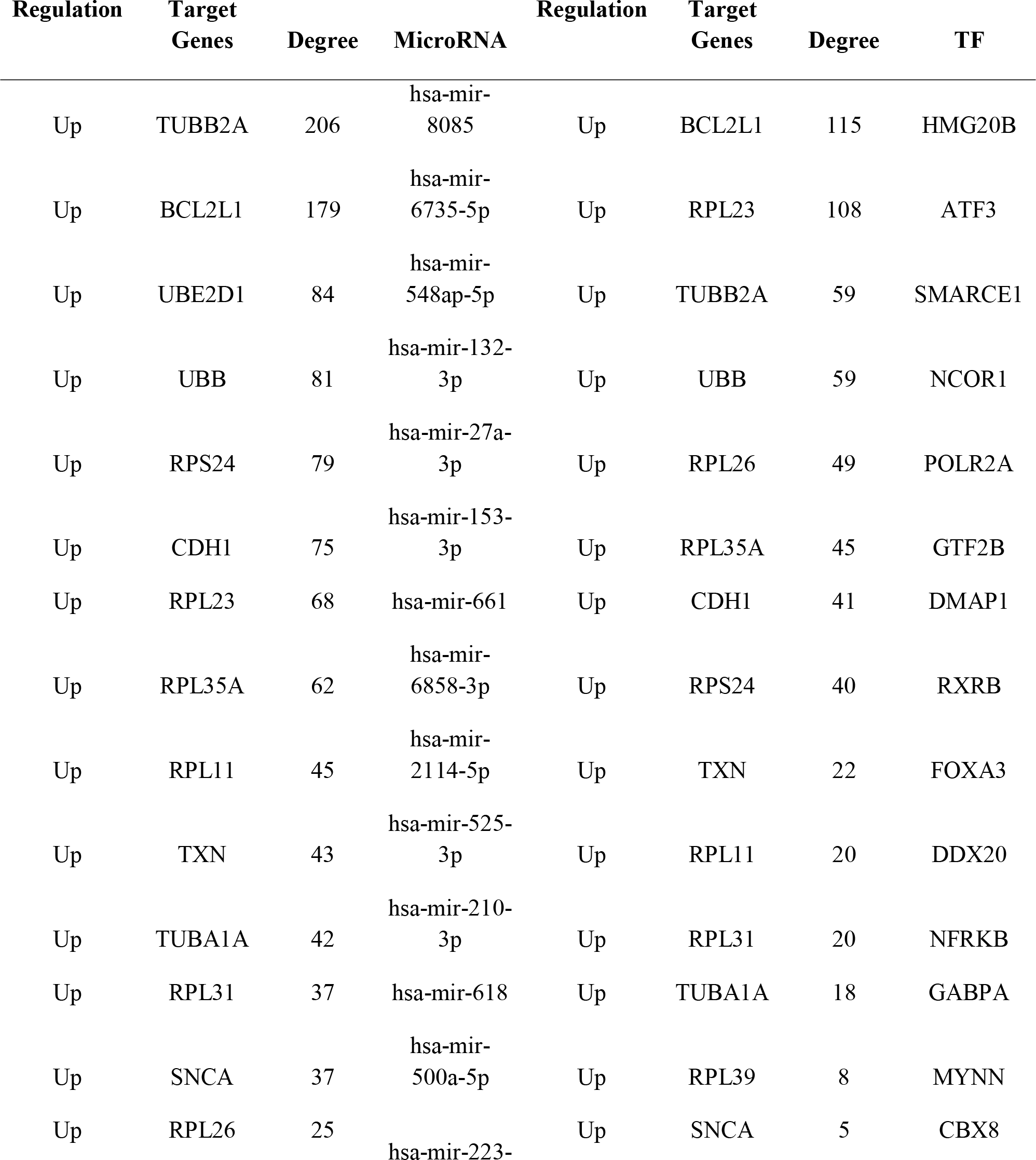
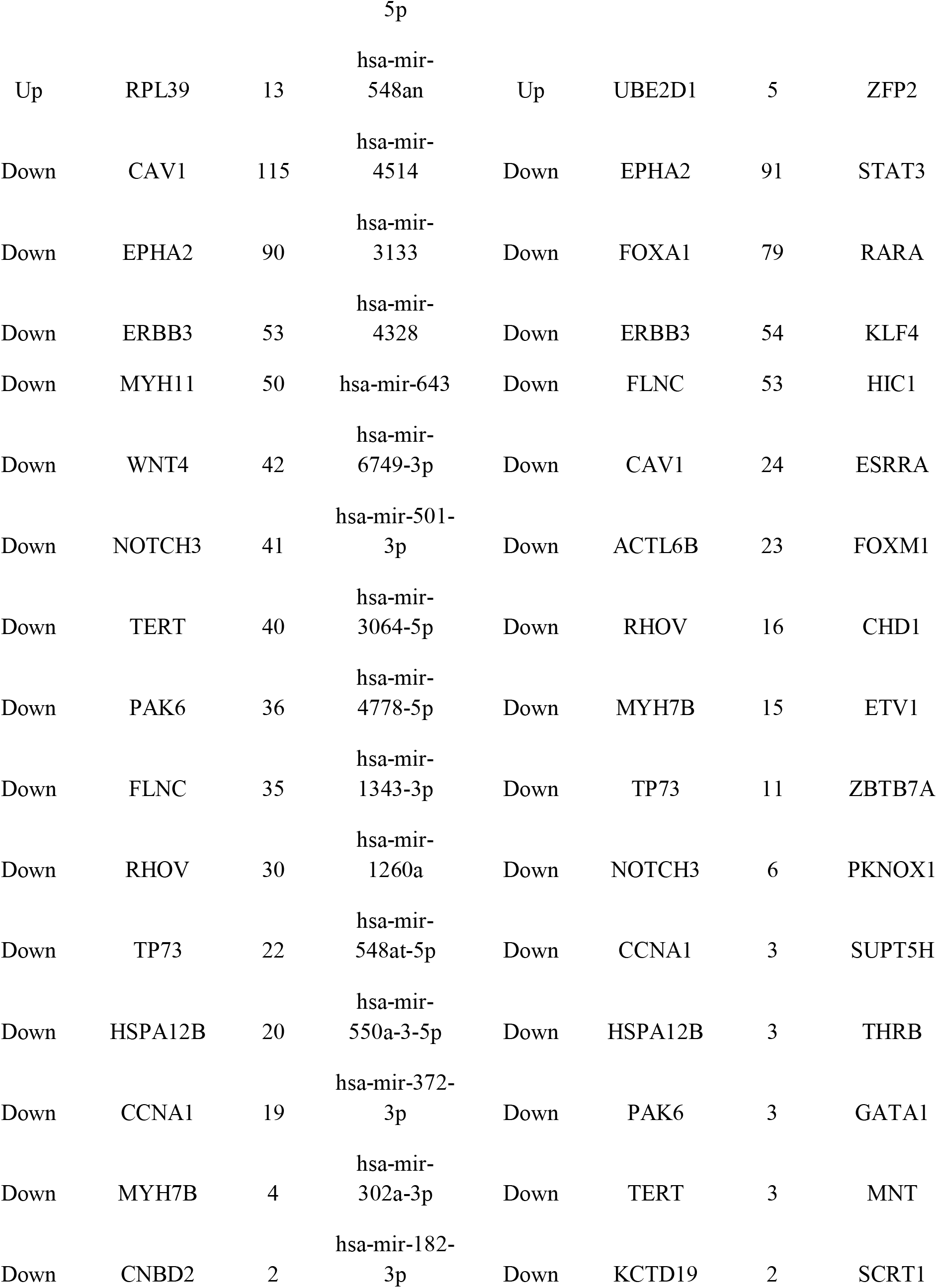
miRNA -target gene and TF -target gene interaction

### TF-hub gene regulatory network construction

To predict the hub genes for the TFs, we used independent online analytical tool (NetworkAnalyst). The TF-hub gene regulatory network contained 582 nodes, including 336 TFs and 246 hub genes, and 6606 edges (Fig. 6). BCL2L1 that was modulated by 115 TFs (ex; HMG20B), RPL23 that was modulated by 108 TFs (ex; ATF3), TUBB2A that was modulated by 59 TFs (ex; SMARCE1), UBB that was modulated by 59 TFs (ex; NCOR1), RPL26 that was modulated by 49 TFs (ex; POLR2A), EPHA2 that was modulated by 91 TFs (ex; STAT3), FOXA1 that was modulated by 79 TFs (ex; RARA), ERBB3 that was modulated by 54 TFs (ex; KLF4), FLNC that was modulated by 53 TFs (ex; HIC1) and CAV1 that was modulated by 24 TFs (ex; ESRRA) and are listed in Table 5.

**Fig. 6.**
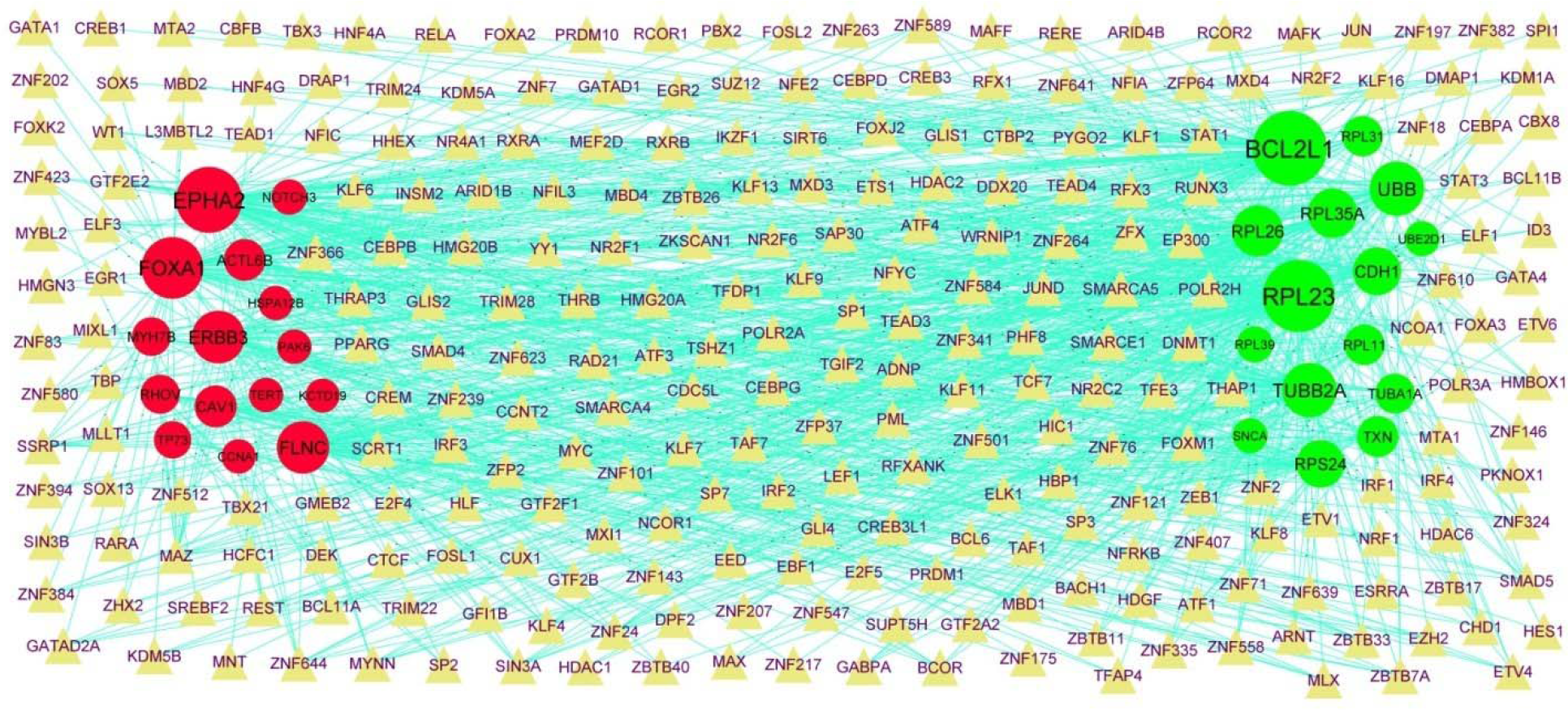
Target gene -TF regulatory network between target genes. The yellow color triangle nodes represent the key TFs; up regulated genes are marked in green; down regulated genes are marked in red.

### Receiver operating characteristic curve (ROC) analysis

ROC curve analyses were performed to verify the hub genes, and area under the curve (AUC) values were calculated. The area under the curve (AUC) values of UBB, UBE2D1, TUBA1A, RPL11, RPS24, NOTCH3, CAV1, CNBD2, CCNA1 and MYH11 were 0.902, 0.907, 0.920, 0.948, 0.940, 0.942, 0.917, 0.922, 0.915 and 0.905, respectively (Fig.7). These result indicated that hub genes could be candidate biomarkers for BD diagnosis.

**Fig. 7.**
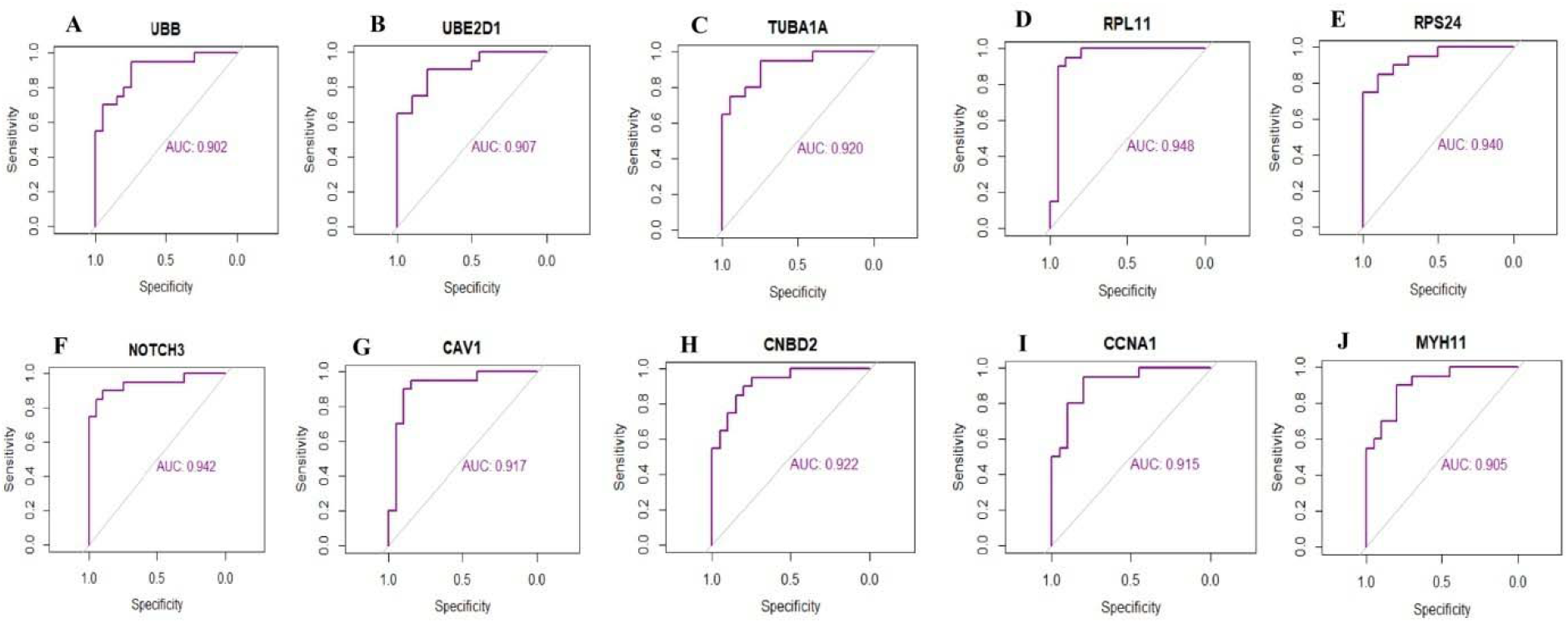
ROC curve analyses of hub genes. A) UBB B) UBE2D1 C) TUBA1A D) RPL11 E) RPS24 F) NOTCH3 G) CAV1 H) CNBD2 I) CCNA1 J) MYH11

## Discussion

The BD is the most common psychiatric disorder worldwide. Although numerous advances have been made in the treatment of BD, the prognosis has remained poor. Therefore, it is crucial to elucidate the molecular mechanism of BD for understanding of the disease progression to develop novel therapeutic targets. Due to the rapid advancement of NGS technology, bioinformatics analysis might contribute to identifying the DEGs and functional pathways involved in the BD.

In this investigation, NGS dataset was selected to identify the DEGs between BD and normal control samples. As a result, 957 DEGs including 477 up regulated and 480 down regulated genes were identified. Wang et al. [44] and Joshi et al. [45] showed altered expression of HBG1 and S100A8 in cardiovascular diseases. Lu et al [46] reported that S100A8 promoted cognitive dysfunction. S100A8 [47] was altered expression in obesity and might serve as a potential prognostic biomarker of obesity. S100A8 [48] and KLHL40 [49] plays an emerging role in pregnancy. The expression of S100A8 [50] is altered in diabetes mellitus.

GO and REACTOME pathway enrichment analysis were conducted to demonstrate interactions of the DEGs. Immune system [51], axon guidance [52], metabolism [53], diseases of metabolism [54] and neuronal system [55] were lined with advancement of BD. Altered expression of SLC6A9 [56] and RHD (Rh blood group D antigen) [57], CHRFAM7A [58], ANXA3 [59], SLC1A5 [60], KCNH2 [61], HP (haptoglobin) [62], SELENBP1 [63], SNCA (synuclein alpha) [64], TGM2 [65], PINK1 [66], B2M [67], QPCT (glutaminyl-peptide cyclotransferase) [68], CBS (cystathionine beta-synthase) [69], NQO2 [70], GLRX5 [71], BASP1 [72], GAS7 [73], GPX1 [74], OLIG2 [75], RPTN (repetin) [76], IL33 [77], SOX10 [78], GRIK1 [79], ZFPM2 [80], SHANK3 [81], ERBB3 [82], ARC (activity regulated cytoskeleton associated protein) [83], GFAP (glial fibrillary acidic protein) [84], PLAT (plasminogen activator, tissue type) [85], GRIK5 [86], CACNB2 [87], NRXN2 [88], TERT (telomerase reverse transcriptase) [89], GNB3 [90], L1CAM [91], EGR3 [92], CAV1 [93], CACNA1B [94], MAGI1 [95], KIR2DL1 [96], PAH (phenylalanine hydroxylase) [97] and CYP3A5 [98] have been shown in schizophrenia. Recent studies showed that SLC6A9 [99], FKBP1B [100], S100A12 [101], SLC6A19 [102], TXN (thioredoxin) [103], TLR9 [104], HP (haptoglobin) [105], ARG1 [106], PINK1 [107], B2M [108], C5AR1 [109], MYADM (myeloid associated differentiation marker) [110], CBS (cystathionine beta-synthase) [111], GPX1 [112], SIAH2 [113], PRDX2 [114], RDH8 [115], CYP11B2 [116], RARRES2 [117], NOX1 [118], IL33 [119], OTC (ornithine transcarbamylase) [120], CYP1A1 [121], NFATC4 [122], TSLP (thymic stromal lymphopoietin) [123], WNT4 [124], MGP (matrix Gla protein) [125], FGFBP1 [126], GHR (growth hormone receptor) [127], ERBB3 [128], GFAP (glial fibrillary acidic protein) [129], CCDC40 [130], CACNB2 [131], CD34 [132], NOTCH3 [133], TERT (telomerase reverse transcriptase) [134], GNB3 [135], TP73 [136], RYR2 [137], ENPEP (glutamyl aminopeptidase) [138], SCN7A [139], WNK4 [140], SFRP5 [141], GDF15 [142], CAV1 [143], KCNA5 [144], FOXC1 [145], ASIC1 [146], VASH2 [147], CXCL8 [148], PAPPA2 [149], KCNMA1 [150], LOX (lysyl oxidase) [151], SPARCL1 [152] and CYP3A5 [153] might have the potential to be used as diagnostic biomarkers of hypertension. Previous studies have reported that RHD (Rh blood group D antigen) [154], S100A12 [155], TXN (thioredoxin) [156], TLR9 [157], S100P [158], TAGLN2 [159], S100A9 [160], CA1 [161], HP (haptoglobin) [162], RPL39 [163], F5 [164], PINK1 [165], B2M [166], S100A11 [167], SLC4A1 [168], CBS (cystathionine beta-synthase) [169], AHSP (alpha hemoglobin stabilizing protein) [170], F12 [171], EPHB4 [172], NFE2 [173], VRK1 [174], GPX1 [175], FOXA1 [176], CYP11B2 [177], NOX1 [178], IL33 [179], NPHS1 [180], OTC (ornithine transcarbamylase) [181], SULF1 [182], CYP1A1 [183], DCN (decorin) [184], ADAMTS7 [185], WNT4 [186], LAMA4 [187], SCN4A [188], CACNB2 [189], GNB3 [190], ENPEP (glutamyl aminopeptidase) [191], WNK4 [192], FOXC1 [193], PAX8 [194], ROBO1 [195], CXCL8 [196], PAPPA2 [197], LOX (lysyl oxidase) [198], NOSTRIN (nitric oxide synthase trafficking) [199], MUC16 [200], MIOX (myo-inositol oxygenase) [201], CYP11A1 [202] and CYP3A5 [203] are involved in the pregnancy complications. Modification in the activity and expression of RHD (Rh blood group D antigen) [204], S100A12 [205], TLR9 [206], ANXA3 [207], S100P [208], TAGLN2 [209], S100A9 [210], KCNH2 [211], HP (haptoglobin) [212], SELENBP1 [213], TANGO2 [214], PINK1 [215], TFR2 [216], B2M [217], LTBP2 [218], PGLYRP1 [219], HRH2 [220], CLEC5A [221], PLSCR4 [222], S100A11 [223], PPBP (pro-platelet basic protein) [224], RAP1GAP [225], CBS (cystathionine beta-synthase) [226], RBM38 [227], MYL4 [228], EPHB4 [229], TNNT1 [230], KBTBD7 [231], GPX1 [232], SIAH2 [113], FOXO4 [233], PRDX2 [234], MAF1 [235], KANK2 [236], SFRP2 [237], BGN (biglycan) [238], HSPB7 [239], ABCG8 [240], CYP11B2 [241], RARRES2 [242], NOX1 [243], CPE (carboxypeptidase E) [244], IL33 [245], OTC (ornithine transcarbamylase) [120], CYP1A1 [246], LRIG3 [247], GJA1 [248], NFATC4 [122], SEMA3F [249], CDH11 [250], DCN (decorin) [251], TSLP (thymic stromal lymphopoietin) [252], ADAMTS7 [253], C1QTNF1 [254], SFTPB (surfactant protein B) [255], WNT4 [256], MGP (matrix Gla protein) [257], SHANK3 [258], ALOX12B [259], MYBPHL (myosin binding protein H like) [260], GHR (growth hormone receptor) [261], ERBB3 [262], PLAT (plasminogen activator, tissue type) [263], ADAMTS2 [264], CACNB2 [265], DAB2IP [266], CD34 [267], COL15A1 [268], MSTN (myostatin) [269], NOTCH3 [270], HSPG2 [271], TERT (telomerase reverse transcriptase) [272], GNB3 [273], MMP15 [274], COL4A2 [275], EGR3 [276], RBM20 [277], RYR2 [278], EPHA2 [279], NDRG4 [280], UNC5B [281], CTNNA3 [282], SFRP5 [283], CUX2 [284], GDF15 [285], AXL (AXL receptor tyrosine kinase) [286], CAV1 [287], KCNA5 [288], FOXC1 [289], RYR1 [290], ROBO1 [291], XIRP2 [292], FZD4 [293], VASH2 [294], CXCL8 [295], KCNMA1 [150], LOX (lysyl oxidase) [296], DNAH11 [297], FMOD (fibromodulin) [298], MFAP4 [299], FLNC (filamin C) [300], LIFR (LIF receptor subunit alpha) [301], MAGI1 [302], SLC39A2 [303] and CYP3A5 [304] are linked to cardiovascular diseases. Studies have also reported that CHRFAM7A [58], ACHE (acetylcholinesterase (Cartwright blood group) [305], HP (haptoglobin) [306], SELENBP1 [63], BASP1 [72], CHRNA1 [307], RPTN (repetin) [76], IL33 [308], TACR1[309], SHANK3 [310], ALOX12B [311], HPN (hepsin) [312], GFAP (glial fibrillary acidic protein) [84], CACNB2 [313], MYT1 [314], NOTCH3 [315], TERT (telomerase reverse transcriptase) [316], GNB3 [317], EGR3 [318], BSN (bassoon presynaptic cytomatrix protein) [319], CUX2 [320], GDF15 [321], CXCL8 [322], MAGI1 [95] and PAH (phenylalanine hydroxylase) [323] are necessary for BD development. S100A12 [324], CDH1 [325], S100A9 [326], ANK1 [327], KCNH2 [328], HP (haptoglobin) [329], FRMD4A [330], SNCA (synuclein alpha) [331], FBXO7 [332], PINK1 [333], B2M [334], KCNH3 [335], CA2 [336], FUZ (fuzzy planar cell polarity protein) [337], UBB (ubiquitin B) [338], C5AR1 [339], CBS (cystathionine beta-synthase) [340], F12 [341], TSPAN5 [342], NQO2 [343], NDUFA1 [344], SRXN1 [345], BASP1 [346], GPX1 [347], OLIG2 [348], EFHC2 [349], FOXA1 [350], PAX4 [351], BGN (biglycan) [352], IL33 [353], LRIG3 [354], GJA1 [355], SHANK3 [356], ARC (activity regulated cytoskeleton associated protein) [357], GFAP (glial fibrillary acidic protein) [358], UNC13A [359], MSTN (myostatin) [360], HSPG2 [361], TERT (telomerase reverse transcriptase) [362], GNB3 [363], L1CAM [364], CALB1 [365], RYR2 [366], MYO15A [367], MYH11 [368], GDF15 [369], CAV1 [370], KIRREL3 [371], COBL (cordon-bleu WH2 repeat protein) [372], LOX (lysyl oxidase) [373], SPARCL1 [374], FLNC (filamin C) [375], TMPRSS4 [376], VWA2 [377], OGDHL (oxoglutarate dehydrogenase L) [378], CYP3A5 [379] and FBXO40 [380] were revealed to be associated with cognitive dysfunction. S100A12 [381], SLC6A19 [382], TXN (thioredoxin) [383], TLR9 [384], S100P [385], S100A9 [386], ANK1 [387], CA1 [388], HP (haptoglobin) [389], ARG1 [390], SNCA (synuclein alpha) [391], STARD10 [392], PINK1 [393], TFR2 [394], B2M [395], PHOSPHO1 [396], CBS (cystathionine beta-synthase) [397], WNT6 [398], NFE2 [399], RNF10 [400], CARM1 [401], GLRX5 [402], PPP2R5B [403], GPX1 [404], FOXO4 [405], ARHGEF12 [406], FOXA1 [407], PAX4 [351], ABCG8 [408], RARRES2 [409], NOX1 [410], CPE (carboxypeptidase E) [411], IL33 [412], ADAMTS7 [413], SFTPB (surfactant protein B) [255], MGP (matrix Gla protein) [257], VTN (vitronectin) [414], PRSS1 [415], GHR (growth hormone receptor) [416], ERBB3 [417], GFAP (glial fibrillary acidic protein) [418], CD34 [419], MSTN (myostatin) [420], NOTCH3 [421], NR5A2 [422], SMOC1 [423], GNB3 [424], CTNNA3 [425], SFRP5 [283], GDF15 [285], AXL (AXL receptor tyrosine kinase) [426], CAV1 [427], LCT (lactase) [428], FZD4 [429], KCNMA1 [430], FMOD (fibromodulin) [431] and CPA6 [432] were revealed to be correlated with disease outcome in patients with diabetes mellitus. Previous studies have demonstrated that SLC6A19 [433], DNAJC6 [434], TXN (thioredoxin) [383], S100A9 [435], HP (haptoglobin) [436], ARG1 [437], SNCA (synuclein alpha) [391], CEP19 [438], CAMP (cathelicidin antimicrobial peptide) [439], PINK1 [440], UBB (ubiquitin B) [441], PHOSPHO1 [396], CBS (cystathionine beta-synthase) [397], PPP2R5B [403], GPX1 [442], PRDX2 [443], LXN (latexin) [444], RGS6 [445], MAF1 [446], HSPB7 [447], PNLIP (pancreatic lipase) [448], NOX1 [410], CPE (carboxypeptidase E) [411], IL33 [449], SP7 [450], ZFPM2 [451], TSLP (thymic stromal lymphopoietin) [123], TACR1 [452], WNT4 [453], MFAP5 [454], GHR (growth hormone receptor) [455], GFAP (glial fibrillary acidic protein) [456], ACTN3 [457], MSTN (myostatin) [420], GNB3 [458], MMP15 [459], KCP (kielin cysteine rich BMP regulator) [460], WNK4 [461], SFRP5 [283], GDF15 [462], CAV1 [427], LCT (lactase) [428], CXCL8 [463], LOX (lysyl oxidase) [464], AIF1L [465] and CYP3A5 [466] are linked with the development mechanisms of obesity. Recent studies have proposed that the DNAJC6 [467], DNAJB2 [468], GPR4 [469] and ZFPM2 [80] are associated with Parkinson Disease. Vauthier et al [434], Cushion et al [470], Barone et al [471], Marcoli et al [472], Qin et al [473], de Nijs et al [474], Bergareche et al [475], Hu et al [476], Liu et al [477], Zhang et al [478], Gorman et al [479], Cherian et al [480], Gururaj et al [481], Belhedi et al [482] and Park et al [483] demonstrated that DNAJC6, TUBB2A, DPM2, SMOX (spermine oxidase), CDYL (chromodomain Y like), EFHC1, SCN4A, DOC2A, ADGRV1, SCAMP5, CACNA1B, KCNT1, KCNT2, CPA6 and CYP3A5 could induce epilepsy. These enriched genes might play essential roles in the advancement of BD and act as novel diagnosis biomarkers or treatment targets of BD.

Construction of PPI network and its modules of DEGs might be helpful for understanding the relationship of developmental BD. Thus, we speculated that UBE2D1, TUBA1A, RPL11, RPS24, CNBD2, CCNA1, RPL31, RPL23, RPL35A and RPL34 might be important hub genes in BD development.

miRNA-hub gene regulatory network and TF-hub gene regulatory network analyses predicted hub genes, miRNAs and TFs. hsa-mir-27a-3p [484], STAT3 [485], KLF4 [486] and ESRRA (estrogen related receptor alpha) [487] were reported to be associated with the prognosis of cognitive dysfunction. Altered expression of ATF3 [488], STAT3 [489] and KLF4 [490] are observed in hypertension, indicating that these biomarkers play a role in hypertension. ATF3 [491], NCOR1 [492], STAT3 [493] and KLF4 [494] have been positively correlated with cardiovascular diseases ATF3 [495], STAT3 [493] and ESRRA (estrogen related receptor alpha) [496] were previously reported to be critical for the development of diabetes mellitus. Ku et al. [497], Su et al. [498], Redonnet et al. [499], Deng et al. [500] and Larsen et al [496] concluded that ATF3, STAT3, retinoic acid receptor, alpha (RARA), KLF4 and ESRRA (estrogen related receptor alpha) were an important participant in obesity. Previous studies have reported that STAT3 [501] and retinoic acid receptor, alpha (RARA) [502] are related to schizophrenia. STAT3 [503] levels are correlated with pregnancy complications. Novel biomarkers include BCL2L1, RPL26, hsa-mir-8085, hsa-mir-6735-5p, hsa-mir-548ap-5p, hsa-mir-132-3p, hsa-mir-4514, hsa-mir-3133, hsa-mir-4328, hsa-mir-6749-3p, HMG20B, SMARCE1, POLR2A and HIC1 might play important roles in the development of BD and act as early diagnosis biomarkers or treatment targets of BD. Therefore, our further study will investigate the interactions of hub genes, miRNA and TFs to shed new light on the molecular mechanisms involved in BD pathology.

In conclusion, the present study provides significant information that might help in understanding the molecular mechanisms of BD. We identified several essential genes that are potentially linked with the development of BD using bioinformatics analyses of DEGs between patients with BD and normal controls. These genes and their pathways will further our understanding of BD etiology, and help improve diagnosis, prevention, and treatment. Our findings suggest that the up regulated expression of UBB, UBE2D1, TUBA1A, RPL11 and RPS24, and the down regulatied of expression of NOTCH3, CAV1, CNBD2, CCNA1 and MYH11 can be considered candidate biomarkers or therapeutic targets for BD. Further studies are needed to confirm our putative finding.

## Acknowledgement

I thank Roel A Ophoff, UCLA, Center for Neurobehavioral Genetics, Los Angeles, CA, USA, very much, the author who deposited their NGS dataset GSE124326, into the public GEO database.

## Ethical approval

This article does not contain any studies with human participants or animals performed by any of the authors.

## Informed consent

No informed consent because this study does not contain human or animals participants.

## Availability of data and materials

The datasets supporting the conclusions of this article are available in the GEO (Gene Expression Omnibus) (https://www.ncbi.nlm.nih.gov/geo/) repository. [(GSE124326) https://www.ncbi.nlm.nih.gov/geo/query/acc.cgi?acc=GSE124326)]

## Consent for publication

Not applicable.

## Author Contributions

B. V. - Writing original draft, and review and editing

C. V. - Software and investigation

